# A Platinum Butterfly Effect: Small Changes Turn an Anticancer Drug into a Non-toxic Metalloantibiotic with In Vivo Efficacy

**DOI:** 10.1101/2025.05.30.657029

**Authors:** Çağrı Özsan, Ann-Britt Schäfer, Abdul Akhir, Obed Akwasi Aning, Sofia Fulgencio, Rahul Maitra, Rupa Rani, Deepanshi Saxena, Fredrik Westerlund, Sidharth Chopra, Michaela Wenzel, Angelo Frei

## Abstract

Widespread resistance to all clinically used antibiotics has sparked investigations into alternative sources for novel and effective antimicrobial agents. Metal-based compounds (metalloantibiotics) have emerged as a promising class of potential antibiotics exhibiting high hit rates against critical bacterial pathogens while not displaying higher toxicity than organic compounds. Here, we describe the exploration of a novel class of non-toxic, Gram-positive acting platinum-based antibacterial agents with micro to nanomolar activity against a range of methicillin and vancomycin-resistant *Staphylococcus aureus* strains. Structure-activity relationship (SAR) studies revealed that modifications of the core scaffold result in reduced antibacterial activity. Mode of action studies investigations showed that lead compound **Pt1** did not impair cell division, RNA, protein, or cell wall synthesis, nor did it affect membrane integrity or potential. Instead, akin to the structurally similar anticancer drug cisplatin (**CisPt**), **Pt1** treatment resulted in reduced DNA staining, visible nucleoid compaction, and activation of DNA damage repair responses. Importantly, we could show that **Pt1** is able to interact with and damage DNA directly, resulting in DNA strand breaks and fragmentation. **Pt1** activity can be reduced significantly by high amounts of a hydroxyl radical scavenger. Derivative **Pt8**, which retained DNA-damaging activity but was less potent in terms of antibacterial activity, was not affected by the presence of radical scavengers, suggesting that **Pt1** possesses a multimodal mechanism. In line with this observation, no resistance development to **Pt1** was observed over the course of 36 passages. Finally, we could demonstrate the *in vivo* activity of **Pt1**, which significantly reduced the bacterial load in a murine *S. aureus* skin infection model. Altogether, these findings shed light on the SAR and antibacterial mode of action of a novel class of platinum metalloantibiotics, validate its *in vivo* efficacy, and pave the way for further exploration of platinum compounds as novel drug candidates with a highly attractive activity profile.

## Introduction

Antimicrobial resistance (AMR) has emerged as one of the most significant health problems of the 21st century.^1^ In 2019, 4.95 million deaths were associated with bacterial AMR, and it was the direct cause of 1.27 million of those. It is forecasted to be the leading cause of mortality by 2050, with approximately 10 million annual deaths.^2^ At the same time, the conventional pipeline for antibacterial drug development, relying on organic medicinal chemistry, shows insufficient development to avert this scenario. Between 2013 and 2022, only 19 new small-molecule drugs have been approved for antibacterial applications. None of these 19 introduced a first-in-class mechanism, with the most recent being the tuberculosis drug bedaquiline in 2012.^3^ This stagnation emphasizes a need to explore novel chemical spaces and unconventional approaches in antibacterial drug discovery.

Most antibiotics do not conform to the traditionally established medicinal chemistry structural criteria such as Lipinski’s Rule of Five.^3,4^ It is hence highly likely that entire classes of antibiotics remain unexplored due to having been excluded from screening campaigns based on inadequate criteria. To investigate previously unexplored chemical spaces, the Community for Open Antimicrobial Drug Discovery (CO-ADD) was launched in 2015. Compounds submitted to CO-ADD are evaluated against key bacterial and fungal pathogens to identify novel antibacterial scaffolds.^5^ A considerable amount of the submitted compounds do not meet the ‘usual drug-likeness’ properties including a notable class: metal complexes. Among 906 submitted metal complexes, representing 29 different metals, 246 demonstrated activity against bacterial strains, a remarkable hit rate of 27%, especially considering that purely organic molecules only showed a hit rate of 1.6%. Moreover, 9.9% of the active metal complexes exhibited antibacterial activity without significant cytotoxicity to human cells, over ten times the rate observed in the general CO-ADD library (0.87%).^6^

Metal complexes could be the key to the ‘escape from flatland’ of purely organic molecules with their highly three-dimensional structures offering greater structural diversity.^7,8^ Morrison *et al*. demonstrated that a library of 71 selected ‘metallofragments’ exhibits significantly greater three-dimensionality compared to a library of over 18,000 organic small molecules.^9^ As molecular interactions with biological systems are fundamentally structure-dependent, molecular shape represents a critical factor in determining biological activity. Consequently, broad structural diversity is a highly desirable property to maximize the likelihood of identifying hits against diverse biological targets.^9,10^ In addition to the geometrical diversity, metal complexes can offer unique modes of action such as ligand exchange or release, redox properties, catalytic characteristics to form toxic species, and exchange of metals that stabilize the native structures of proteins causing malfunction and denaturation.^6,8,11^

The exploration of inorganic or metal-based compounds as medicines is not a novel area of research. Indeed, many inorganic compounds are used in the clinics today and many more are under clinical investigation.^12^ The golden age of inorganic medicinal chemistry started with the approval of the first platinum-based chemotherapy agent, cisplatin (**CisPt**), in 1978. Today, platinum-drugs are still used as a gold-standard first-line chemotherapy against many cancer types.^13,14^ In the aforementioned CO-ADD study, platinum complexes represented the highest number of active metal-based compounds without cytotoxicity.^6^ Interestingly, the antibacterial potential of platinum complexes had already been noticed even before **CisPt**, discovered by Rosenberg and colleagues as an unexpected result of an electrochemical experiment with *Escherichia coli.*^15,16^ Until Lippard and co-workers reported monofunctional platinum complexes as inducers of bacterial filamentation and lysis in lysogenic *E. coli* in 2014, there had been no investigation of platinum complexes as antibacterial agents as the focus shifted to cancer research.^17^ In recent years, there have been more studies on the antimicrobial properties of platinum complexes.^18–24^

In 2021, Frei *et. al* established 1,5-cyclooctadiene (COD) platinum complexes (PtCOD) as promising antibacterials against Gram-positive bacteria. Structure-activity relationship (SAR) studies of this compound class revealed that PtCOD complexes bearing two halogen ligands (**Pt1**, **Figure 1**) maintain promising activity against a broad panel of Gram-positive bacteria. Additionally, *in vivo* toxicity was evaluated using larvae of the greater wax moth *Galleria mellonella*, and no toxicity was observed up to 0.4 mM (6-7.5 mg/kg).^25^

**Figure 1.**
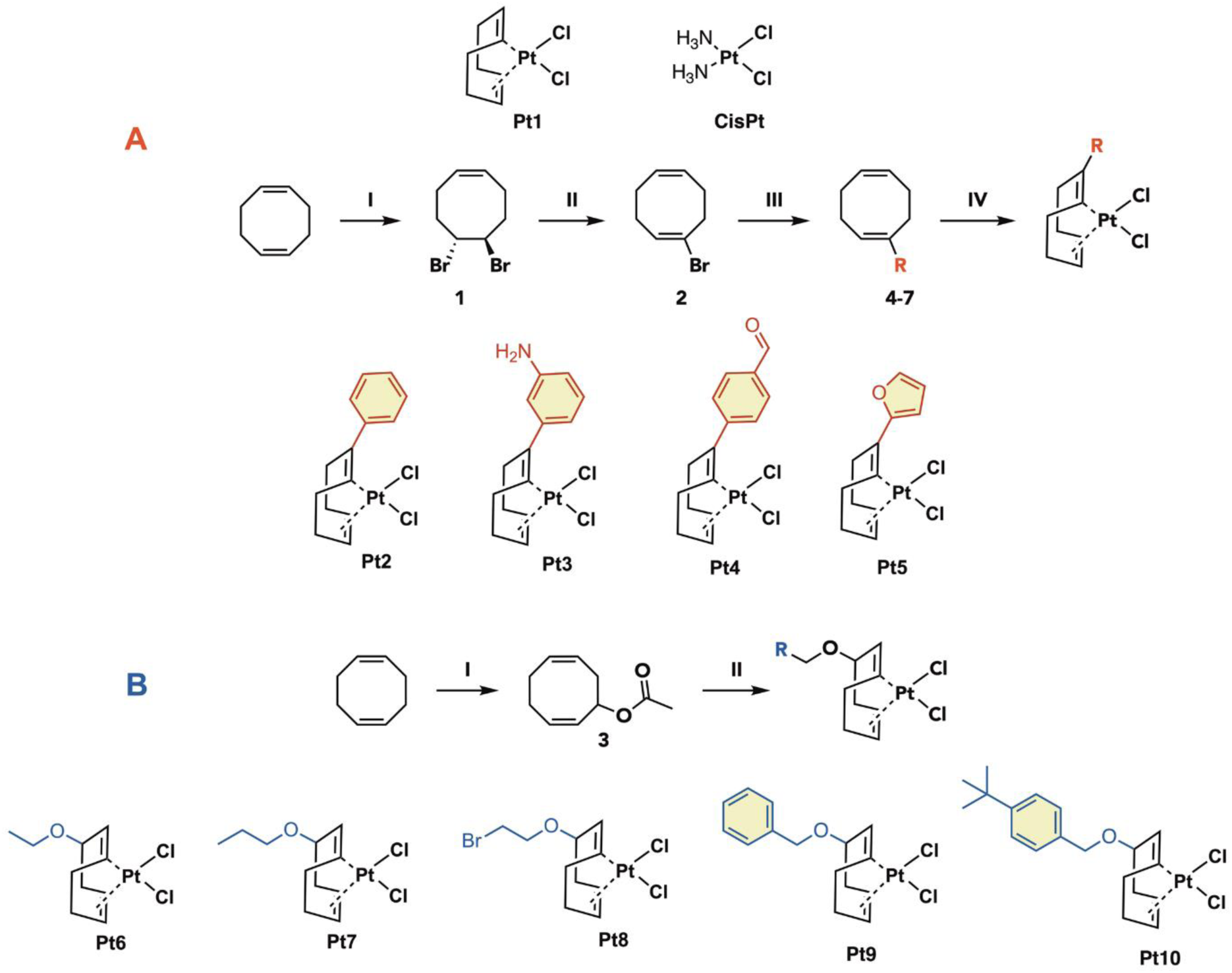
Reaction scheme for double bond-modified COD ligands and corresponding Pt(II) complexes. **(A) I:** Br2, DCM, 30 mins, r.t., 77%; **II:** DBU, DMF, overnight, 80°C, 21%; **III:** RB(OH)2, Pd(PPh3)4, Na2CO3, Dioxane:H2O, 4 h, reflux, 28-41%; **IV:**K2PtCl4, EtOH:H2O, 1 h, MW, 90°C, 12-55%. (**B**) Reaction scheme for allylic position modified Pt(II) complexes. **I:** *t*-butylhydroperoxide, CuCl, AcOH, 52 h, reflux, 7%; **II:** K2PtCl4, H2O:RCH2OH, 1 h, MW, 90°C, 7-13%.

Overall, PtCOD complexes have shown promising antibacterial activity with no *in vitro* or *in vivo* toxicity. To further explore the potential of PtCOD compounds as metalloantibiotics, we embarked on an in-depth exploration of their structure-activity and antibacterial mechanisms. Herein, we report a detailed study of the biological profile of this compound class. By preparing a series of new derivatives and conducting a range of mechanistic studies we elucidate the mechanism of action of **Pt1**, evaluate its propensity to induce resistance, and assess its *in vivo* efficacy in a murine skin infection model.

## Results

### Synthesis of Pt(COD-R)Cl2 Complexes

In previous work^25^, PtCOD complexes bearing halogen ligands have been investigated and found as promising antibacterial agents against Gram-positive bacteria, **Pt1** becoming a new lead compound.^25^ To further elucidate their SAR, novel derivatives of this compound class were synthesized. Given the observed significance of the halogen ligands in conferring antibacterial activity, we aimed to explore modifications on the COD ring. The modifications were performed either at the double-bond **(Figure 2A, Pt2-5**) or allylic position (**Figure 2B, Pt6-Pt10**) starting from commercially available COD. The modified ligands were reacted with K2PtCl4 to obtain the final complexes. All final complexes were purified by silica chromatography and characterised by NMR and Mass Spectrometry (cf. Supporting Information).

**Figure 2.**
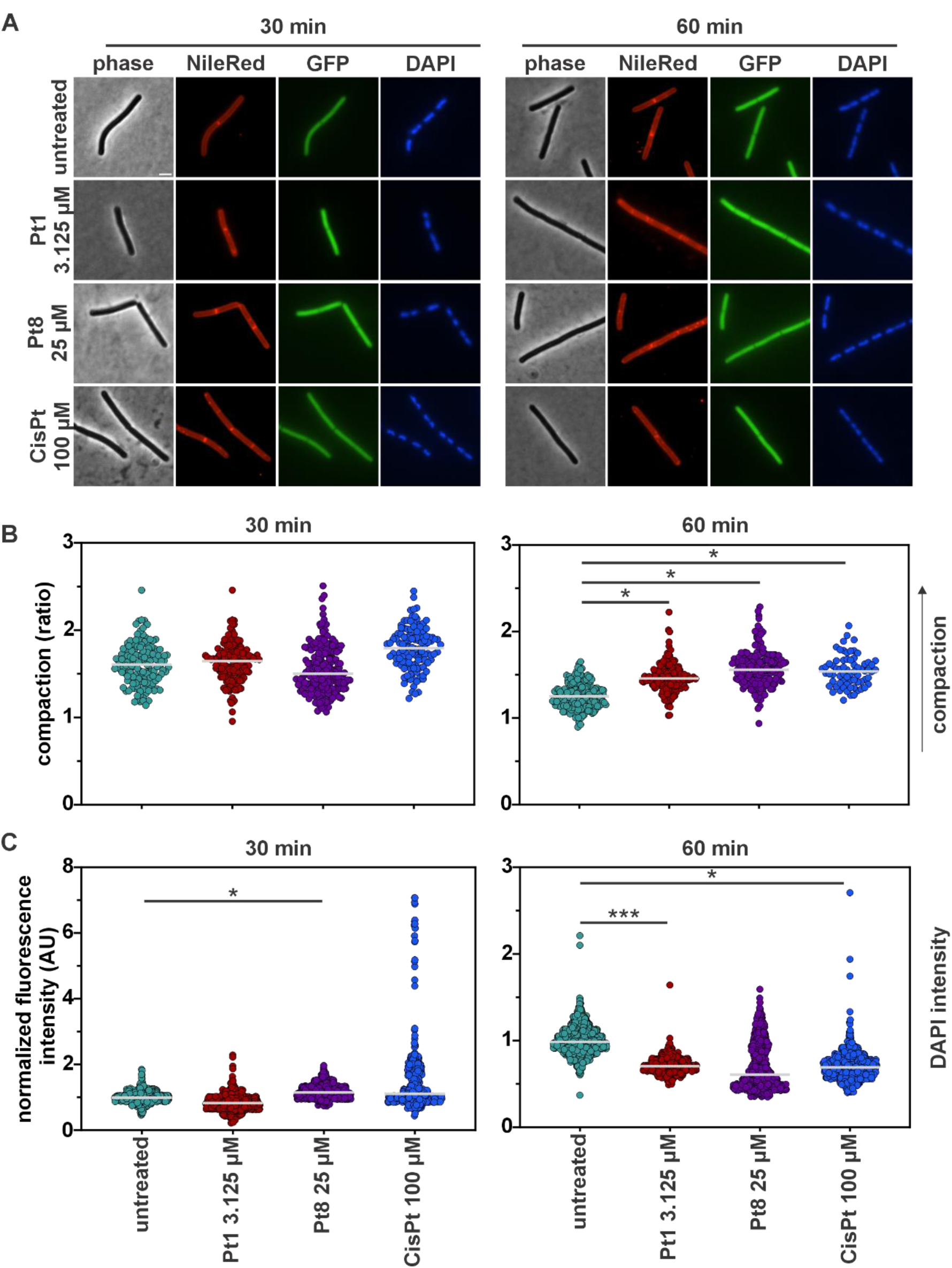
Bacterial cytological profiling of *B. subtilis* bSS82, constitutively expressing GFP from the *PrpsD* promoter, stained with Nile red (membrane) and DAPI (DNA). Cultures were treated with **Pt1** (3.125 µM), **Pt8** (25 µM), and **CisPt** (100 µM) for 30 and 60 min, respectively. (**A**) Phase contrast and fluorescence microscopy images. Scale bar 2 µm. (**B-C**) Quantification of nucleoid compaction measured as a ratio of the whole cell area vs the area occupied by the DAPI-stained nucleoid (**B**) and DAPI fluorescence intensity (**C**). Grey lines indicate the median. Graphs show pooled data from three biological replicates. See **Figure S6** for cell populations divided by individual replicates. Significance was tested with nested t-tests (p values: *≤0.5, **≤0.01, ***≤0.001).

### Antibacterial activity of Pt(COD-R)Cl2 complexes

To assess the antibacterial activity of the complexes, minimum inhibitory concentration (MIC) assays were conducted against representative Gram-positive bacteria, namely methicillin-sensitive *Staphylococcus aureus* (MSSA), methicillin-resistant *S. aureus* (MRSA), and *Bacillus subtilis* (**Table 1A**). Substituting the double bond with benzyl or furan rings (**Pt2- 5**) significantly reduced activity against Gram-positive strains. While **Pt2** showed limited activity (MIC 50–100 µM), the other complexes were inactive in this concentration range. In contrast, modifications at the allylic position demonstrated good antibacterial activity (**Pt8**), yet did not exceed the activity of the parent compound **Pt1**.

**Table 1.**
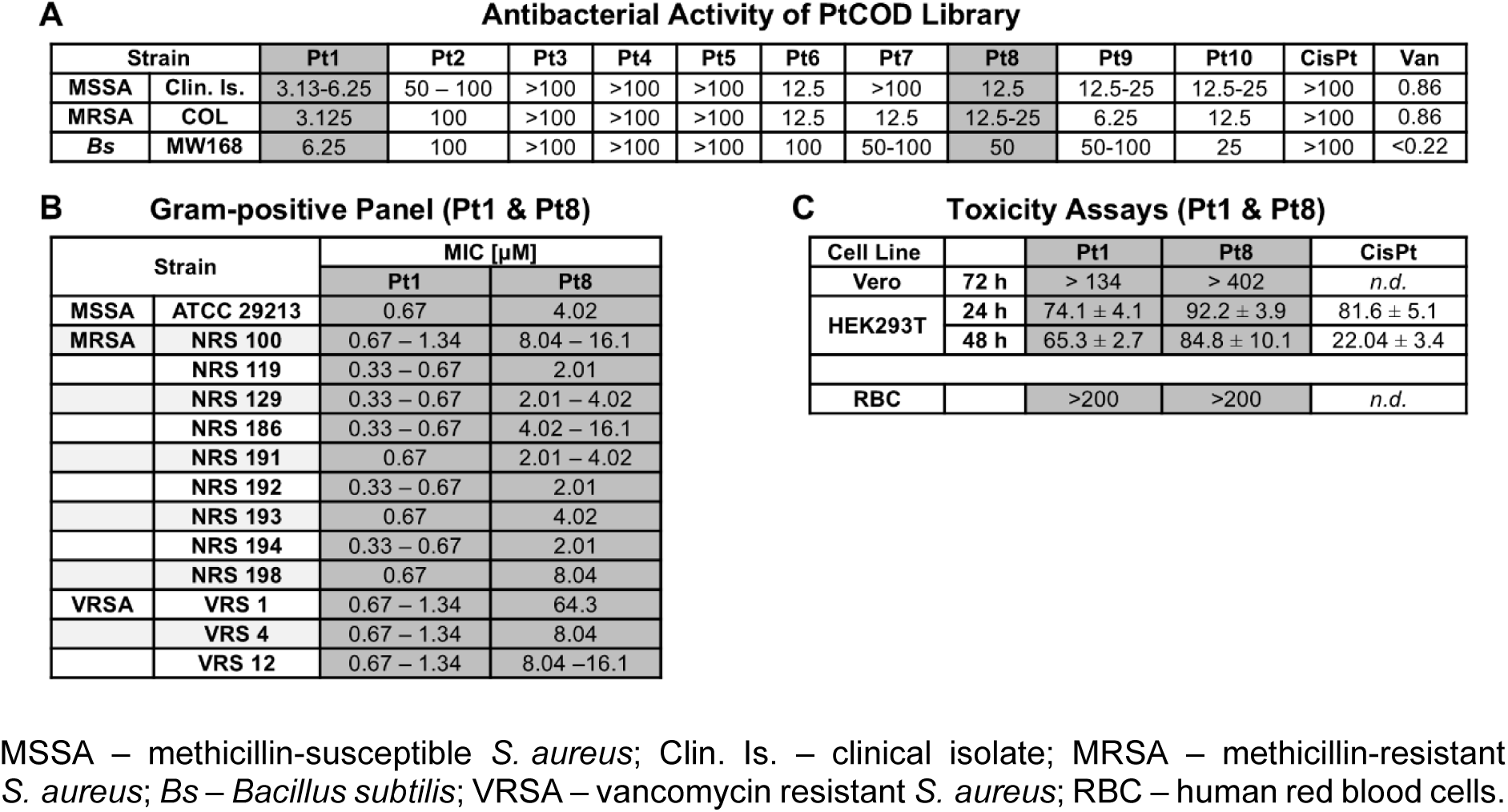
Antibacterial activity and cytotoxicity. (**A**) MICs (µM) of synthesized PtCOD complexes against a selection of Gram-positive strains. (**B**) MICs (µM) of lead compounds **Pt1** and **Pt8** against an extended panel of drug-resistant *S. aureus* strains. (**C**) IC50 and HC50 values (µM) of **Pt1** and **Pt8** against mammalian cell lines and human red blood cells.

None of the complexes showed antibacterial activity against *E. coli* (**Table S1**). The structural similarity between the platinum-based anticancer drug **CisPt** and our lead compounds led us to examine the antibacterial effects of the anticancer drugs **CisPt** and oxaliplatin against Gram-positive bacteria, yet neither showed activity up to 100 µM (**Table 1A** and **S2**). The antibacterial activity of **CisPt** has been investigated several times since the first report by Rosenberg *et al.* However generally only moderate levels of activity were observed and generally this was lower against Gram-positive strains compared to Gram-negative strains.^26,27^ To confirm whether the observed antibacterial activity originates from the complexes themselves, rather than the ligand (COD) or platinum alone, the antibacterial effects of 1,5- cyclooctadiene and K₂PtCl₄ against *S. aureus* and *E. coli* were examined. Neither of them was found to be active up to 200 µM (**Table S3)**.

The parent compound **Pt1** as well as **Pt8,** which was the most promising amongst the newly synthesized allylic-modified complexes, were selected as lead compounds for further characterization. To this end, their antibacterial activity against an extended Gram-positive panel including a range of MRSA and vancomycin-resistant *S. aureus* (VRSA) strains was assessed (**Table 1B,** see **Table S4** for corresponding values in µg/mL). **Pt1** showed excellent activity against the whole panel, while **Pt8** was slightly less active. Both lead compounds were also tested against an extended Gram-negative panel consisting of *E. coli*, *Klebsiella pneumoniae*, *Acinetobacter baumannii*, and *Pseudomonas aeruginosa.* However, neither compound was active up to the highest tested concentrations (**Table S4)**.

### Haemolytic and cytotoxic activity of Pt(COD-R)Cl2 complexes

Selectivity of antibacterial compounds for bacteria over mammalian cells is essential. Therefore, 50% haemolytic concentrations (HC50) were determined against human red blood cells (RBC). All tested PtCOD complexes showed HC50 values above 200 µM and were thus classified as not haemolytic (**Table 1C** and **S5**).

To test cytotoxicity of the lead compounds against mammalian cell lines, 50% inhibitory concentrations (IC50) were determined against African Green Monkey kidney cells (Vero) and Human embryonic kidney cells (HEK293T). Neither **Pt1** nor **Pt8** significantly reduced the viability of Vero cells up to the highest tested concentrations (134 µM and 402 µM respectively, **Table 1C** and **S6**, **Figure S1-2**). IC50 values of **Pt1** (74.1 ± 4.1 µM) and **Pt8** (92.2 ± 3.9 µM) against HEK293T cells after 24 h were comparable to those of **CisPt** (81.6 ± 5.1 µM). However, neither **Pt1** (65.3 ± 2.7 µM) nor **Pt8** (84.8 ±10.1 µM) showed a large increase of toxicity after 48 h, which is in stark contrast to **CisPt** (22.04 ± 3.4 µM, **Table 1C**, **Figure S3**).

Taken together, **Pt1** and **Pt8** show promising antibacterial activity against a broad panel of Gram-positive pathogens but do not cause haemolysis and exhibit only minor toxicity against mammalian cell lines.

### Mode of action studies of Pt1 and Pt8

Encouraged by these results, we proceeded to study the modes of action of **Pt1** and **Pt8** in comparison with **CisPt** using phenotypic analysis of the Gram-positive model organism *B. subtilis*. To this end, we first confirmed the MICs against *B. subtilis* 168CA, the background strain of our reporter constructs (**Pt1**: 6.25 µM; **Pt8**: 50 µM). Based on these values, we determined the optimal stressor concentrations (OSCs) in acute shock experiments. The optimal stressor concentration is a concentration that leads to a 50-70% reduction of bacterial growth during mid-exponential phase. It imposes sufficient stress to elicit a clear phenotype without causing excessive cell death or lysis, which would mask the direct effects of the test compound.^29^ We determined OSCs of 3.125 µM for **Pt1** and 25 µM for **Pt8** (**Figure S4**). No growth inhibition could be observed with **CisPt** (**Figure S4**), leading to the decision to proceed with the highest tested concentration (100 µM). Both **Pt1** and **Pt8** showed a reproducible delay until the onset of growth inhibition (∼30 min for **Pt1** and ∼60 min for **Pt8**, **Figure S4**). Hence, we chose 30 and 60 min after antibiotic addition as timepoints for phenotypic analysis experiments.

### Bacterial cytological profiling

To get an overview of the cellular effects of the compounds, we employed bacterial cytological profiling (BCP). To this end, we used a strain intracellularly expressing green-fluorescent protein (GFP) from the strong, constitutive *PrpsD* promoter (*B. subtilis* bSS82^29^), stained with the membrane dye Nile red and the DNA dye DAPI. Combined with phase contrast microscopy, this setup allows simultaneous assessment of cell morphology, cell lysis, pore formation, membrane morphology, and nucleoid morphology in one assay.^28^

None of the compounds caused apparent differences in overall cell morphology (phase contrast) or compromised membrane integrity (GFP) (**Figure 2A**). Nile red foci, indicating membrane stress, were observed in **Pt1**-treated samples. However, quantification of the percentage of cells exhibiting Nile red foci could not statistically support this observation due to high variability among treated samples (**Figure S5**). These results suggest that membrane damage may be a secondary component of the **Pt1** mode of action that may occur to varying degrees, but it is unlikely to be the primary mechanism of action.

Striking effects of **Pt1** and **Pt8** were observed on the bacterial nucleoid (DAPI), showing visible changes in both nucleoid morphology and fluorescence intensity (**Figure 2A**). When we quantified these differences, both **Pt1** and **Pt8** as well as **CisPt** showed clear nucleoid compaction after 60 min of treatment (**Figure 2B**). This is indicative of DNA packing defects. Similarly, **Pt1** and **CisPt** caused a significant reduction of the DAPI signal in the same timeframe, **Pt8** showing a similar trend. This phenotype points towards compromised DNA integrity and can indicate DNA fragmentation or other structural changes that decrease the availability of the minor groove to the DAPI dye (**Figure 2C**). These results gave rise to the hypothesis that **Pt1** and **Pt8** affect the structural integrity of DNA, and consequently nucleoid packing, possibly in a manner similar to **CisPt**.

### Cell envelope stress

Nile red stains suggested that **Pt1** could partially act through membrane activity. We followed up on this lead to explore whether it may allow molecules smaller than GFP to pass the cell membrane.^28^ We first tested the uptake of the membrane-impermeable, red-fluorescent dye propidium iodide (PI). Neither **Pt1** nor **Pt8** showed increased PI uptake confirming that they do not compromise membrane integrity (**Figure 3A**). Likewise, no membrane depolarization could be observed when measured with the membrane potentiometric fluorescent dye 3,3’- dipropylthiadicarbocyanine iodide (DiSC(3)5)^30^ (**Figure 3B**), showing that no relevant ion translocation is caused by the compounds. This finding was independently confirmed by assessing the localization of the cell division regulation protein MinD, which localizes at midcell and cell poles in healthy cells, but delocalizes into membrane-associated clusters upon loss of membrane potential^31^ (**Figure 3C**). These results show that neither **Pt1** nor **Pt8** affect membrane integrity.

**Figure 3.**
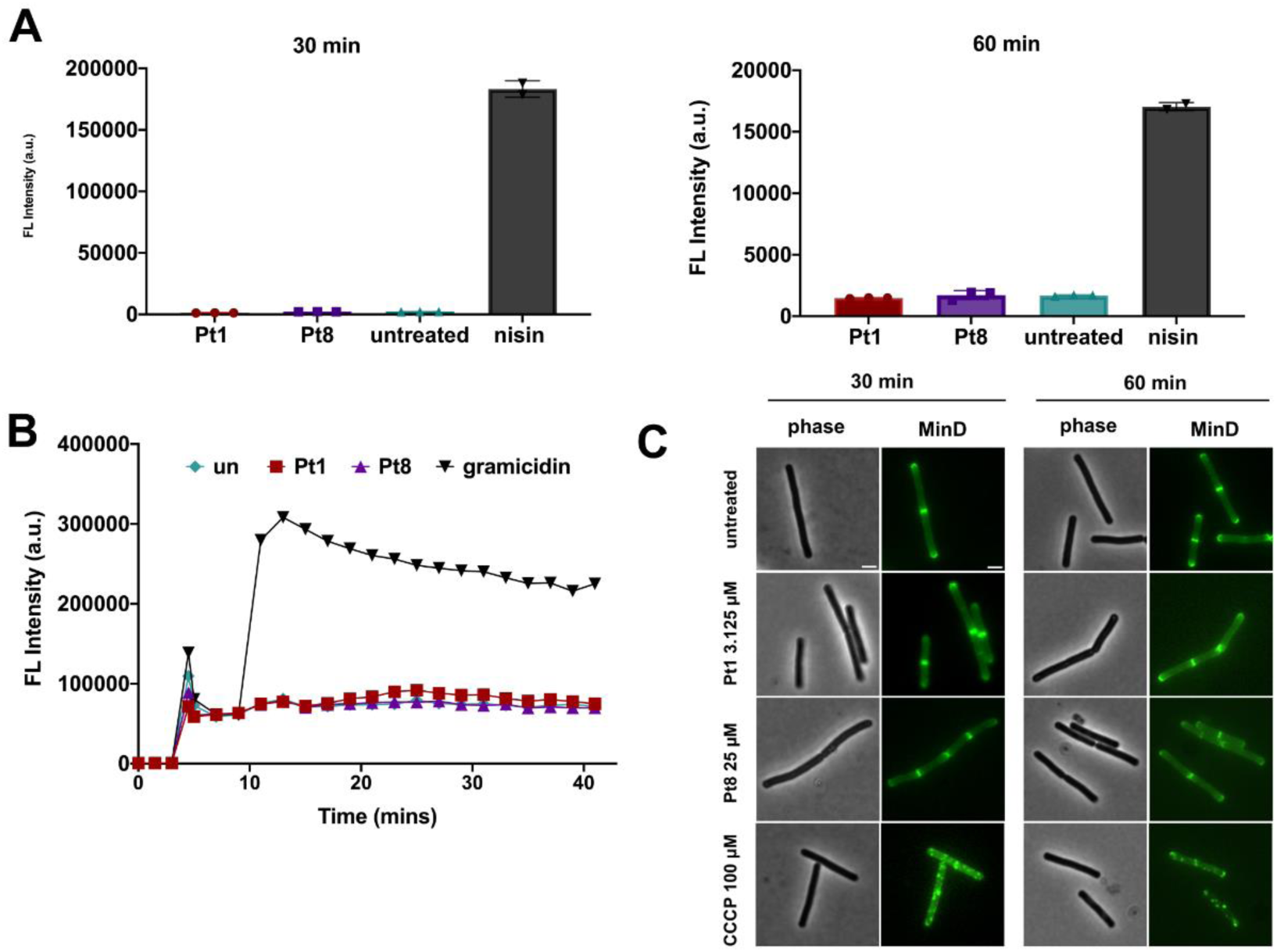
Effects on membrane integrity. (**A**) Propidium Iodide staining of *B. subtilis* 168CA (pore formation assay). 0.23 µM nisin was used as positive control. Note the different scales at 30 and 60 min. (**B**) DiSC3(5)) assay in *B. subtilis* 168CA (membrane depolarization assay). 0.53 µM gramicidin was used as positive control. (**C**) Localization of MinD-GFP in *B. subtilis* TB35 (*Pxyl-gfp-minD*). Localization of MinD depends on the membrane potential. 100 µM CCCP were used as positive control. All assays were performed after treatment with 3.125 µM **Pt1** and 25 µM **Pt8**. Scale bar 2 µm.

Despite not being statistically significant, the tendency of **Pt1** and **Pt8** to cause membrane foci was apparent (**Figure S5**). In addition to direct membrane damage, an impairment of cell wall synthesis or cell division can also cause Nile red foci.^28,32^ To assess possible effects on these processes, we examined the localization of the proteins MreB and FtsZ. MreB is an actin homologue that moves around the lateral cell axis driving the cell wall elongation machinery forward. This movement stops when cell wall synthesis is impaired, making MreB mobility a reliable reporter for cell wall synthesis inhibition.^28,33,34^ However, neither **Pt1** nor **Pt8** affected MreB mobility, showing that they do not inhibit this process (**Figure S7**). Likewise, the localization of the tubulin homologue FtsZ, the major cell division protein in bacteria^35^, remained unaffected demonstrating that the division machinery is not disturbed (**Figure S8**). These results suggest that **Pt1** and **Pt8** do not elicit their antibacterial activity through impairment of cell envelope function.

### Effects on transcription and translation

Strong GFP signal in the BCP suggested that the platinum compounds did not interfere with protein expression. However, *PrpsD* is a strong, constitutive promoter and mild effects will not be visible against the background of already accumulated GFP. To confirm that **Pt1** and **Pt8** do not affect the transcription and translation machineries, we assessed the localization of the RNA polymerase β’ subunit RpoC and the ribosomal protein RpsB, which have been used as reporters for transcription and translation inhibition, respectively.^36^ Neither protein showed a distinct phenotype after treatment with **Pt1** and **Pt8**, confirming our notion (**Figure S9-10**).

We then directly tested their capacity to impair protein production using inducible GFP-MinD as readout. To this end, we added the respective compounds prior to induction of protein expression with xylose. No differences in GFP-MinD expression were observed, demonstrating that protein expression is unaffected by **Pt1** and **Pt8** (**Figure S11**).

### DNA Damage

BCP revealed that **Pt1** and **Pt8** elicited their strongest effect on the bacterial nucleoid, both in terms of nucleoid packing and fluorescence intensity of the DAPI stain (**Figure 2**). A reduced DAPI signal indicates less efficient DNA binding of the dye, suggesting inaccessibility of the minor groove^37^ and thus structural changes to the DNA. To confirm whether the compounds induce DNA damage in bacterial cells, the localization of RecA was assessed. RecA is part of the bacterial SOS response to DNA damage. It is a DNA-associated protein that recognizes single-stranded DNA, which occurs as a consequence of strand breaks. If strand breaks are present, RecA accumulates at the damaged sites, appearing as foci or elongated structures.^38^ Since the RecA response is strongest shortly after stress induction^39^, we additionally examined a 10 min timepoint in this assay. Indeed, clear RecA accumulations were found on the nucleoids of cells treated with **Pt1**, **Pt8**, and **CisPt**, with the strongest response induced after 10 min and **Pt1** showing the most pronounced effects (**Figure 4**).

**Figure 4.**
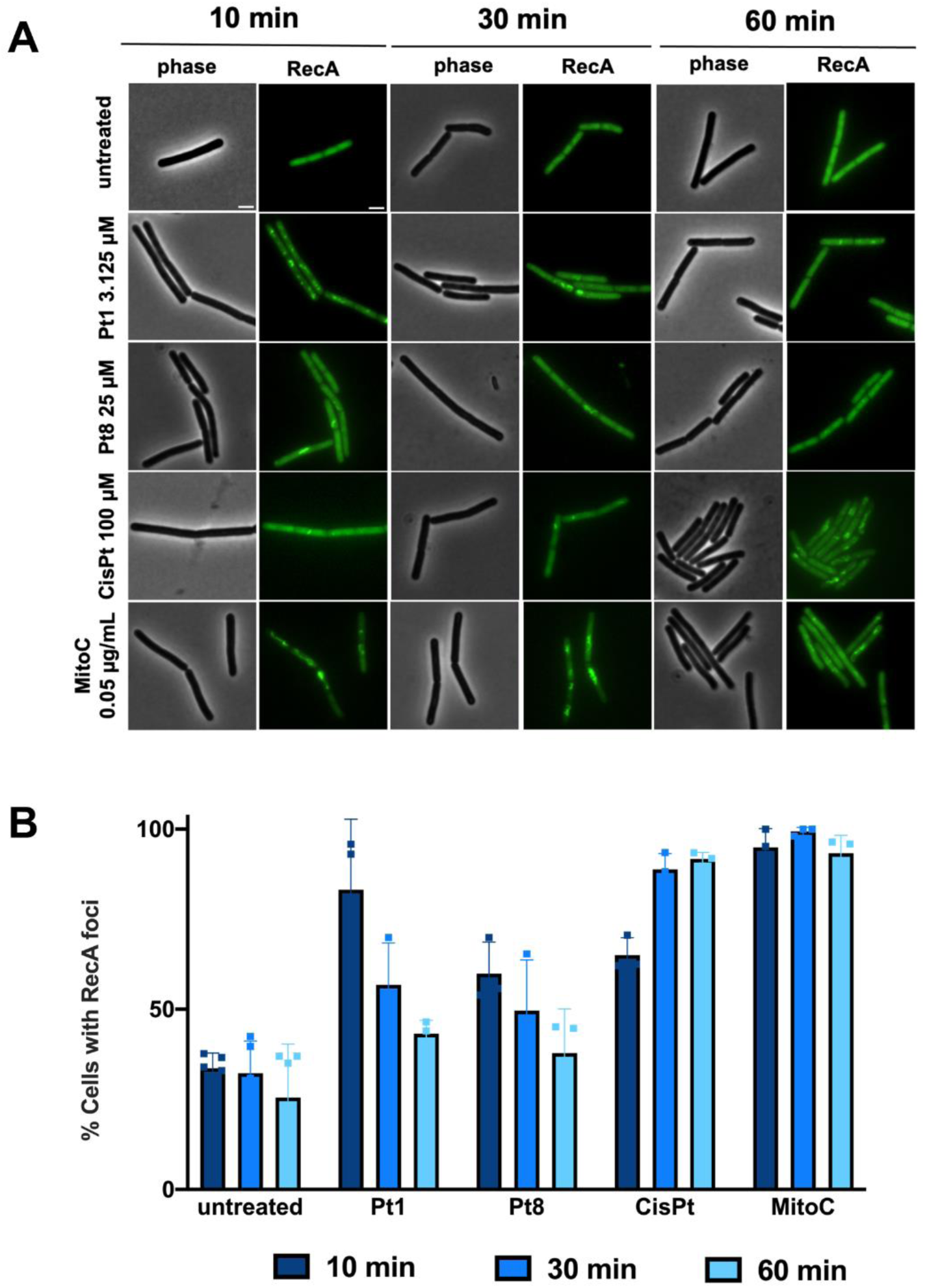
Localization of RecA-GFP (*B. subtilis* UG10) after treatment with **Pt1** (3.125 µM), **Pt8** (25 µM), and **CisPt** (100 µM). Mitomycin C (**MitoC**, 0.05 µg/mL) was used as positive control. (**A**) Phase contrast and fluorescence microscopy images. Scale bar 2 µm. (**B**) Quantification of microscopy images. Cells with distinct RecA foci were counted and expressed relative to the number of total cells. A minimum of 50 cells were counted per condition per replicate. Error bars show standard deviation of at least three biological replicates.

We then wondered whether the compounds directly interact with and damage DNA. To address this, commercially available λ-phage DNA (ʎ-DNA) was incubated with **Pt1**, **Pt8**, and **CisPt**, followed by staining with the DNA dye YOYO-1 and stretching of DNA strands on functionalized glass slides. Samples were then subjected to single-molecule fluorescence imaging followed by length analysis of individual DNA strands.^40^ We observed a clear shift of the DNA molecule length, revealing shorter fragments in samples treated with 3.125 µM **Pt1**, 25 µM **Pt8**, and 100 µM **CisPt** (**Figure 5**). These findings demonstrate that the compounds do indeed directly interact with and damage DNA. The data further shows that both **Pt1** and **Pt8** induce strand breaks, which was expected based on the RecA recruitment results (**Figure 4**). These observations match the known mechanism of **CisPt**, which causes 1,2-intrastrand cross-links of purine bases leading to single strand DNA breaks.^41^ Taken together, these results confirm DNA damage as the primary mechanism of **Pt1** and **Pt8**.

**Figure 5.**
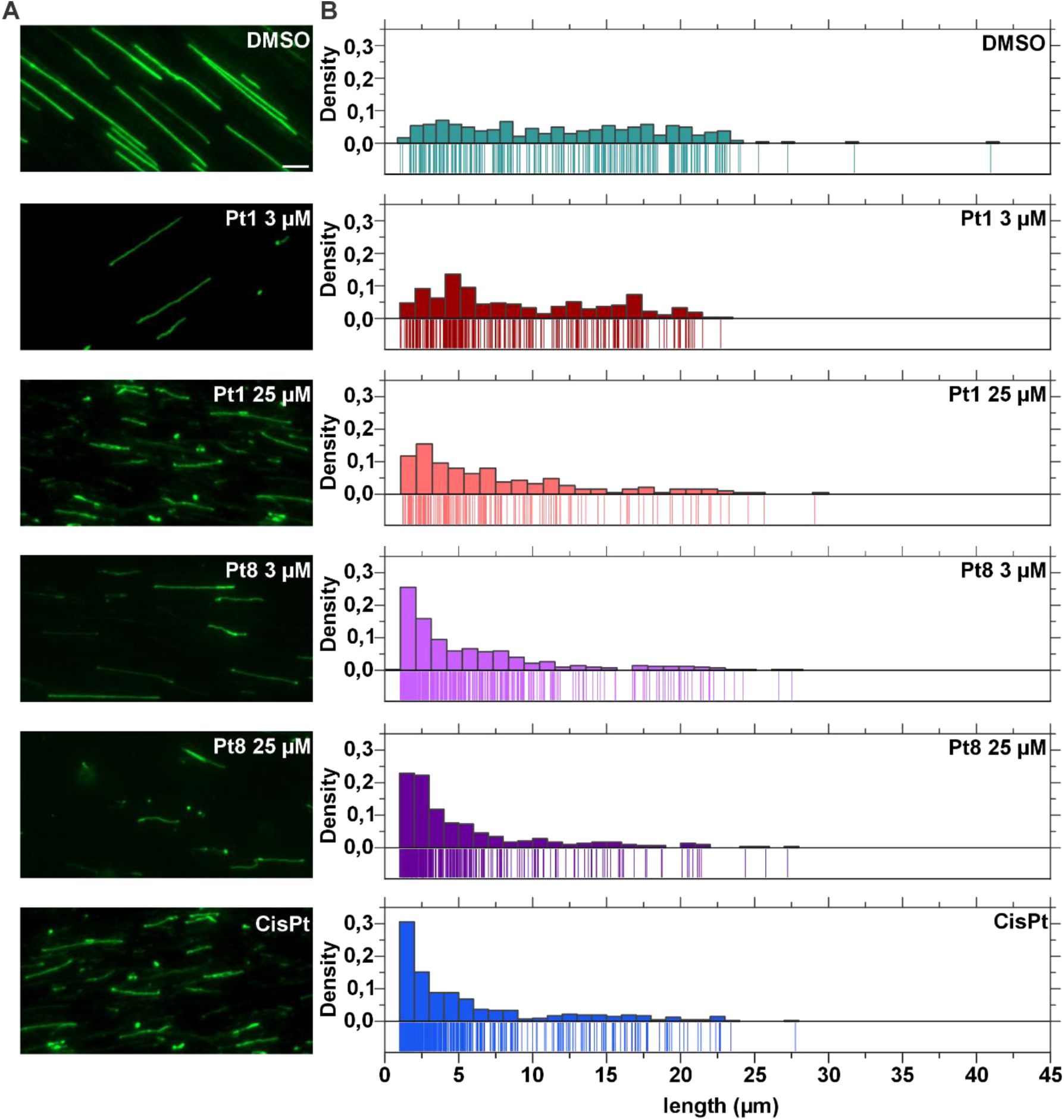
Single-molecule imaging of YOYO-1-labeled λ-DNA. (**A**) Fluorescence microscopy images of λ phage DNA exposed to the different lead compounds. DMSO was included as negative control. Scale bar 5 µm. (**B**) DNA strand length distribution measured from microscopy images (pooled from three biological replicates).

Interestingly, **Pt1**, which was the most potent antibacterial agent, had the mildest effects on λ-DNA. We wondered whether this observation may be due to the different concentrations, which were based on the OSC determined for *B. subtilis* cultures. Therefore, we additionally tested 25 µM **Pt1** and 3.125 µM **Pt8** to directly compare equimolar concentrations. While an increase of **Pt1** clearly showed stronger DNA fragmentation, the decrease of **Pt8** did not diminish its effect (**Figure 5**). These results suggest that **Pt8** is a more potent DNA-damaging agent.

### Compound uptake

Following this observation, we hypothesized that the difference in antibacterial activity may be due to less efficient uptake of **Pt8** compared to **Pt1**, and that this is likely also the reason underlying the poor antibacterial activity of **CisPt**. Therefore, we used inductively-coupled plasma mass spectrometry (ICP-MS) to assess compound uptake into MRSA based on detection of cellular Pt levels.^42^ Indeed, **Pt1** showed much more efficient uptake than **Pt8**, while **CisPt** showed the lowest uptake (**Figure 6A** and **S12**). Although the uptake percentages differ drastically between the lead compounds and **CisPt**, the absolute amounts of platinum in bacteria after treatment (**Pt1**: 3.5 µg/L, **Pt8**: 4.0 µg/L, **CisPt**: 2.6 µg/L) did not show a marked difference. While the variation in uptake clearly plays an important role in the differences in antibacterial activity, other molecular factors likely contribute as well.

**Figure 6.**
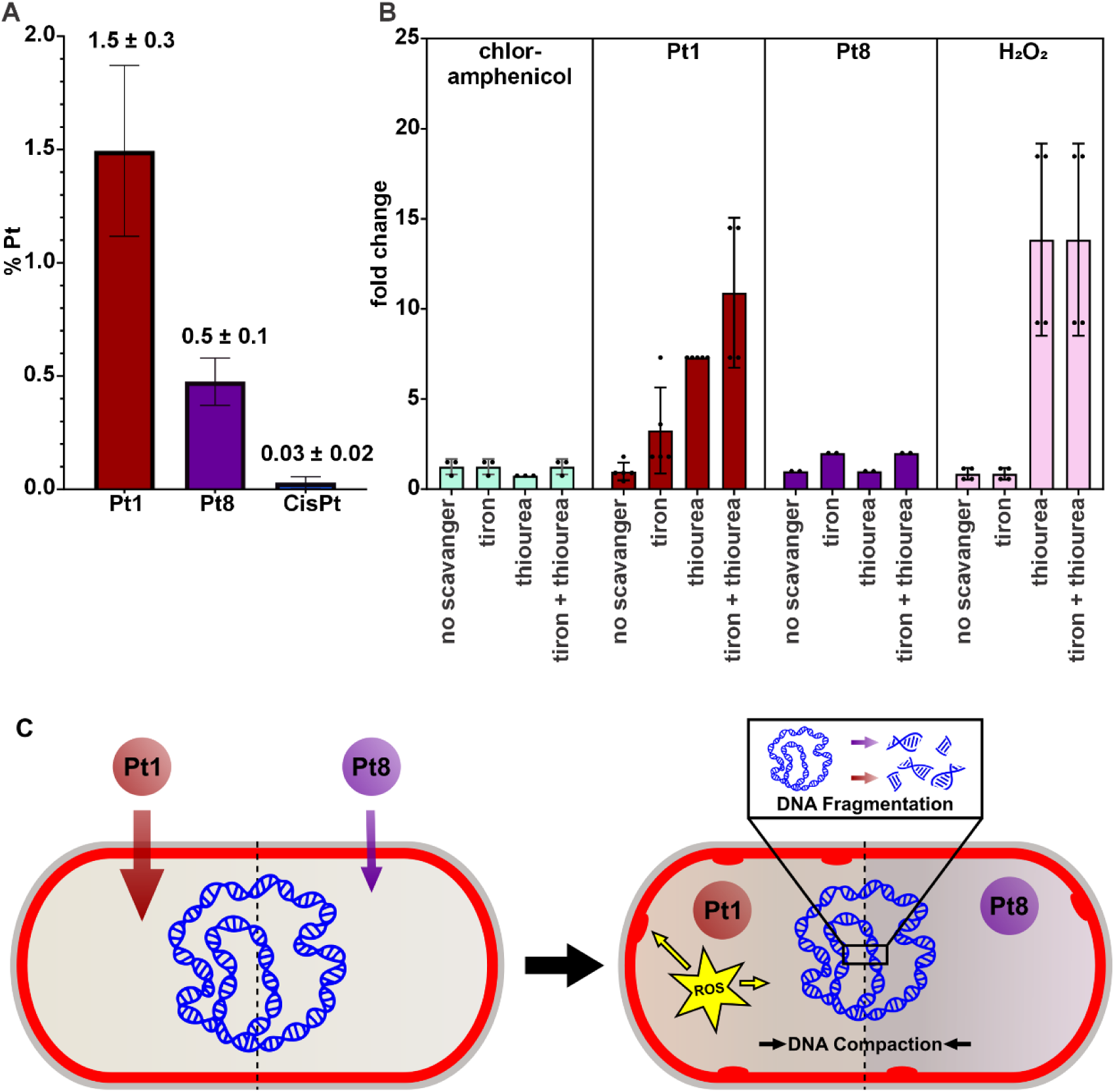
Compound uptake and ROS play a role in the increased potency of **Pt1** compared to **Pt8**. (**A**) Percent Pt content of MRSA cell lysate after 60 min treatment with **Pt1**, **Pt8**, and **CisPt**. (**B**) MIC fold changes (increase) in the presence of ROS scavengers compared to regular medium without scavengers). Error bars represent standard deviation of at least three biological replicates. (**C**) Mode of action model based on phenotypic analysis of *B. subtilis*. Both Pt compounds damage DNA, causing DNA strand breaks and fragmentation, resulting in nucleoid compaction and reduced DAPI staining. **Pt1** additionally generates ROS, mainly hydroxyl radicals, further increasing its potency through multimodal cell damage, e.g., on the cell membrane.

### Reactive oxygen species

Metal complexes often induce oxidative stress in cells, which can significantly contribute to DNA damage, mostly through generation of hydroxyl radicals. While platinum cannot substitute for iron or copper in a Fenton reaction to create hydroxyl radicals from hydrogen peroxide, it has been shown to increase hydroxyl radical-mediated DNA damage^43^ in a system capable of the Fenton reaction. Bacterial cells contain significant amounts of iron and the Fenton reaction is a common source of hydroxyl radicals.^44^ Exacerbation of DNA-damaging reactive oxygen species (ROS) in addition to direct DNA-damaging action constitutes a mechanism that could explain the higher antibacterial activity of **Pt1** compared to **Pt8**, in addition to more efficient uptake. To test whether ROS play a role in the activity of the compounds, a scavenger assay was employed. In this modified MIC assay, the activity of a test compound is determined in scavenger-free medium, medium containing the superoxide scavenger tiron, the hydroxyl radical scavenger thiourea, or a mix of both.^39,45^ If ROS contribute to antibacterial activity, an increase of the MIC value is expected in medium containing the scavenger specific to the ROS that is generated. Thus, the antibacterial activity of hydrogen peroxide, a hydroxyl radical donor, was not significantly affected by the superoxide scavenger tiron, but in presence of the hydroxyl radical scavenger thiourea, the MIC increased by a factor of 14, and no additive effect was observed when tiron and thiourea were combined (**Figure 6B**). The same trend was observed with **Pt1**, which showed a threefold MIC increase in medium containing tiron, a tenfold increase in the presence of thiourea, and a 13-fold increase in medium containing both scavengers. These results suggest that mostly hydroxyl radicals and to a lesser extent superoxide contribute to the activity of **Pt1**. Importantly, **Pt8** showed no notable MIC increase in the presence of ROS scavengers, confirming our notion that the increased activity of **Pt1** is due to the generation of ROS in addition to direct DNA damage. This finding could also explain the tendency of **Pt1** to induce Nile red foci, since ROS can also damage membrane lipids^46^, likely contributing to the potency of the compound (**Figure 6C**).

### Advanced biological evaluation of Pt1

Amongst all evaluated PtCOD derivatives, **Pt1** displayed the best properties. It showed nano to low micromolar antibacterial activity across a panel of Gram-positive bacteria, displayed low toxicity, both in cell lines and in live *G. mellonella,*^25^ and possesses a new mode of action. Furthermore, **Pt1** reduced the fungal load in a *G. mellonella* model of *C. albicans* infection in previous studies.^24^ Thus, we further characterized its properties towards *in vivo* applications.

### Stability

The stability of **Pt1** was examined by ^1^H and ^195^Pt NMR. NMR of approximately 7 mM **Pt1** in DMSO-*d6* was recorded immediately after preparation, and after 24 h and 48 h of incubation at 37 °C (**Figure S13-14**). No differences in the NMR spectra could be identified, suggesting that **Pt1** remains intact for at least 48 h under the tested conditions.

### Combination with other antibiotics

Due to its unique DNA-targeting effect, we aimed to study whether **Pt1** could synergistically interact with approved antibiotics. We conducted a combination study with nine antibiotics commonly used against *S. aureus* (ATCC 29213). Neither synergistic nor antagonistic effects were observed with any of the tested antibiotics, indicating that the mode of growth inhibition of **Pt1** is unrelated to that of other antibiotics (**Table S7**).

### Resistance development

We then explored the potential of **Pt1** to induce resistance in *S. aureus* (ATCC 29213), by growing the culture in the presence of compound at ½ MIC for 36 days. Only a minimal increase in the MIC (0.125 to 0.25 µg/mL) was observed over this time period, which is in stark contrast to a 256x fold MIC increase of the antibiotic levofloxacin under the same conditions (**Figure 7A**).

**Figure 7:**
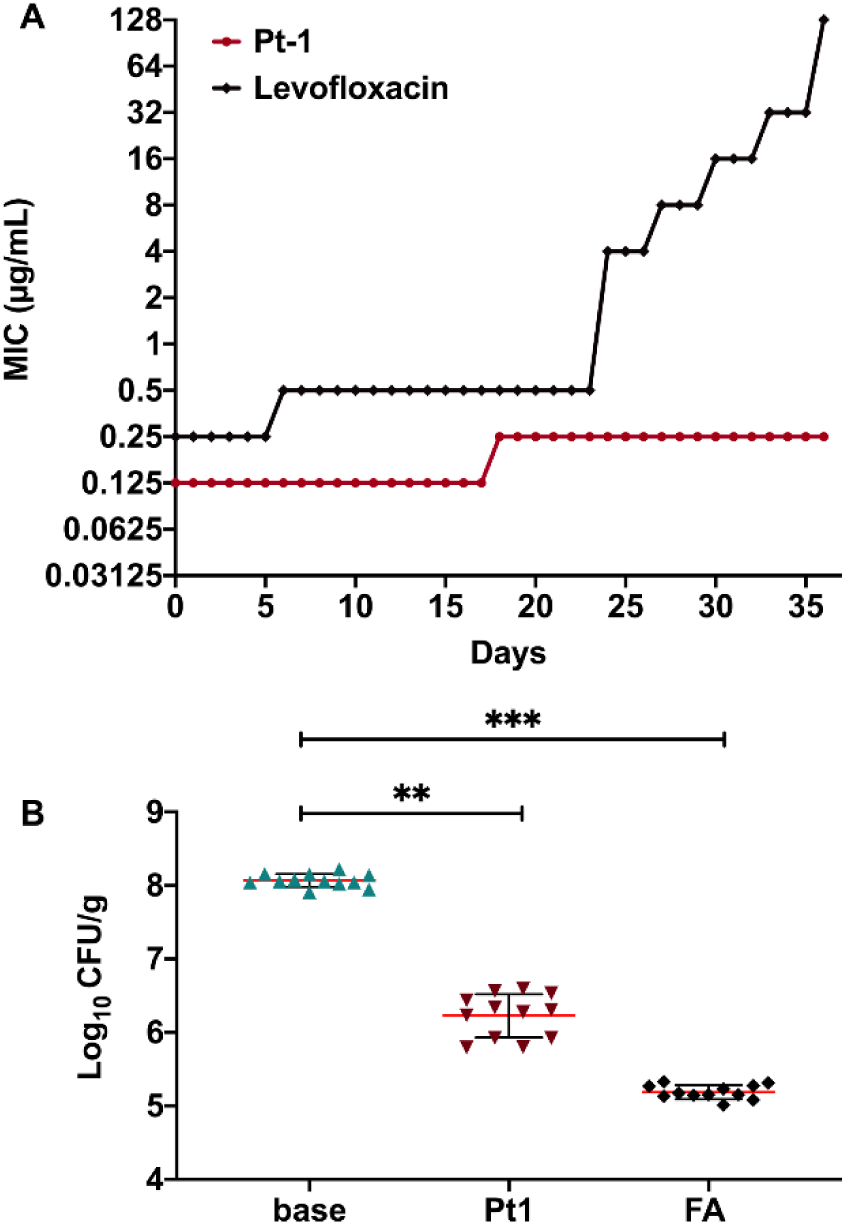
Resistance development and *in vivo* efficacy of **Pt1**. (**A**) Growth pattern of *S. aureus* ATCC 29213 grown in the presence of **Pt1** and levofloxacin at 0.5x MIC for 36 days. (**B**) Reduced bacterial load in an *in vivo* skin infection model in mice. Fusidic acid (**FA**) was used as positive control.

We then used the strain resulting from 36 passages to interrogate whether cross-resistance to other antibiotics could be observed (**Table S8**). As expected, based on the preceding combination study and the very low resistance, no cross-resistance was observed, further emphasising the unique nature of the mechanism of **Pt1**.

The efficacy of **Pt1** in suppressing bacterial growth after a brief exposure to the compound was assessed by studying its post-antibiotic effect (PAE). **Pt1** displayed a PAE of ∼5.5 hr compared to the control drugs levofloxacin and vancomycin, which displayed PAEs of 1.5 h and 2.5 h, respectively, at 10x MIC (**Table S9**). Taken together, **Pt1** exhibits a long PAE while not inducing resistance, thus positively impacting its dosing regimen.

### *In vivo* skin infection model

Following establishment of a favourable biological profile for **Pt1**, we proceeded to evaluate its efficacy in a topical *S. aureus* skin infection model (**Figure 7B**). To this end, 5–7 weeks-old Swiss mice (22-25 g) were given a superficial wound on the dorsal surface, followed by infection with a bacterial load of ∼10^7^ CFU/mL. Mice treated with a 2% **Pt1** cream formulation displayed a ∼1.9 log10 CFU/g reduction in bacterial load compared to the mice in the vehicle-treated group. Thus, **Pt1** reduced the bacterial load to an extent that is comparable to the established treatment (fusidic acid, ∼2.9 log10 CFU/g). This experiment demonstrated the antibacterial *in vivo* efficacy of a platinum-containing compound in a mouse model.

## Discussion

In the search for novel classes of antimicrobial agents we have identified platinum cyclooctadiene complexes with promising activity against Gram-positive bacteria and low toxicity against healthy human cells. Structure-activity relationship studies in this and previous work revealed several key insights. The halogen ligands on the platinum center seem to be crucial as alteration to methyl or hydroxy-containing ligands led to reduced activity and/or increased toxicity.^25^ More scope remains to explore the effect of other, potentially bidentate, ligands (in conjunction with the COD) on the antibacterial activity and toxicity of these compounds. Switching the metal center to palladium was shown to result in complete loss of antibacterial properties and significant haemolysis^25^, letting us conclude that the platinum is key. Finally, in this work, we explored the synthetically more challenging modifications of the COD ligand. We observed that any modification of the ligand at the double bond position resulted in loss of antibacterial activity in the final platinum compound. Conversely, modification on the allylic COD position maintained some of the antibacterial properties. However, the best modified complex, **Pt8** was still significantly less active than the original hit- compound **Pt1**. Both **Pt1** and **Pt8** showed broad activity against a range of clinical isolates of drug-resistant *S. aureus* and low cytotoxicity and haemolysis values. The structural similarity of **Pt1** with the anticancer drug **CisPt** prompted us to re-examine the antibacterial properties of the latter, which had been noted to induce bacterial elongation in the past.^17^ In our hands, **CisPt** showed no antibacterial activity up to 100 µM against any of the evaluated bacterial strains. However, we did observe slight growth defects and a mild cell elongation phenotype in *B. subtilis* treated with 100 µM **CisPt** (**Figure S15**). Uptake studies suggest that its poor activity is partially due to inefficient membrane translocation of the compound. Indeed the group of Nolan has shown that coupling platinum chemotherapeutics to siderophores leads to significant antibacterial activity.^47,48^

We examined the antibacterial mechanisms of action of the lead compounds using BCP, which revealed clear effects on the bacterial DNA. **Pt1** additionally caused a noticeable, yet not significant formation, of Nile red foci in the cell membrane. Thorough investigation of the effects of **Pt1** and **Pt8** on membrane integrity and potential as well as on cell wall synthesis and cell division returned negative. Similarly, no distinct phenotype on fluorescent transcription and translation reporters could be observed, letting us conclude that these platinum compounds do not affect bacterial protein production. Taken together, these results indicate that the main target of both **Pt1** and **Pt8** is bacterial DNA.

Closer investigation of this target, through RecA as reporter for induction of the SOS response to DNA damage, revealed that both **Pt1** and **Pt8** induce a rapid response at 10 min after treatment that drops off over time. This is distinct from **CisPt**, which induces an increasing response over the 60 min measurement period. Importantly, we could demonstrate that all three platinum compounds are able to directly cause DNA fragmentation using single molecule imaging of fluorescently labelled λ-DNA. This effect was milder for **Pt1** compared to **Pt8**.

Studying the effect of different ROS quenchers on the antibacterial activity of the hit compounds revealed that the activity of **Pt1** is strongly suppressed in the presence of a hydroxyl radical scavenger. While a superoxide quencher alone did not affect activity, addition of both quenchers together achieved an even stronger reduction in **Pt1** activity, suggesting that hydroxyl radicals considerably contribute to its activity while superoxide may play a secondary role. In contrast, the activity of **Pt8** was not affected by any ROS quenchers.

Taken together, we propose a mode of action model for this new compound class (**Figure 6C**). Both **P1** and **Pt8** directly bind to and damage bacterial DNA, leading to strand breaks and consequently DNA fragmentation. While **Pt8** is the more efficient DNA damaging agent, **Pt1** shows higher antibacterial activity. This seeming conundrum can be explained by (i) more efficient compound uptake, **Pt1** being taken up 50x more efficiently than **CisPt** and 3x more efficiently that than **Pt8**, and (ii) by an additional mechanism of **Pt1**, namely the generation of ROS. This multimodal effect of **Pt1** further explains its tendency to induce membrane stress (Nile red membrane foci), possibly through lipid peroxidation, as well as the absence of any high-level resistance emerging over 36 passages.

Finally, we evaluated the ability of **Pt1** to treat a topical *S. aureus* skin infection in a mouse model and observed a ∼1.9 log10 CFU/g reduction in bacterial load. Altogether, these observations make **Pt1** a highly interesting lead compound with potent activity against critical antibiotic-resistant pathogens, low cytotoxicity, a unique and multimodal mechanism of action that does not permit effective resistance development, and potent efficacy in an *in vivo* mouse infection model.

DNA is an underexplored antibacterial target and only few antibiotics that specifically target DNA in bacteria are on the market.^49^ Metal compounds such as ruthenium polypyridyl complexes have long been known to intercalate DNA through non-covalent interactions.^50,51^ The herein-reported class of platinum compounds represents the first non-cisplatin-based compounds with a demonstrated direct DNA effect, high selectivity for bacteria and *in vivo* efficacy, validating further explorations into the development of platinum compounds as new antibiotic class with a highly attractive biological profile.

## Methods

### Syntheses of COD ligands and Pt(II) complexes

The detailed synthetic procedures and characterization of compounds can be found in the supplementary information (**S1. General Material** and **S2. Synthetic Procedures**).

### Antibiotic susceptibility testing of lead compounds

#### Determination of antibacterial activity: Minimum inhibitory concentration assay

The MIC protocol was adapted from Cai *et al*.^52^ A single colony of bacteria was grown in Luria-Bertani (LB) medium overnight at 37 °C. Stock solutions of 5 mM of the samples were prepared in DMSO and diluted to a starting concentration of 100 μM in Mueller Hinton (MH) medium. The overnight culture was diluted and incubated until the OD600 reached 0.6–1.0. The bacteria concentration was measured by measuring the optical density at 600 nm and diluted to an OD600 of 0.022 in MH medium. 5 μL of the diluted bacterial solution was used to inoculate 150 μL of the sample solutions, resulting in a final inoculation of about 5 × 10^5^ cells/mL. The plates were then incubated at 37 °C for 18 hours. For each assay, a control of broth only and a growth control of broth with bacterial inoculum without antibiotics were included in two columns of the plate. Polymyxin B and vancomycin were used as control antibiotic for *E. coli,* MRSA, *B. subtilis,* and MSSA, respectively. The growth was measured by analysing the absorbance of the bacterial suspension at 600 nm. Each experiment was performed in triplicate.

### Determination of antibacterial activity: Minimum inhibitory concentration assay against selection of Gram-positive and Gram-negative strains^53^

The data is given in supporting information (**Table S4**). Before starting the experiment, a single colony was picked from MHA plate, inoculated in MHBII and incubated overnight at 37 °C with shaking for 18–24 hours to get the starter culture. Antibiotic susceptibility testing was conducted according to the CLSI guidelines using the broth microdilution assay. 10 mg/mL stock solutions of test compounds were prepared in DMSO. Bacterial cultures were inoculated in MHBII and optical density (OD) was measured at 600 nm, followed by dilution to achieve ∼10^6^ CFU/mL. The compounds were tested from 64–0.5 mg/L in two-fold serial diluted fashion with 2.5 μL of each concentration added to well of a 96-well round bottom microtiter plate. Later, 97.5 μL of bacterial suspension was added to each well containing either test compound or appropriate controls. The plates were incubated at 37 °C for 18-24 h following which the MIC was determined. The MIC is defined as the lowest concentration of the compound at which there is absence of visible growth. For each test compound, MIC determinations were carried out independently three times using duplicate samples.

### Determination of cytotoxicity of lead compounds against HEK293T cells^52^

Stock solutions of 5 mM of the compounds were prepared in DMSO and the stock solution of cisplatin and oxaliplatin were prepared in 1% NaCl as 4.3 mM and 2.5 mM respectively. The control for the experiments was DMSO. HEK293T cells were cultivated in DMEM high glucose supplemented with 10% heat-inactivated fetal bovine serum, 100 U/mL of penicillin and streptomycin and maintained at a temperature of 37 °C with 5% CO2. Initially, 4000/well of cells were seeded into 96-well plates and incubated for 24 hours before being exposed to compounds at concentrations ranging from 0.78, 50, and 100 µM. After 24 hours of incubation, the compounds were added and serially diluted and incubated for 24 and 48 hours at 37 °C 5% CO2, the cells were stained with alamar Blue, and the fluorescence excitation/ emission 540 nm/ 590 nm was measured using 96 well-plate reader. CC50 is defined as the lowest concentration of compound which leads to a 50% reduction in cell viability compared to untreated and DMSO control. Each experiment was repeated in triplicate.

### Determination of Cytotoxicity of Lead Compounds against Vero Cells

Cell toxicity was performed against Vero cells using the MTT assay^54^. ∼10^3^ cells/well were seeded in 96 well plate and incubated at 37 °C with 5% CO2 atmosphere. After 24 hours, compound was added ranging from 100-12.5 µg/mL concentration and incubated for 72 h. After the incubation was over, MTT was added in each well, incubated at 37°C for further 4 h, residual medium was discarded, 0.1 mL of DMSO was added to solubilise the formazan crystals and OD540 was taken for the calculation of CC50. Doxorubicin was used as positive control and each experiment was repeated in triplicate.

### Determination of haemolytic activity in hRBCs: Haemolysis assay

The compounds were tested on human red blood cells (hRBCs) using a haemolysis assay, adapted from previously published procedure^55^. 1.5 mL of whole blood was centrifuged at 3000 rpm for 15 minutes at 4 °C, and the plasma was discarded. The hRBC pellet was washed three times with PBS (pH 7.4) and then resuspended to a final volume of 10 mL in PBS. For the determination of HC50 of the lead compounds, the samples serially diluted starting from 200 μM. Samples stock solution was 5 mM in DMSO. Each plate included a blank medium control (PBS) and a haemolytic activity control (0.1% Triton TM X-100). The hRBC suspension was incubated with the samples in PBS in a V-shaped 96-well plate for 4 hours at 20 °C. After the incubation, 100 μL of supernatant was carefully pipetted to a flat bottom, clear 96-wells plate. Haemolysis was measured by analysing the absorbance at 540 nm using a plate reader. The percentage of haemolysis at each concentration was determined and the HC50 was calculated by inhibitor vs. normalized response fit. The haemolytic activity was measured by analysing the absorbance of free hemoglobin in the supernatants at 540 nm. Minimum inhibitory concentration (MHC) was determined by eye. Each experiment was repeated in triplicate.

### Bacterial strains and growth conditions for mode of action analysis: Determination of optimal stressor concentrations (OSC)

To establish a starting concentration for the OSC assay, MICs against parent strain 168CA were determined under conditions equivalent to those used for mode of action assays. MICs were determined according to the Clinical Laboratory Standardization Institute (CLSI) in a serial microdilution assay.^36^ The compounds were serially 2-fold diluted in a 96-well plate and *B. subtilis* (168CA) was added to a final cell count of 5 x 10^5^ CFU/mL. Cultures were incubated for 16 h at 37 °C under constant agitation. MICs were used as basis for OSC determination. To this end, growth curves were recorded using optical density measurements. Culture volumes of 20 mL were grown until reaching an OD600 of 0.3. Subsequently, the parent culture was split into 2 ml aliquots, followed by addition of different MIC multiples of Pt compounds.^56^ An untreated culture served as control. Growth was followed until cultures reached the stationary phase. A concentration achieving 50-70% growth inhibition in exponential phase was selected as optimal stressor concentration. The following concentrations were selected: 3.125 µg/mL **Pt1**, 25 µg/mL **Pt8**, 100 µM **CisPt**. As controls, 0.23 µM nisin, 0.53 µM µM gramicidin, 100 µM CCCP, 0.67 µM vancomycin, 1.2 µM rifampicin, and 323.12 µM chloramphenicol were included as controls for specific assays as appropriate.

### Fluorescence light microscopy

Microscopy images of bacteria were acquired on a Nikon Eclipse Ti2 equipped with a CFI Plan Apochromat DM Lambda 100X Oil objective (N.A. 1.45, W.D. 0.13 mm), a Photometrics PRIME BSI camera, a Lumencor Sola SE II FISH 365 light source, and an Okolab temperature incubation chamber. Images were obtained using the NIS elements AR software version 5.21.03/ 5.42.02. DNA stretching images were obtained on a Zeiss Observer.Z1 fluorescence microscope equipped with a Colibri 7 LED illumination system and an Andor iXON Ultra EMCCD camera. Image analysis was perfomred using ImageJ^57^, MicrobeJ^58^, and ObjectJ^59^.

### Bacterial cytological profiling (BCP)^56^

BCP was performed according to Wenzel *et al*..^56^ In short, *B. subtilis* strain MW54 (*PrpsD-gfp*) was grown to an OD600 of 0.3 prior to splitting of parent cultures and addition of antibiotics. Samples were taken for microscopy after 30, 60, and 120 min of antibiotic treatment. Five minutes prior to sampling, cultures were stained with 0.5 µg/mL Nile red and 1 µg/mL DAPI for 5 min. Samples of 0.5 µL were then spotted onto 1% agarose slides^60^ and images were taken immediately.

### BCP image analysis

Cell length and DAPI intensity were analyzed using MicrobeJ^58^. To this end, the fit shape, rod-shaped setting was used with a minimum area of 0.5 µm, while all other settings remained at default. *B. subtilis* chains were seperated based on the membrane stain. Membrane stress was counted using the ObjectJ plugin^59^, using the membrane setting to seperate individual cells in chains. Membrane damage was expressed as ratio of cells with visible Nile red foci and total detected cells. DNA compaction was analyzed by determining the area of the DNA and the whole cell are using advanced weka segmentation. Cells were segmented according to the GFP signal. Automatic segmentation that did not fit the DNA or cell shape was corrected by manual tracing. Cells that were out of focus or lysed were excluded from the analysis. Statistical analysis was peformed using GraphPad Prism 10 as specified in the individual figure legends.

### Propidium iodide pore assay^56^

Propidium iodide staining was performed as described previously *B. subtilis* 168CA was grown to an OD600 of 0.3 prior to splitting of the parent culture and treatment with the different antibiotics. After 15 and 45 min, samples were stained with 13.3 µg/mL propidium iodide for 15 min (total treatment times 30 and 60 min), and washed 3 times with phosphate-buffered saline (PBS). Fluorescence signals were recorded at an excitation wavelength of 535-15 nm and an emission wavelength of 617-20 nm in a BMG Clariostar Plus plate reader.

### DiSC3(5)-based membrane potential measurements^56^

The membrane potential was measured according to Winkel *et al.* with minor modifications.^60^ *B. subtilis* 168CA was grown in LB containing 50 µg/mL bovine serum albumin (BSA). Cells were transferred to black 96-well polystyrene microtiter plates after reaching an OD600 of 0.3. DiSC3(5) was added to a final concentration of 1 µM, maintaining a constant concentration of 1% DMSO to avoid precipitation of the dye. After the fluorescence baseline had stabilized, antibiotic compounds were added, and measurements were continued for 30 min. Fluorescence was recorded at an excitation wavelength of 610-30 nm and an emission wavelength of 675-50 nm in a BMG Clariostar Plus plate reader.

### GFP localization studies^39^

*B. subtilis* strains were grown in LB containing appropriate inducer concentrations as specified in **Table S10**. After reaching an OD600 of 0.3, cultures were split and treated with antibiotics. Samples were withdrawn after 30 and 60 min of treatment, spotted on glass slides covered with 1% agarose films^60^, and immediately imaged.

### Translation inhibition assay

*B. subtilis* strain TB35 (*Pxyl-minD-gfp*) was grown in LB without inducer. After reaching an OD600 of 0.3, antibiotics and inducer (0.1% xylose) were added consecutively. Samples were withdrawn for microscopy after 60 min of treatment, spotted on glass slides covered with 1% agarose films^60^, and imaged without delay.

### Single-molecule imaging of DNA

DNA stretching was performed on silanized glass coverslips (18 × 18 mm^2^) using a silanization protocol adapted from Wei *et al*..^61^ Briefly, glass coverslips were arranged in a coverslip rack and submerged overnight in acetone solution containing 1 % APTES and 1% ATMS (v/v). The activated coverslips were then rinsed with a (2:1 v/v) acetone:water solution and dried with air purging. Prior to imaging, 138.6 ng of λ-DNA were treated with the respective compounds for 10 min, followed by staining with 320 nM YOYO-1 in 0.5× Tris-borate-EDTA (TBE) buffer. To minimize photobleaching, 2% β-mercaptoethanol (BME) was added. 3.2 μL of the stained DNA sample were placed at the interface between a silanized coverslip and a clean microscope slide to stretch the DNA molecules. A band-pass excitation filter (475/40 nm) and a band-pass emission filter (530/50 nm) were used for detecting YOYO-1 fluorescence.

### DNA length analysis

To analyse the length of individual DNA molecules, the ImageJ^57^ plugin MicrobeJ^58^ was used. DNA molecules were detceted using medial axis detection with 1 µm minimum length, 0.3 to 1.5 µm width range with 0-0.04 variation, and a maximum circularity value of 0.7. DNA molecules that were not correctly detected were selected manually. DNA length measurements were analyzed and visualized with Origin 2023.

### ROS scavenger assay

LB media were prepared to contain either 10 mM tiron, 150 mM thiourea, 10 mM tiron and 150 mM thiourea, or no scavanger. Serial two-fold dilutions of the different antibiotics were prepared in the respective media and transferred to 96-well plates. *B. subtilis* 168CA was added to a final cell count of 5 x 10^5^ CFU/mL and cultures were incubated at 37 °C for 16 h under constant agitation. The OD600 was meassured in a BMG Clariostar Plus plate reader.

### Antibiotic uptake assay^42^

A single colony of bacteria was grown in Luria-Bertani (LB) medium overnight at 37 °C. Stock solutions of 5 mM of the samples were prepared in DMSO for **Pt1** and **Pt8**, and 5.3 mM in 0.9% NaCl(aq) for **CisPt**. The overnight culture was diluted in 80 mL LB medium and grown until it reached an OD600 of 0.3-04. 10 mL of the culture were distributed to falcon tubes and cells were stressed with 1.56 µM **Pt1**, 6.25 µM **Pt8** and 100 µM **CisPt** for 30 and 60 min. Then, the cells were harvested at 4500 rpm for 10 min and the supernatant was collected. The cell pellets were washed with 1 mL washing buffer (100 mM Tris, 1 mM EDTA, pH 7.3) and centrifuged at 4500 rpm for 10 min each. Subsequently, cell pellets were resuspended in 1 mL PBS (pH 7.5), treated with 500 µL lysing buffer (0.1 M glycine-HCl, pH 2.5), and incubated overnight at room temperature. Following incubation, samples were centrifuged at 14000 rpm for 5 min and supernatant (cell lysis) and pellet (cell debris) were collected. After pellets were resuspended in 750 µL PBS (pH 7.5), all samples were frozen in liquid N2 and lyophilized overnight. Then, samples were treated with 300 µL nitric acid (69.7% (w/mL) in double-distilled water) and incubated for a week at room temperature, followed by incubation at 65°C for 4 hours. Then, the samples were diluted to 2% nitric acid content with double-distilled water and filtered through 0.22 µm syringe filters.

The ICP-MS system was operated in kinetic energy discrimination (KED) mode and the collision gas was He50 (AlphaGaz), pre-treated with a helium cell gas filtration system (Perkin Elmer, N8146004). The gas flow for KED was 5.3 mL/min. 2% HNO3 (*w*/*w*) was used as matrix for all sample, calibration, and internal standard (IS) solutions. 2% HNO3 (*w*/*w*) itself was prepared from 69.3% (*w*/*w*) HNO3 (BASF, < 1 ppb ECME), diluted with ultrapure water (*ρ* = 18.2 MΩ˖cm). Calibration solutions of 0.1, 1, 10 and 50 μg/L Mg, Mn, and Pt were prepared. For Mg and Mn, the ‘ICP multi-element standard solution VIII’ (Merck, 1.09492) was used as a stock solution. ‘Platinum Standard for ICP’ (Sigma-Aldrich, 19078) was used as a stock solution for Pt. All elements were combined in the same calibration solutions. 20 μg/L Ir and 20 μg/L Y in 2% HNO3 were used as internal standards (IS) during the ICP-MS analysis. Y was used as IS for Mg and Mn. Ir was used as IS for Pt. Stock solutions were ‘Iridium ICP standard’ (Supelco, 1.70325) and ‘Yttrium ICP standard’ (Merck, 1.70368). Both elements were combined in the same IS solutions. The respective isotopes measured were: ^24^Mg, ^26^Mg, ^55^Mn, ^89^Y, ^195^Pt, and ^193^ Ir. Each sample measurement consisted of 7 replicates and no automatic corrections for potential interferences were applied.

### Determination of Pt1 stability

The stability of **Pt1** complex was measured by ^1^H and ^195^Pt NMRs. Approximately 7 mM of **Pt1** in DMSO-*d6* was prepared and NMR spectra were immediately measured. Measurements were repeated after the 24 h and 48 h of incubations at 37°C.

### Drug interaction study

Interaction of **Pt1** with FDA-approved antibiotics was determined using chequerboard assays following CLSI guidelines. All compounds were freshly prepared for the experiment. Eight serial two-fold dilutions of **Pt1** were prepared from 1 to 0.0078 µg/mL. Test antibiotics were prepared in 12 serial two-fold dilutions as follows: ceftazidime 64–0.03125 µg/mL, daptomycin and vancomycin 8–0.0039 µg/mL, gentamicin, levofloxacin, meropenem, and minocycline 2- 0.00095 µg/mL, linezolid 16–0.0078 µg/mL, rifampicin 0.06–0.000014 µg/mL. Dilution series were inoculated with 5×10^5^ *S. aureus* ATCC 29213 CFU/mL and incubated for 16-20 h at 37 °C. Following incubation, the fractional inhibitory concentration index (FICI) was calculated with the formula FICI=FICA+FICB, whereby FICA= MIC of drug A in combination/MIC of drug A alone and FICB= MIC of drug B in combination/MIC of drug B alone. A drug combination is considered synergistic when ∑FICI is ≤0.5, indifferent when ∑FICI is >0.5 to 4, and antagonistic when ∑FICI is >4.

### Postantibiotic effect

Post antibiotic effect (PAE) was determined as follows. Overnight culture of *S. aureus* ATCC 29213 were diluted in MHBII to ∼10^5^ CFU/mL, exposed to 1x and 10x MIC of **Pt1**, levofloxacin, and vancomycin, and incubated at 37°C for 1 h. Following incubation, cultures were centrifuged and washed twice with pre-warmed MHBII to remove any traces of antibiotics. Finally, cells were resuspended in drug-free MHBII and incubated further at 37 °C. Samples were taken every 1 h, serially diluted, and plated on tryptic soy agar (TSA) for enumeration of CFU. PAE was calculated as PAE = T – C; where T is the difference in time required for 1 log10 increase in CFU vs. CFU observed immediately after the removal of the drug, and C in a similarly treated drug-free control.

### Resistance induction^62^

*S. aureus* ATCC 29213 was serially passaged in the presence of 0.5×MIC of either **Pt1** or levofloxacin (positive control). The MICs of the passaged cultures were determined every 3 days and compared to that of wild type *S. aureus* ATCC 29213. MIC changes were recorded for 36 passages.

### Murine skin infection model^63^

The *in vivo* efficacy of **Pt1** was determined in a superficial skin infection model. Briefly, male Swiss mice, 4–6 weeks old, weighing ∼22–25 g, were used throughout the study. Mice were caged alone in an individually vented cage (IVC) to check cross-contamination and to maintain aseptic conditions throughout the experiment. Ketamine and xylazine were prepared in distilled water, 40 mg/kg and 8 mg/kg of body weight, respectively, and injected intraperitoneally (IP) as a mixture of 100 μL in each mouse for anaesthesia. Then, we proceeded with the removal of fur by applying depilatory cream. The area was cleaned with sterile distilled water. Approximately 2 cm^2^ of skin area was scratched until it became visibly reddened and glistening without any bleeding. For bacterial infection, a 10 μL droplet containing 10^7^ CFU/mL of *S. aureus* ATCC 29213 was applied to the reddened area of skin. To confirm the infection, post 4 h, untreated mice were sacrificed, skin homogenized, and various dilutions of the treatment groups (6 mice per group) were plated on TSA plates. Following this, treatment with 2% fusidic acid (positive control), 2% **Pt1**, and base (vehicle) was started, followed by a second dose post 16 h from the first dose. Henceforth, the drug regimen was comprised of twice daily application (in the morning and the evening, with an 8 h interval) for 4 days. Control mice were regularly treated with 25–30 mg each of 2% fusidic acid (LEO Pharma, Ballerup, Denmark), 2% infuzide, and base (vehicle). All groups were sacrificed post-18 h of the last dose to avoid carryover effects of the treatment group. Around ∼2 cm^2^ of wounded skin were excised post sacrifice and homogenized in 500 μL PBS in 2 mL MP tissue grinding Lysing Matrix F tubes using MP FastPrep-24 set at 4.0 M/S for 30 s (3 cycles). Dilutions were plated post homogenization on TSA plates, followed by overnight incubation at 37 °C, to determine the bacterial CFU count in each treatment group. Each experiment was repeated three times in duplicate and the mean and SEM of Log10 CFU/g were plotted.

## Author contributions

Conceptualization: ÇÖ, ABS, SC, MW, AF

Data Curation: ÇÖ, ABS, RR, OAA, SC, MW, AF

Formal Analysis: ÇÖ, ABS, RR, OAA

Funding Acquisition: ABS, FW, MW, AF

Investigation: ÇÖ, AA, RM, DS, ABS, RR, OAA

Methodology: ÇÖ, AA, RM, DS, ABS, FW, MW

Project Administration: FW, MW, AF Resources: FW, SC, MW, AF

Software:

Supervision: ABS, OAA, FW, SC, MW, AF

Validation: ÇÖ, ABS, OAA

Visualization: ÇÖ, AA, RM, DS, ABS, SC, AF

Writing – Original Draft Preparation: ÇÖ, ABS, MW, AF

Writing – Review & Editing: ÇÖ, ABS, RR, OAA, FW, SC, MW, AF

## Acknowledgements

We sincerely thank Prof. Jean-Louis Reymond for hosting and supporting our research group. Prof. Christoph von Ballmoos and Dr. Nicolas Dolder are acknowledged for giving the opportunity to use their microscope facility. Nicola Christian Lüdi is acknowledged for ICP-MS measurements. AF and CO gratefully acknowledge funding from the Swiss National Science Foundation Ambizione grant PZ00P2_202016 and from a Research Partnership Grant and support by the University of Bern. MW received funding from Chalmers University of Technology. RR and OAA were supported by the Wenner Gren Foundations (UPD-2021-0186 and UPD-2021-0036). Parts of the project were conducted during a study visit funded by the National Doctoral Programme in Infection and Antibiotics (NDPIA, to ABS) and a study visit of C.O. to Sweden was kindly funded by University of Bern.

