## Supplementary Information for "A Platinum Butterfly Effect: Small Changes Turn an Anticancer Drug into a Non-toxic Metalloantibiotic with In Vivo Efficacy"

ıx

**S1. General Materials**

**S2. Synthetic Procedures**

**S3. Supplementary Tables and Figures**

**TableS1:** Antibacterial activity of synthesized PtCOD complexes against *E. coli*.

**Table S2:** Antibacterial activity of the Pt-drugs against Gram-positive strains.

**Table S3:** Antibacterial activity of COD ligand and Pt precursor against *S. aureus* (CCUG 62707) and *E. coli* (NCTC 13476).

**Table S4:** Antibacterial activity of the lead complexes in µg/mL against a selection of Gram-positive and Gram-negative strains.

**Table S5:** HC_50_ values of synthesised **Pt** complexes against hRBC.

**Figure S1:** Growth-normalized % viability of HEK293T cells after treatment with **Pt1**, **Pt8**, CisPt and DMSO (negative control) after incubation for 24 h (**A**) and 48 h (**B**).

**Figure S2:** DMSO-normalized % viability of HEK293T cells after treatment with **Pt1**, **Pt8**, and **CisPt** after incubation for 24 h (**A**) and 48 h (**B)**.

**Table S6:** DMSO-normalized cytotoxicity (IC_50_ in µM) of **Pt1** and **Pt8** against HEK293T cells.

**Figure S3:** Percent viability of Vero cells upon treatment with **Pt1** (**A**) and **Pt8** (**B**) after incubation for 72 h.

**Figure S4:** Determination of optimal stressor concentrations (indicated in bold) of **Pt1**, **Pt8**, and **CisPt**.

**Figure S5:** Quantification of membrane stress measured as percent of *B. subtilis* bSS82 cells with Nile red foci.

**Figure S6:** Quantification of nucleoid compaction (**A**) and DAPI fluorescence intensity (**B**) corresponding to **Figure 2**, separated into individual biological replicates.

**Figure S7:** Fluorescence overlays of *B. subtilis* MW10 (*Pxyl-gfp-mreB*).

**Figure S8:** Localization of FtsZ in *B. subtilis* 2020 (*Pxyl-gfp-ftsZ*).

**Figure S9:** Localization of RpoC-GFP (*B. subtilis* 1048).

**Figure S10:** Localization of RpsB-GFP (*B. subtilis* 1049).

**Figure S11:** MinD translation induction assay.

**Figure S12:** Compound uptake after 60 min incubation with **Pt1**, **Pt8**, and **CisPt**.

**FigureS13:** ^1^H-NMR of P**t1** in DMSO-d_6_ at immediately after being dissolved (**A**), after incubated at 37°C for 24h (**B**), and 48h (**C**).

**FigureS14:** ^195^Pt-NMR of **Pt1** in DMSO-d_6_ at immediately after being dissolved (**A**), after incubated at 37°C for 24h (**B**), and 48h (**C**).

**Table S7:** Results from checkerboard assays combining **Pt1** with standard antibiotics.

**Table S8:** Antibiogram of *S. aureus* ATCC 29213 grown in the presence of sub-inhibitory concentration of **Pt-1**/levofloxacin for a period of 36 days.

**Table S9**: *In vitro* Post Antibiotic effect (PAE) of **Pt-1** at different concentrations.

**Figure S15**: Cell length measured from BCP images.

**Table S10**: *Bacillus subtilis* strains used in mode of action assays.

**Figure S16-54:** NMR spectra of Pt compounds

**Figure S55-66** MS spectra of Pt compounds

**S4. Supplementary references**

**S1) General Materials**

All reagents were available and used without further purification unless otherwise noted. Column chromatography was performed by using thick-walled glass columns and silica Gel 60 (Sigma Aldrich 230 – 400 mesh). Thin layer chromatography on pre-cut plates (Marcherey-Nagel, ALUGRAM Xtra Sil G/UV_254_), and visualization was provided by UV lamp at 254 nm and potassium permanganate stain. Argon was used as an inert gas. ^1^H NMR ,^13^C NMR and ^195^Pt spectra were recorded at room temperature on Bruker (operating at 300 MHz and 400 MHz for ^1^H NMR, 75 MHz for ^13^C NMR, and 86 MHz for^195^Pt NMR) in CDCl_3_, CD_2_Cl_2_-d*_2_*, DMSO and Acetone-d*_6_* with tetramethylsilane (TMS) as internal standard. Coupling constants *(J values*) are given in Hz and chemical shifts are reported as parts per million (ppm). Splitting patterns are designated as s (singlet), d (doublet), t (triplet), q (quartet), m (multiplet), and p (pentet). NMR spectra were processed with MestReNova. High resolution mass analyses were performed in positive ion mode with a ThermoScientific LTQ Orbitrap XL, fitted with an NSI ion source. The compounds were tested for their antimicrobial activity against *Escherichia coli* W3110, *Staphylococcus aureus* COL (MRSA) clinical isolate methicillin susceptible *Staphylococcus* *aureus* (MSSA)*,* and *Bacillus subtilis* MW168. The lead compounds were tested also against of *Escherichia coli* ATCC 25922, MSSA ATCC 29213; MRSA NRS 100, S119, 129, 186, 191, 192, 193, 194, 198, and vancomycin-resistant *Staphylococcus* *aureus (*VRSA*)* VRS 1, 4 and 12*.* Blood was obtained from Interregionale Blutspende SRK AG in Bern, Switzerland for haemolysis experiments. The HEK293T human epithelial cells were purchased from American Type Culture Collection (ATCC, Manassas, VA, USA). For cell viability, alamar Blue HS Cell Viability Reagent was purchased from Thermo Fisher Scientific. The relative proportions of solvents in chromatography solvent mixtures refer to the volume: volume ratio. The absorbance values were measured with a TECAN well plate reader instrument M1000. For MIC experiments, TPP tissue culture plates were used. MHC experiments were performed with Thermo Scientific Nunc™ 96-Well Polystyrene Conical Bottom MicroWell™ plates. The plots and statistical analysis were done with Prism. Microscopy experiments were performed on a Nikon Eclipse Ti2 equipped with a CFI Plan Apochromat DM Lambda 100x oil objective (N.A. 1.45, W.D. 0.13 mm), a Photometrics PRIME BSI camera, a Lumencor sola SE II FISH 365 light source, and an Okolab temperature incubation chamber. The microscopy images were processed with ImageJ^1^. Inductively coupled plasma mass spectrometry (ICP-MS) analysis was performed with a NexION 2000 ICP-MS (PerkinElmer).

**S2) Synthetic Procedures**

Synthesis of **1**^2^

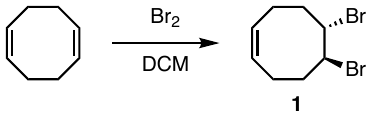

1,5-cyclooctadiene (1.00 g, 9.24 mmol) was dissolved in DCM (30 mL) and a bromine (0.200 mL, 3.85 mmol) solution in DCM (6 mL) was added dropwise over 30 minutes. The reaction progress was monitored by TLC. After it was stirred for another 30 minutes at room temperature, reaction mixture was concentrated *in vacuo*. Compound **1** was obtained through column chromatography (Heptane) as colourless oil (0.616 g, 77% yield). ^1^H NMR (CDCl_3_, 300 MHz) δ 5.64 (t, *J* = 4.1 Hz, 2H), 4.67 – 4.63 (m, 2H), 2.75 – 2.52 (m, 4H), 2.30 – 2.17 (m, 4H); ^13^C NMR (CDCl_3_, 75 MHz) δ 128.8, 59.6, 35.4, 25.0.

Synthesis of **2**^3^

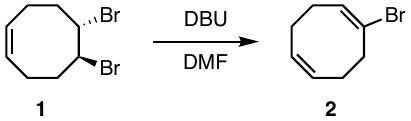

Compound **1** (1.13 g, 4.22 mmol) was dissolved in anhydrous DMF (12 mL) under inert conditions, and 1,8-Diazabicyclo(5.4.0)undec-7-ene(DBU) (0.674 g, 4.43 mmol) was added dropwise. The reaction mixture was stirred overnight at 80 °C. Reaction progress was monitored by TLC. Upon completion, 1M HCl (20 mL) was added and washed three times with EtOAc. The organic phase was dried over Na_2_SO_4_ and concentrated *in vacuo*. Compound **2** was obtained through column chromatography (Heptane) as a yellow oil (1.23 g, 21% yield). ^1^H NMR (300 MHz, CDCl_3_) δ 6.03 (p, *J* = 3.4 Hz, 1H), 5.64 – 5.47 (m, 2H), 2.82 (dd, *J* = 8.0, 5.3 Hz, 2H), 2.47 – 2.37 (m, 2H), 2.37 – 2.27 (m, 4H). ^13^C NMR (75 MHz, CDCl_3_) δ 130.3, 128.3, 127.9, 125.1, 39.0, 28.4, 27.5, 27.3.

Synthesis of **3**^4,5^

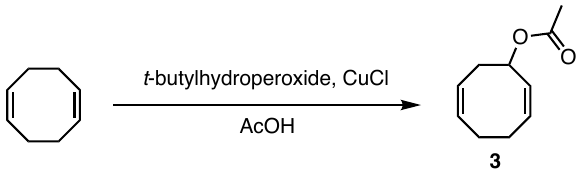

1,5-cyclooctadiene (5.29 g, 48.9 mmol) was dissolved in acetic acid (1.6 mL) and *t*-butyl hydroperoxide (1.87 g, 20.8 mmol), and a catalytic amount of CuCl was added. The reaction mixture was refluxed at 120 °C for 52 hours. Reaction progress was monitored by TLC. Upon completion, the reaction mixture was diluted with Et_2_O and washed twice with distilled water and one time with brine The organic phase was dried over Na_2_SO_4_ and concentrated *in vacuo*. Compound **3** was obtained through column chromatography (3:1 Heptane: EtOAc) as a yellow oil (2.95 g, 11 % yield). ^1^H NMR (300 MHz, CDCl_3_) δ 5.51 – 5.36 (m, 1H), 5.29 – 4.92 (m, 4H), 2.49 – 1.59 (m, 6H), 1.56 (dd, *J* = 5.6, 0.7 Hz, 3H); ^13^C NMR (75 MHz, CDCl_3_) δ 170.3, 129.6, 129.1, 129.0, 125.1, 72.3, 33.8, 27.9, 27.8, 21.2.

**General synthetic procedure for double bond-modified 1,5-cyclooctadiene ligands**

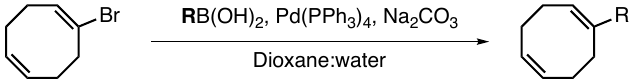

Compound **2** (0.200 g, 1.06 mmol) was suspended in 1,4-dioxane (3 mL) and water (1.5 mL). Boronic acid derivative (2.5 eq), tetrakis-(triphenylphosphine)-palladium (0.03 eq.), and Na_2_CO_3_(3 eq.) were added to the reaction mixture. The reaction mixture was refluxed for 5 hours at 110 °C. Reaction progress was monitored by TLC, and upon completion, the reaction mixture was diluted with EtOAc. The organic phase was separated, and the aqueous phase was washed three times with EtOAc. The combined organic phase was washed with brine. The combined organic phases were dried over Na_2_SO_4_ and concentrated *in vacuo*. The desired products were obtained through column chromatography (Heptane:EtoAc).

Compound **4**

Colorless oil, 0.0404 g, 36% yield. ^1^H NMR (300 MHz, CDCl_3_) δ 7.22 – 7.08 (m, 5H), 5.78 (t, *J* = 6.5 Hz, 1H), 5.51 (t, *J* = 6.0 Hz, 2H), 2.76 – 2.70 (m, 2H), 2.41 (s, 6H); ^13^C NMR (75 MHz, CD_2_Cl_2_) δ 145.4, 140.1, 128.8, 128.5, 128.1, 127.4, 127.3, 127.1, 126.4, 126.2, 31.2, 28.2, 28.2, 28.1.

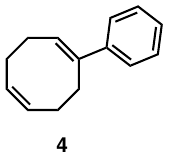

### Compound 5

Pale yellow oil, (0.077 g, 41% yield). ^1^H NMR (300 MHz, CDCl_3_) δ 7.09 (t, *J* = 7.8 Hz, 1H), 6.79 – 6.73 (m, 1H), 6.67 (t, *J* = 1.9 Hz, 1H), 6.56 (ddd, *J* = 7.8, 2.4, 1.0 Hz, 1H), 5.86 (t, *J* = 6.4 Hz, 1H), 5.68 – 5.56 (m, 2H), 3.60 (*br*. s, 2H), 2.83 – 2.76 (m, 2H), 2.57 – 2.40 (m, 6H); ^13^C NMR (75 MHz, CDCl_3_) δ 146.6, 146.2, 140.2, 129.0, 128.7, 128.6, 126.9, 116.9, 113.4, 31.2, 28.3, 28.1, 28.1; HRMS (ES+): m/z calculated for C_14_H_18_O^+^ 200.1434: found 200.1434 [M+H]^+^.

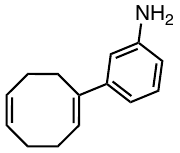

### Compound 6

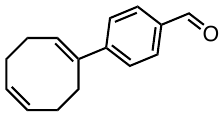

Yellow oil, 0.225 g, 41% yield. ^1^H NMR (300 MHz, CDCl_3_) ^1^H NMR (300 MHz, CDCl_3_) δ 9.94 (s, 1H), 7.76 (d, *J* = 8.6 Hz, 2H), 7.44 (d, *J* = 8.3 Hz, 2H), 5.98 (t, *J* = 6.6 Hz, 1H), 5.57 (p, *J* = 2.9 Hz, 2H), 2.80 (t, *J* = 6.8 Hz, 2H), 2.57 – 2.38 (m, 6H); ^13^C NMR (75 MHz, CDCl_3_) δ 192.0, 151.5, 139.3, 134.6, 130.3, 129.8, 128.8, 128.4, 126.7, 117.5, 30.8, 28.3, 28.2, 27.8. HRMS (ES+): m/z calculated for C_15_H_17_O^+^ 213.1279: found 213.1272 [M+H]^+^.

### Compound 7

Pale yellow oil, 0.110 g, 24% yield. ^1^H NMR (300 MHz, CDCl_3_) δ 7.33 (d, *J* = 2.6 Hz, 1H), 6.39 – 6.30 (m, 2H), 6.20 (d, *J* = 3.5 Hz, 1H), 5.65 – 5.58 (m, 2H), 2.80 – 2.74 (m, 2H), 2.62 – 2.47 (m, 6H). ^13^C NMR (75 MHz, CDCl_3_) δ 156.3, 141.1, 129.4, 128.7, 128.5, 124.2, 111.0, 103.7, 28.20, 27.9. HRGCMS: t_R_: 10.7 min: m/z calculated for C_12_H_14_O 174.1045; found 174.1042 [M].

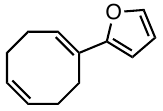

**General synthetic procedure for Pt(II) complexes with double bond-modified 1,5-cyclooctadiene ligands**^6^

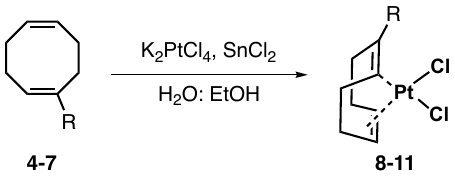

K_2_PtCl_4_ (0.100 g, 0.240 mmol) was suspended in water (3 mL) and ethanol (3 mL) and COD ligand (0.960 mmol, 4 eq.) was added. Then, catalytic amount of SnCl_2_ was added to the reaction mixture and stirred for 2 days at room temperature. Upon precipitation, the reaction mixture was concentrated *in vacuo* and desired products were obtained through column chromatography (Heptane: EtOAc).

Compound **8**

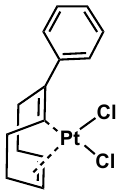

Pale yellow solid, (0.052 g, 52% yield). ^1^H NMR (400 MHz, CDCl_3_) δ 7.53 (d, *J* = 7.2 Hz, 2H), 7.37 (dd, *J* = 22.8, 11.3 Hz, 3H), 6.25 – 6.04 (m, 1H), 5.73 (d, *J* = 45.0 Hz, 2H), 3.23 – 3.11 (m, 1H), 3.03 (m, 1H), 2.92 – 2.84 (m, 1H), 2.73 – 2.61 (m, 2H), 2.56 – 2.35 (m, 2H), 2.15 – 2.02 (m, 1H); ^13^C NMR (101 MHz, CDCl_3_) δ 141.3, 130.1, 128.4, 127.7, 120.9, 100.4, 98.3, 91.9, 38.1, 33.5, 32.8, 29.4; ^195^Pt NMR (86 MHz, DMSO) δ -3187.27; HRMS (ES+): m/z calculated for C_14_H_20_Cl_2_NPt^+^ 467.0615: found 467.0610 [M+NH_4_]^+^.

Compound **9**

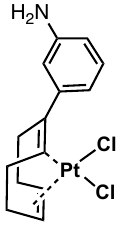

Brown solid, (0.0099 g, 22% yield). ^1^H NMR (300 MHz, Acetone) δ 7.42 (s, 1H), 7.21 – 7.09 (m, 3H), 6.40 (s, 1H), 5.89 (t, *J* = 6.5 Hz, 1H), 5.55 (t, *J* = 3.7 Hz, 2H), 2.78 (dd, *J* = 7.6, 6.1 Hz, 2H), 2.47 (tt, *J* = 12.7, 5.2 Hz, 6H).HRMS (ES+): m/z calculated for C_14_H_17_ClNPt^+^ 429.0697: found 429.0690 [M-Cl]^+^.

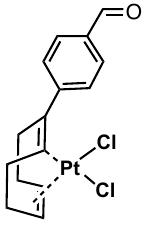
Compound **10**

Yellow solid, (0.042 g, 33% yield). ^1^H NMR (400 MHz, CDCl_3_) δ 9.99 (s, 1H), 7.82 (d, *J* = 8.5 Hz, 2H), 7.68 (d, *J* = 8.2 Hz, 2H), 6.17 (d, *J* = 9.7 Hz, 1H), 5.87 – 5.71 (m, 2H), 3.26 – 3.13 (m, 1H), 3.05 (m, 1H), 2.84 – 2.75 (m, 1H), 2.70 – 2.60 (m, 2H), 2.58 – 2.49 (m, 2H), 2.45 – 2.36 (m, 2H).^13^C NMR (101 MHz, CDCl_3_) δ 191.6, 147.4, 136.7, 129.7, 128.5, 116.9, 101.6, 99.6, 93.7, 37.8, 33.5, 33.0, 29.2;^195^Pt NMR (86 MHz, CDCl_3_) δ -3190.91. HRMS (ES+): m/z calculated for C_15_H_20_Cl_2_NOPt^+^ 495.0564: found 495.0579 [M+NH_4_]^+^.

Compound **11**

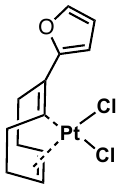

Yellow solid, (0.034 g, 29% yield). ^1^H NMR (300 MHz, CD_2_Cl_2_) δ 7.61 (dd, *J* = 1.8, 0.8 Hz, 1H), 6.79 (dd, *J* = 3.6, 0.8 Hz, 1H), 6.46 (dd, *J* = 3.5, 1.8 Hz, 1H), 5.91 (dd, *J* = 7.4, 3.4 Hz, 1H), 5.68 (td, *J* = 7.7, 4.3 Hz, 1H), 5.57 (td, *J* = 7.3, 3.1 Hz, 1H), 3.12 (ddd, *J* = 14.5, 9.2, 4.8 Hz, 1H), 3.03 – 2.78 (m, 3H), 2.78 – 2.63 (m, 2H), 2.53 – 2.39 (m, 2H);^13^C NMR (101 MHz, CD_2_Cl_2_) δ 151.6, 145.7, 114.1, 113.9, 112.9, 101.4, 98.1, 87.5, 35.5, 31.7, 31.5, 30.6; ^195^Pt NMR (86 MHz, Acetone) δ -3151.48. HRMS (ES+): m/z calculated for C_12_H_14_ClOPt^+^ 404.0381: found 404.0373 [M-Cl]^+^.

**General synthetic procedure for Pt(II) complexes with allylic modified 1,5-cyclooctadiene ligands**

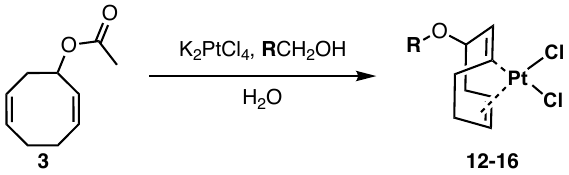

K_2_PtCl_4_ (1 eq.) was suspended in water (1.5 mL) and a primary alcohol (20 eq.) and compound **3** (5 eq.) was added. The reaction mixture was heated by microwave to 90°C for 75 minutes (low absorbance setting). Upon completion, the reaction mixture was concentrated *in vacuo* and the desired product was obtained through column chromatography (Heptane: EtOAc).

Compound **12**

Light brown solid, (0.009 g, 9% yield). ^1^H NMR (400 MHz, CDCl_3_) δ 5.77 – 5.64 (m, 2H), 5.55 – 5.46 (m, 1H), 5.25 – 5.10 (m, 1H), 4.65 (td, *J* = 8.7, 6.0 Hz, 1H), 3.60 (qd, *J* = 7.0, 3.4 Hz, 2H), 3.01 – 2.82 (m, 2H), 2.57 (ddt, *J* = 20.5, 11.4, 5.2 Hz, 2H), 2.29 (ddd, *J* = 21.4, 14.1, 6.5 Hz, 2H), 1.19 (s, 3H); ^13^C NMR (101 MHz, CDCl_3_) δ 103.5, 99.5, 94.2, 88.8, 78.9, 76.3, 76.0, 75.7, 65.5, 32.8, 31.5, 27.7, 14.5; ^195^Pt NMR (86 MHz, CDCl_3_) δ -3346.0; HRMS (ES+): m/z calculated for C_10_H_20_Cl_2_NOPt^+^ 435.0570: found 435.0573 [M+NH_4_]^+^.

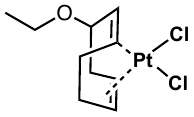

Compound **13**

White solid, (0.014 g, 13% yield). ^1^H NMR (300 MHz, CDCl_3_) δ 5.77 – 5.63 (m, 2H), 5.51 (td, *J* = 8.2, 5.9 Hz, 1H), 5.21 (tdd, *J* = 8.0, 6.5, 1.2 Hz, 1H), 4.64 (td, *J* = 8.5, 5.9 Hz, 1H), 3.49 (td, *J* = 6.6, 1.9 Hz, 2H), 3.05 – 2.79 (m, 2H), 2.68 – 2.49 (m, 2H), 2.30 (tt, *J* = 14.1, 6.6 Hz, 2H), 1.61 (m, 2H), 0.92 (t, *J* = 7.4 Hz, 3H). ^13^C NMR (101 MHz, CDCl_3_) δ 104.4, 100.5, 95.2, 89.9, 80.1, 72.7, 33.9, 32.4, 28.8, 23.2, 10.5; ^195^Pt NMR (86 MHz, CDCl_3_) δ -3343.2; HRMS (ES+): m/z calculated for C_11_H_18_Cl_2_KOPt^+^ 470.0014: found 470.0011 [M+K]^+^.

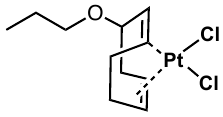

Compound **14**

Dark brown solid, (0.036 g, 10% yield). ^1^H NMR (400 MHz, CDCl_3_) δ 5.76 (ddd, *J* = 16.6, 8.2, 3.3 Hz, 4H), 4.16 – 4.02 (m, 1H), 3.96 – 3.81 (m, 2H), 3.50 (t, *J* = 5.9 Hz, 2H), 2.79 – 2.66 (m, 2H), 2.61 – 2.46 (m, 2H), 2.18 – 2.08 (m, 1H), 1.90 – 1.79 (m, 1H); ^13^C NMR (101 MHz, CDCl_3_) δ 101.4, 101.0, 97.6, 97.5, 81.0, 69.8, 39.8, 32.5, 30.6, 30.1; ^195^Pt NMR (86 MHz, CDCl_3_) δ -3638.5.

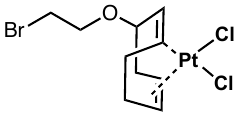

Compound **15**

Yellow solid, (0.014 g, 8% yield). ^1^H NMR (400 MHz, CDCl_3_) δ 7.41 – 7.29 (m, 5H), 5.77 – 5.60 (m, 2H), 5.49 (td, *J* = 8.2, 5.9 Hz, 1H), 5.17 (q, *J* = 7.3 Hz, 1H), 4.72 (td, *J* = 8.8, 6.0 Hz, 1H), 4.61 (s, 2H), 2.97 – 2.81 (m, 2H), 2.66 – 2.51 (m, 2H), 2.40 – 2.25 (m, 2H). ^13^C NMR (101 MHz, CDCl_3_) δ 137.03, 128.76, 128.5, 127.8, 104.7, 100.5, 94.8, 89.5, 79.3, 76.7, 72.9, 33.7, 32.6, 28.7;^195^Pt NMR (86 MHz, CDCl_3_) δ -3342.0. HRMS (ES+): m/z calculated for C_15_H_22_Cl_2_NOPt^+^ 497.0721: found 497.0717 [M+NH_4_]^+^.

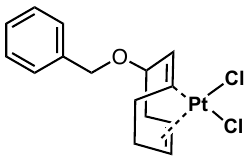

Compound **16**

Yellow solid, (0.022 g, 12% yield). ^1^H NMR (400 MHz, CDCl_3_) δ 7.31 (d, *J* = 8.2 Hz, 2H), 7.18 – 7.13 (m, 2H), 5.66 – 5.58 (m, 1H), 5.57 – 5.48 (m, 1H), 5.47 – 5.35 (m, 1H), 5.12 – 4.95 (m, 1H), 4.66 – 4.56 (m, 1H), 4.48 (s, 1H), 2.88 – 2.73 (m, 2H), 2.55 – 2.41 (m, 2H), 2.22 (ddd, *J* = 21.0, 11.8, 7.3 Hz, 3H), 1.23 (s, 9H); ^13^C NMR (101 MHz, CDCl_3_) δ 151.6, 133.9, 127.8, 126.9, 125.7, 104.7, 100.6, 95.0, 89.6, 79.0, 72.6, 34.7, 33.6, 32.6, 31.3, 28.7; ^195^Pt NMR (86 MHz, CDCl_3_) δ -3343.4. HRMS (ES+): m/z calculated for C_19_H_30_Cl_2_NOPt^+^ 553.1347: found 553.1341 [M+NH_4_]^+^.

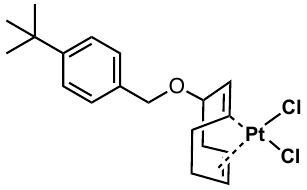

**S3) Supplementary Tables and Figures**

**TableS1:** Antibacterial activity of synthesized PtCOD complexes against *E. coli*.

|  | ***E. coli*** |
| --- | --- |
| **Pt1** | >100 |
| **Pt2** | >100 |
| **Pt3** | >100 |
| **Pt4** | >100 |
| **Pt5** | >100 |
| **Pt6** | >100 |
| **Pt7** | >100 |
| **Pt8** | >100 |
| **Pt9** | >100 |
| **Pt10** | >100 |

**Table S2:** Antibacterial activity of the Pt-drugs against Gram-positive strains.

|  | **Minimum Inhibitory Concentration (MIC) [µM]** | | | |
| --- | --- | --- | --- | --- |
|  | **MRSA** | **MSSA** | ***B. subtilis*** | ***E. coli*** |
| **Carboplatin** | >100 | >100 | >100 | >100 |
| **Oxaliplatin** | >100 | >100 | >100 | >100 |

|  | **Minimum Inhibitory Concentration (MIC) [µM]** | |
| --- | --- | --- |
|  | ***S. aureus*** | ***E. coli*** |
| **1,5-Cyclooctadiene** | >200 | >200 |
| **K_2_PtCl_4_** | >200 | >200 |

**Table S3:** Antibacterial activity of COD ligand and Pt precursor against *S. aureus* (CCUG 62707) and *E. coli* (NCTC 13476).

**Table S4:** Antibacterial activity of the lead complexes in µg/mL against a selection of Gram-positive and Gram-negative strains.

| Minimum Inhibitory Concentration (MIC) [µg/mL] | | | | |
| --- | --- | --- | --- | --- |
|  |  | | **Pt1** | **Pt8** |
| MSSA | **ATCC 29213** | | 0.25 | 2 |
| MRSA | **NRS 100** | | 0.25 – 0.5 | 4 – 8 |
|  | **NRS 119** | | 0.125 – 0.25 | 1 |
|  | **NRS 129** | | 0.125 – 0.25 | 2 – 1 |
|  | **NRS 186** | | 0.125 – 0.25 | 2 – 8 |
|  | **NRS 191** | | 0.25 | 1 – 2 |
|  | **NRS 192** | | 0.125 – 0.25 | 1 |
|  | **NRS 193** | | 0.25 | 2 |
|  | **NRS 194** | | 0.125 – 0.25 | 1 |
|  | **NRS 198** | | 0.25 | 4 |
| VRSA | **VRS 1** | | 0.25 – 0.5 | 32 |
|  | **VRS 4** | | 0.25 – 0.5 | 4 |
|  | **VRS 12** | | 0.25 – 0.5 | 4 – 8 |
| *E. coli* | **ATCC 25922** | | >64 | >64 |
| *K. pneumoniae* | **BAA 1705** | | >64 | >64 |
| A. *baumanii* | **BAA 1605** | | >64 | >64 |
| *P. aeruginosa* | **ATCC 27853** | | >64 | >64 |

**Table S5.** HC_50_ values of synthesised Pt complexes against hRBC.

|  | **HC_50_ (µM)** |
| --- | --- |
| **Pt1** | >200 |
| **Pt2** | >200 |
| **Pt3** | >200 |
| **Pt4** | >200 |
| **Pt5** | >200 |
| **Pt6** | >200 |
| **Pt7** | >200 |
| **Pt8** | >200 |
| **Pt9** | >200 |
| **Pt10** | >200 |

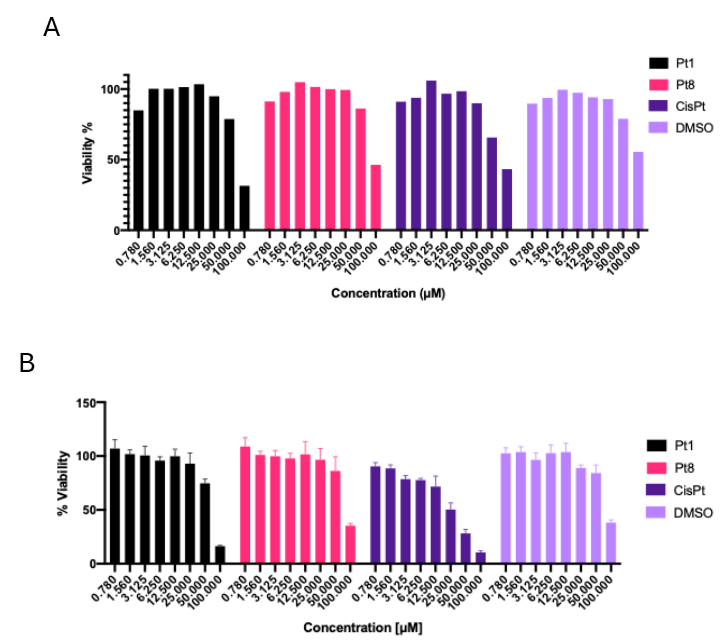

**Figure S1:** Growth-normalized % viability of HEK293T cells after treatment with **Pt1**, **Pt8**, **CisPt,** and DMSO (negative control) after incubation for 24 h (**A**) and 48 h (**B**).

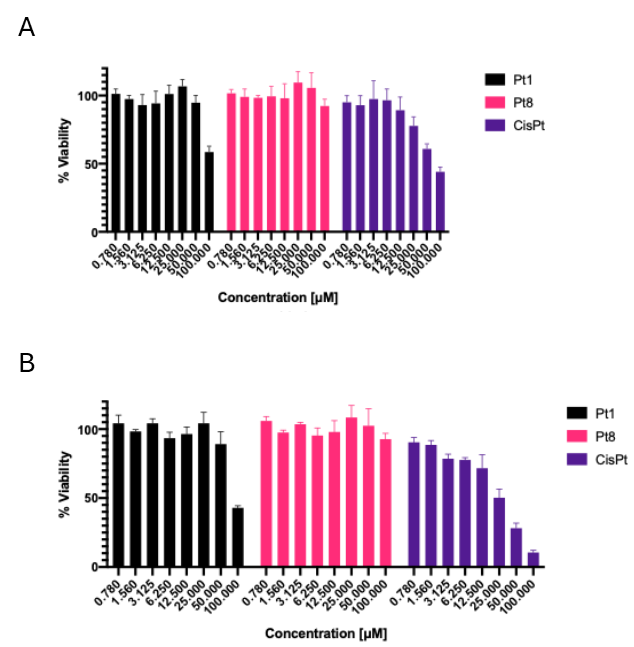

**Figure S2:** DMSO-normalized % viability of HEK293T cells after treatment with **Pt1**, **Pt8**, and **CisPt** after incubation for 24 h (**A**) and 48 h (**B)**. For solubility reasons, the Pt lead compounds were dissolved in 100% DMSO. This resulted in final DMSO concentrations above the acceptable range (0.5-1%) in samples with the highest tested compound concentrations. To account for this the toxic effect of DMSO, data was normalized to the corresponding DMSO controls.

**Table S6:** DMSO-normalized cytotoxicity (IC_50_ in µM) of **Pt1** and **Pt8** against HEK293T cells.

|  | **HEK293T** | |
| --- | --- | --- |
|  | **24h** | **48h** |
| **Pt1** | >100 | 92.9 ± 1.8 |
| **Pt8** | >100 | >100 |

| **A**  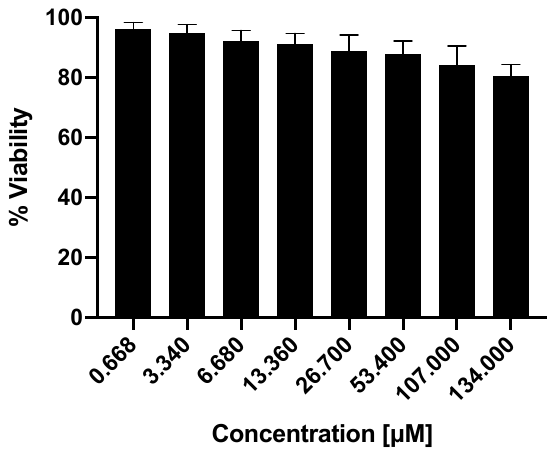 | **B**  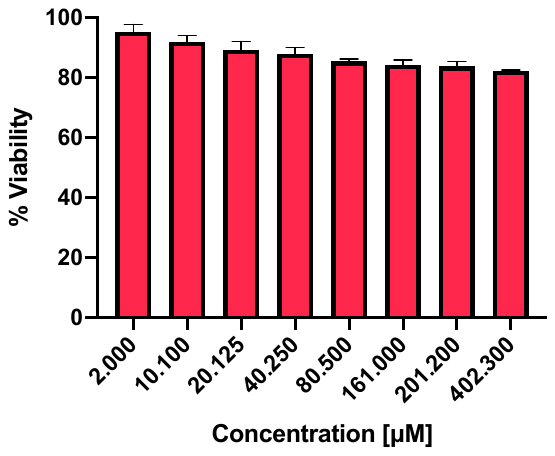 |
| --- | --- |

**Figure S3:** Percent viability of Vero cells upon treatment with **Pt1** (**A**) and **Pt8** (**B**) after incubation for 72 h.

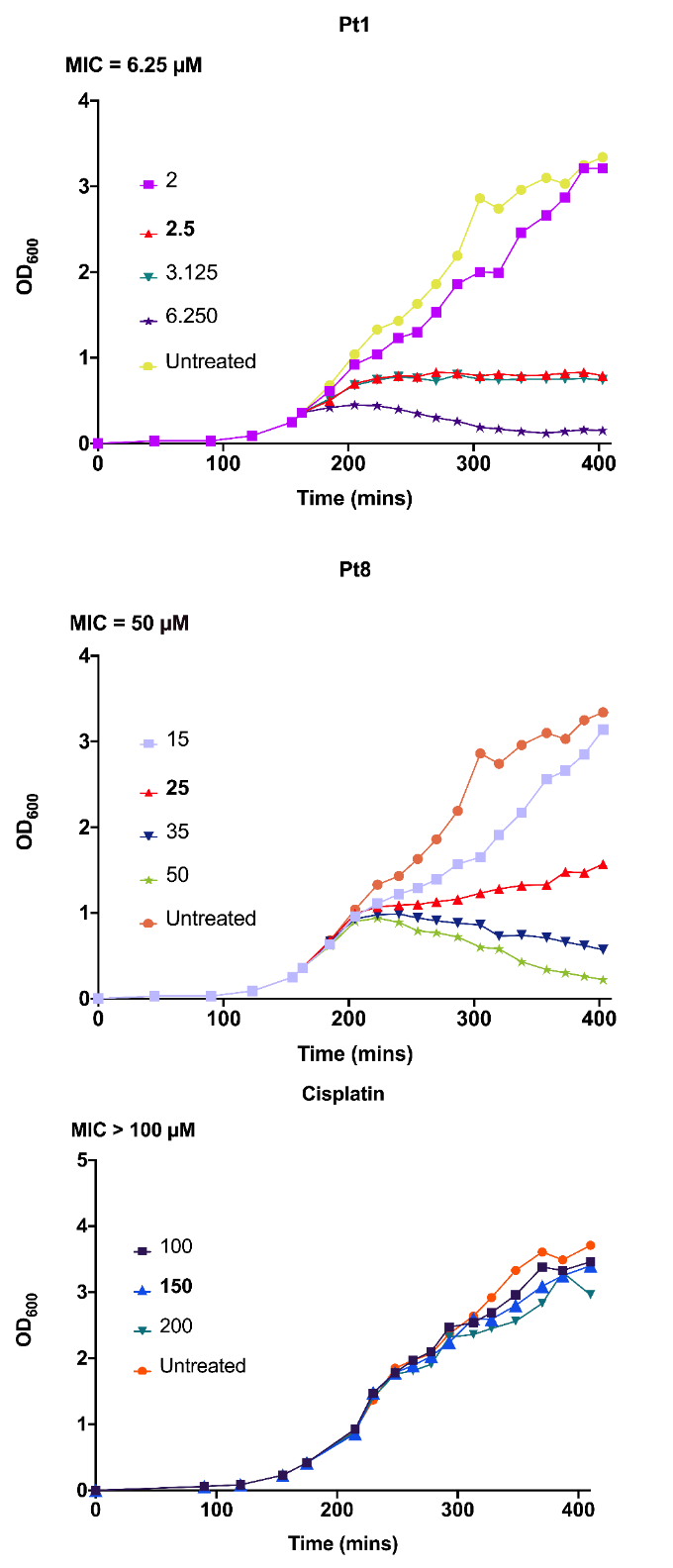

**Figure S4:** Determination of optimal stressor concentrations (indicated in bold) of **Pt1**, **Pt8**, and **CisPt**.

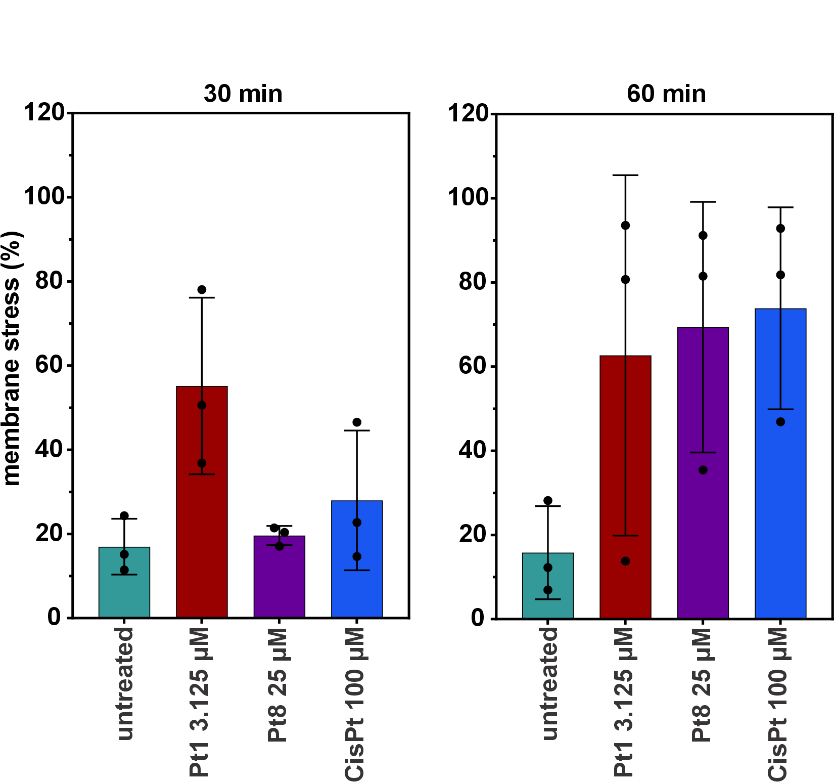

**Figure S5:** Quantification of membrane stress measured as percent of *B. subtilis* bSS82 cells with Nile red foci. Error bars show standard deviation of three biological replicates. No statistically significant changes were observed (p≤0.05). For statistical analysis, the Mann-Whitney (unpaired, nonparametric) test was used.

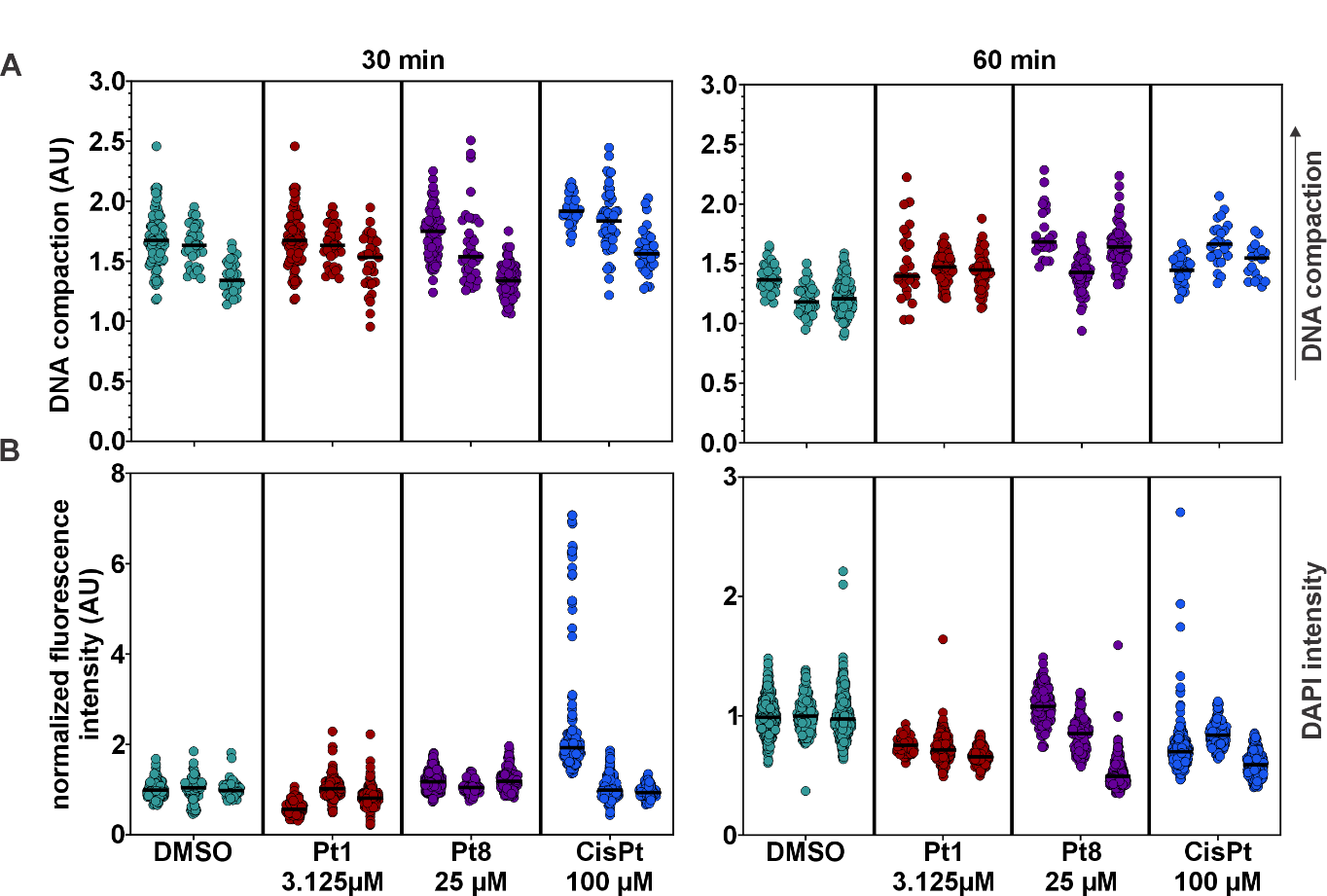

**Figure S6:** Quantification of nucleoid compaction (**A**) and DAPI fluorescence intensity (**B**) corresponding to **Figure 2**, separated into individual biological replicates.

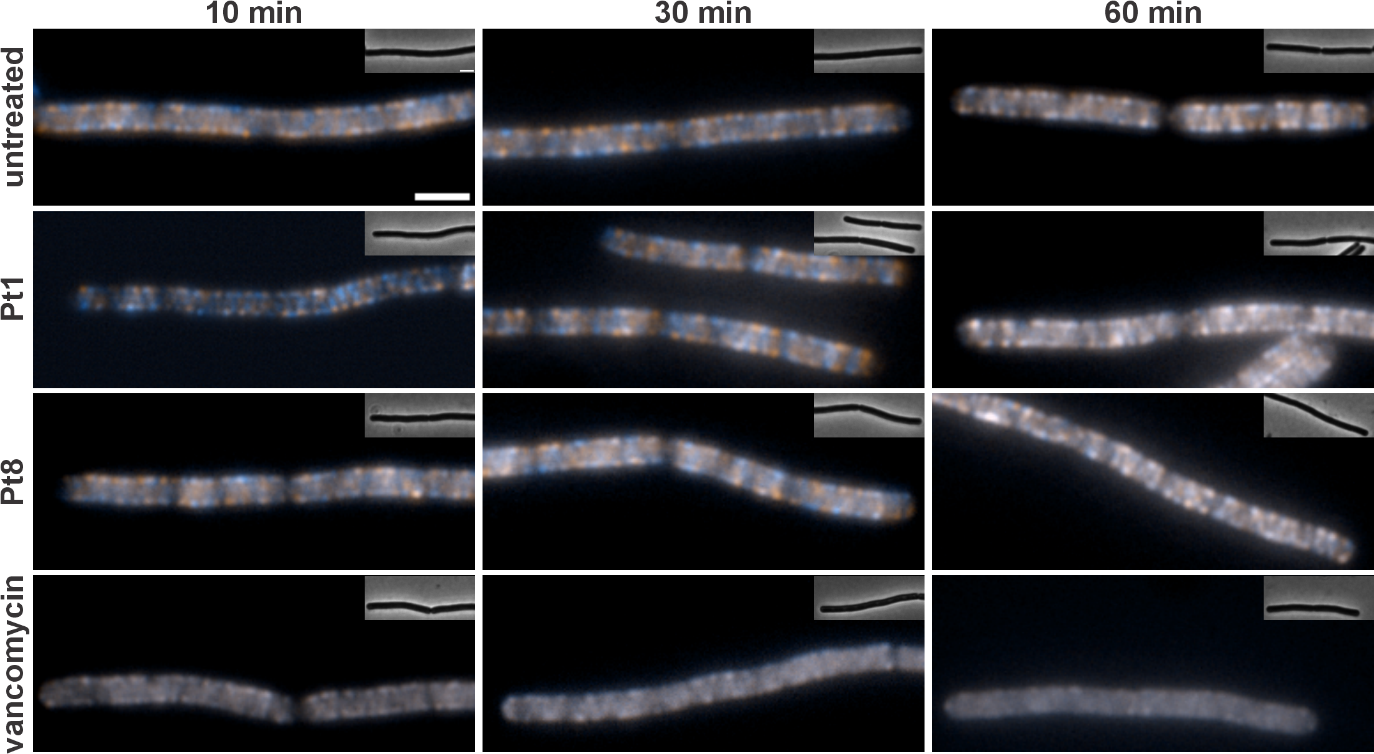

**Figure S7:** Fluorescence overlays of *B. subtilis* MW10 (*Pxyl-gfp-mreB*). Cultures were treated with 3.12 µM **Pt1**, 25 µM **Pt8**, or 0.67 µM **vancomycin** for 30 and 60 min. Images show overlays of two consecutive images captured from the same cells in a 30-second interval. Images were false-colored in blue and orange with overlapping signal appearing white. Distinct blue and orange spots indicate movement of MreB foci. Entirely white images indicate a stop of MreB moility (vancomycin control). Scale bar 2 µm.

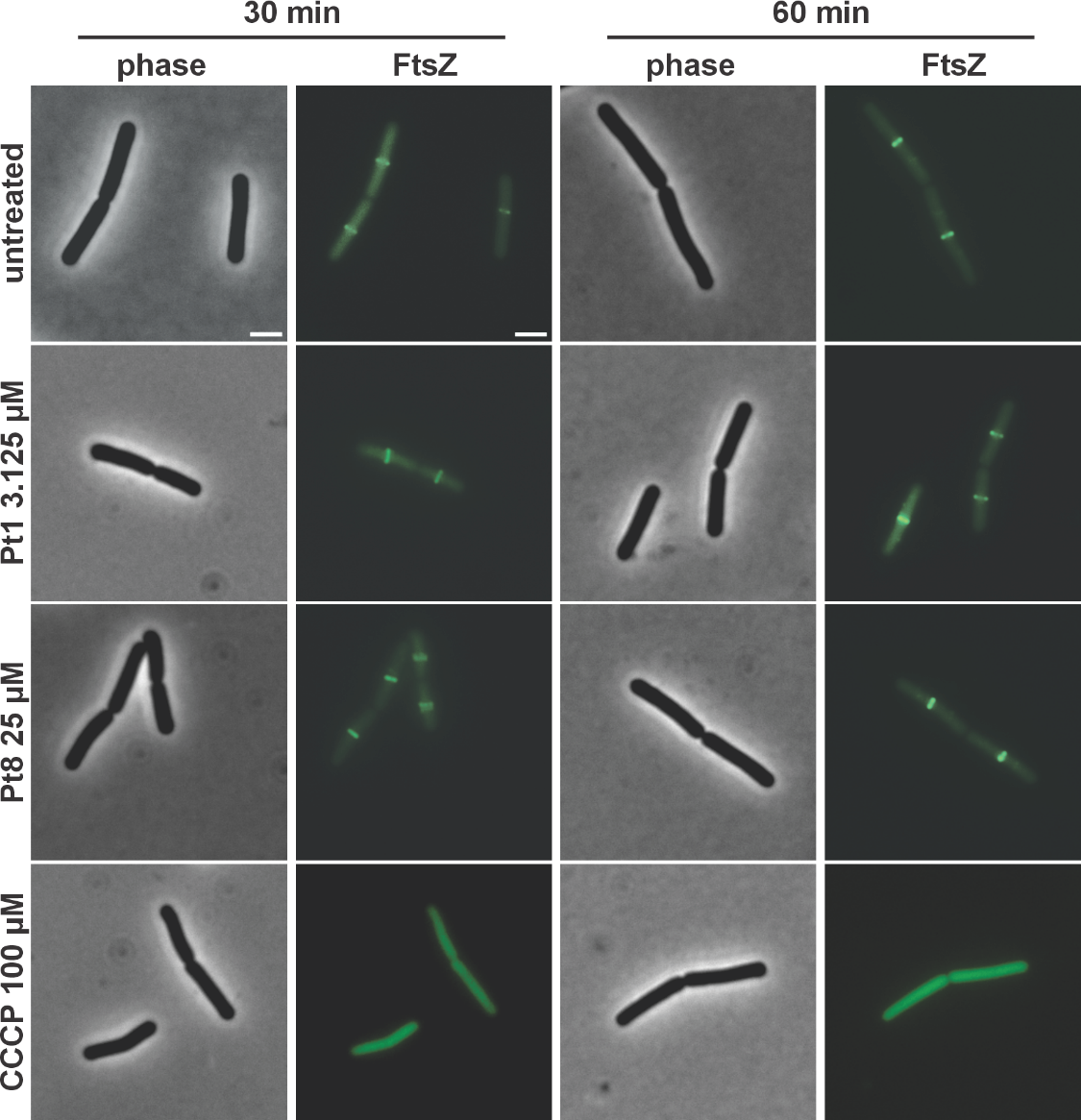

**Figure S8:** Localization of FtsZ in *B. subtilis* 2020 (*Pxyl-gfp-ftsZ*). Cultures were treated with 3.12 µM **Pt1**, 25 µM **Pt8**, or 100 µM **CCCP** for 30 and 60 min. Scale bar 2 µm.

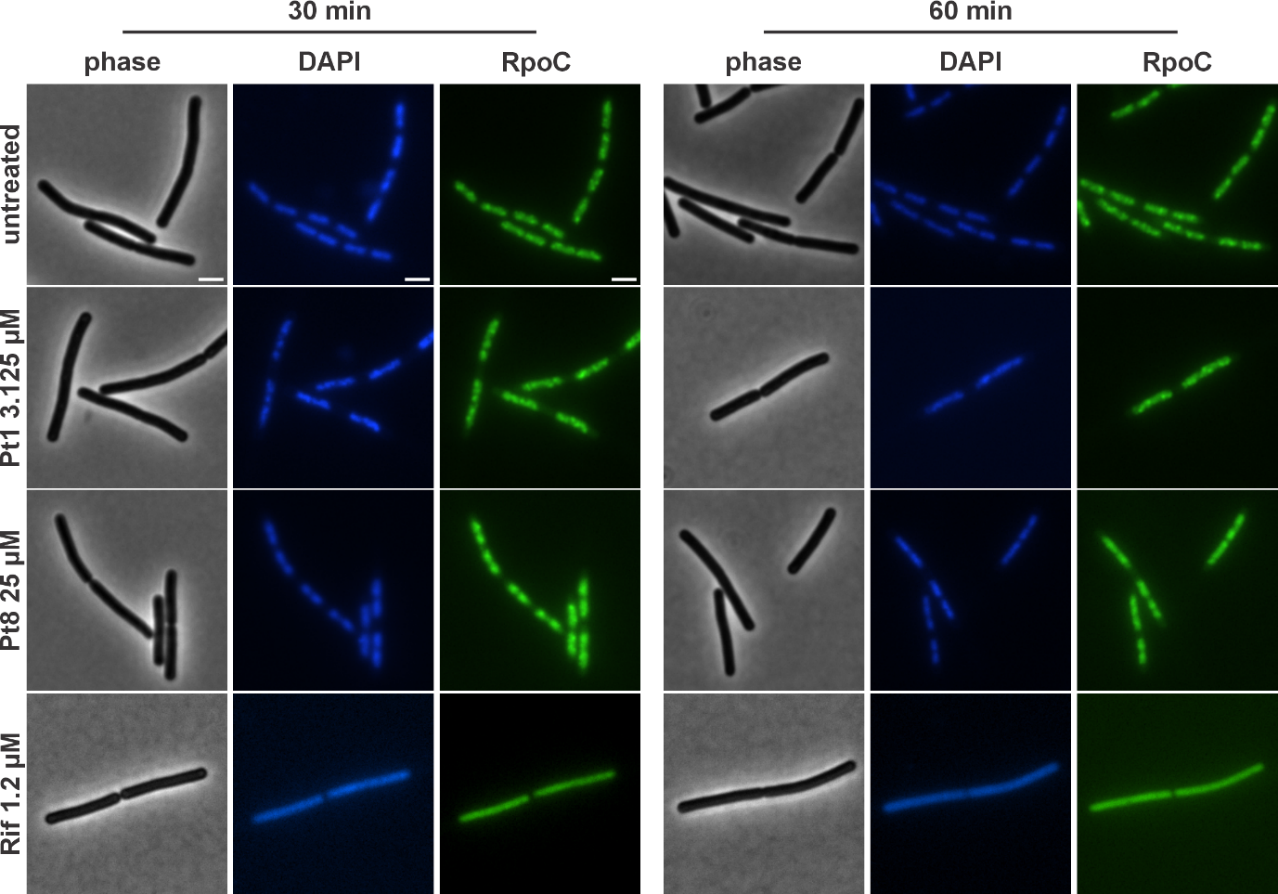

**Figure S9:** Localization of RpoC-GFP (*B. subtilis* 1048). Cultures were treated with 3.12 µM **Pt1**, 25 µM **Pt8**, or 1.2 µM rifampicin (**RIF**) for 30 and 60 min. Scale bar 2 µm.

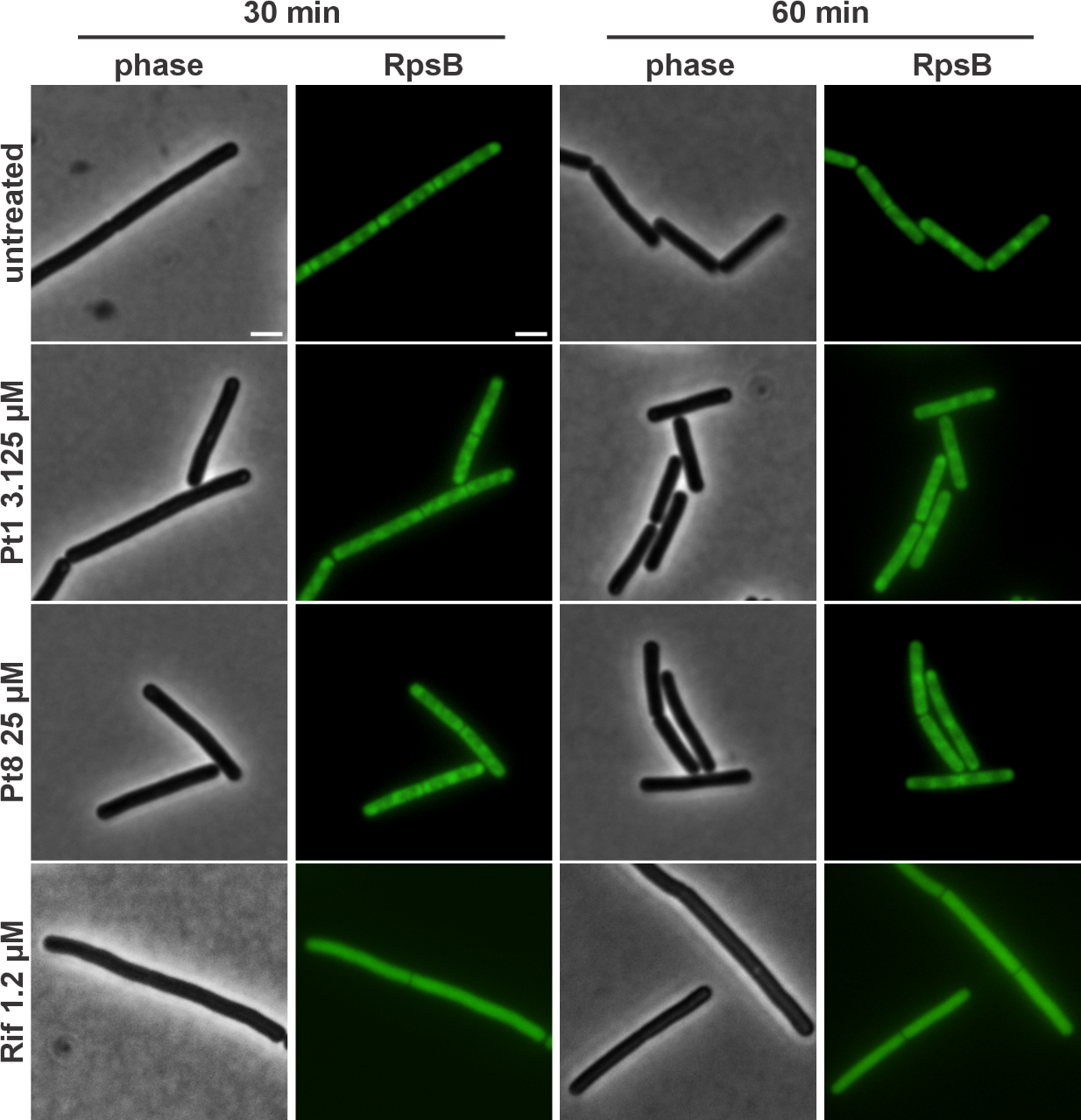

**Figure S10:** Localization of RpsB-GFP (*B. subtilis* 1049). Cultures were treated with 3.12 µM **Pt1**, 25 µM **Pt8**, or 1.2 µM rifampicin (**Rif**) for 30 and 60 min. Scale bar 2 µm.

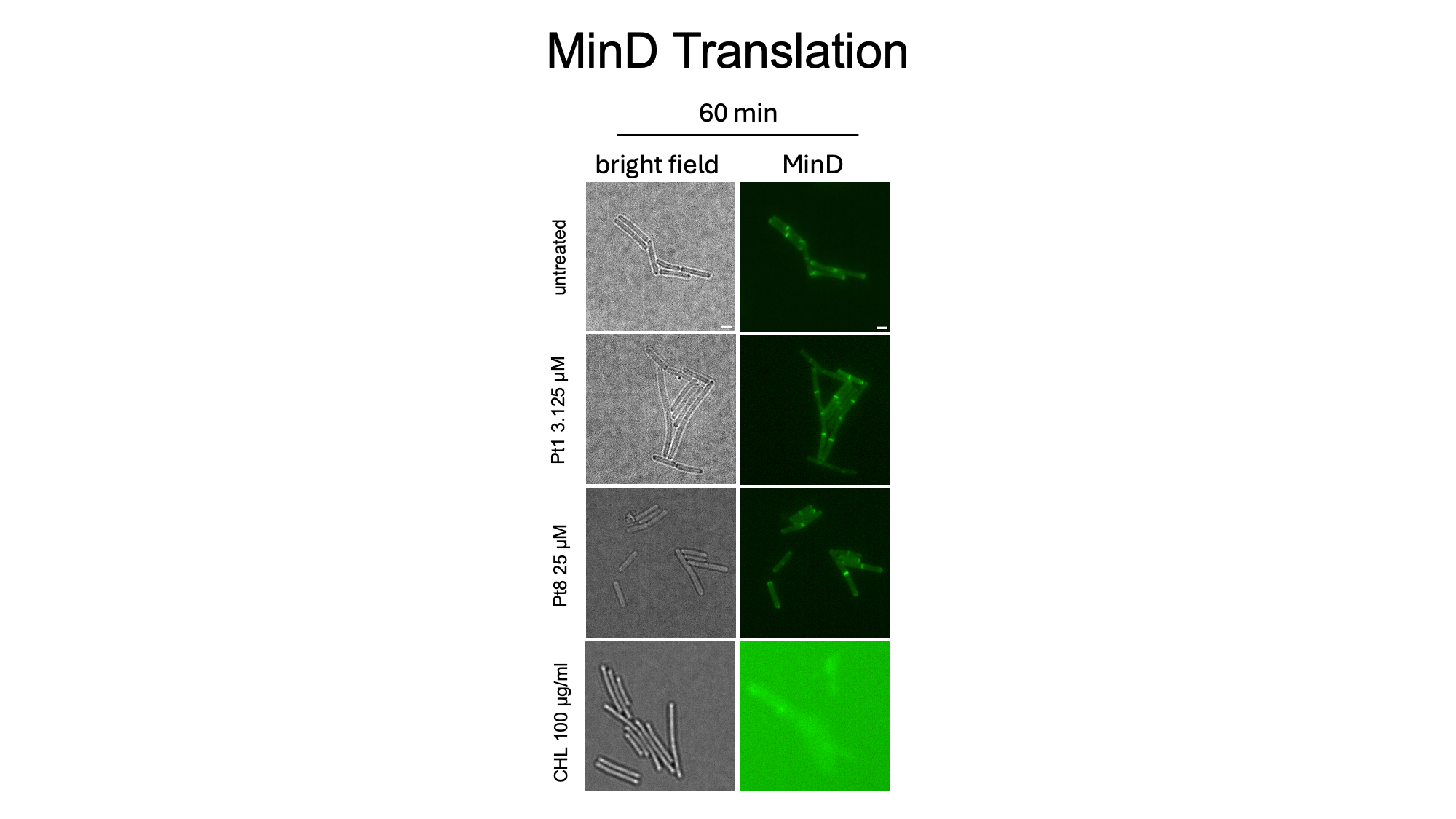

**Figure S11:** MinD translation induction assay. *B. subtilis* LH131 (*Pxyl-gfp-minD*) was grown without xylose, thus not expressing GFP-MinD. In log phase, cells were simultaneously induced with 0.1% xylose and treated with 3.125 µM **Pt1**, 25 µM **Pt8**, or 309.5 µM (100 µg/ml) chloramphenicol (**CHL**). Images were taken after 60 min of induction/treatment. Scale bar 2 µm.

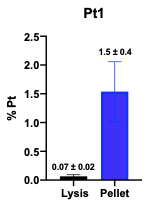

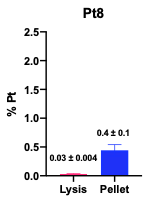

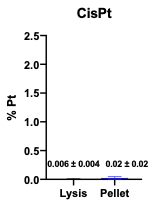

**Figure S12:** Compound uptake after 60 min incubation with **Pt1**, **Pt8**, and **CisPt**. Pt content was measured with ICP-MS.

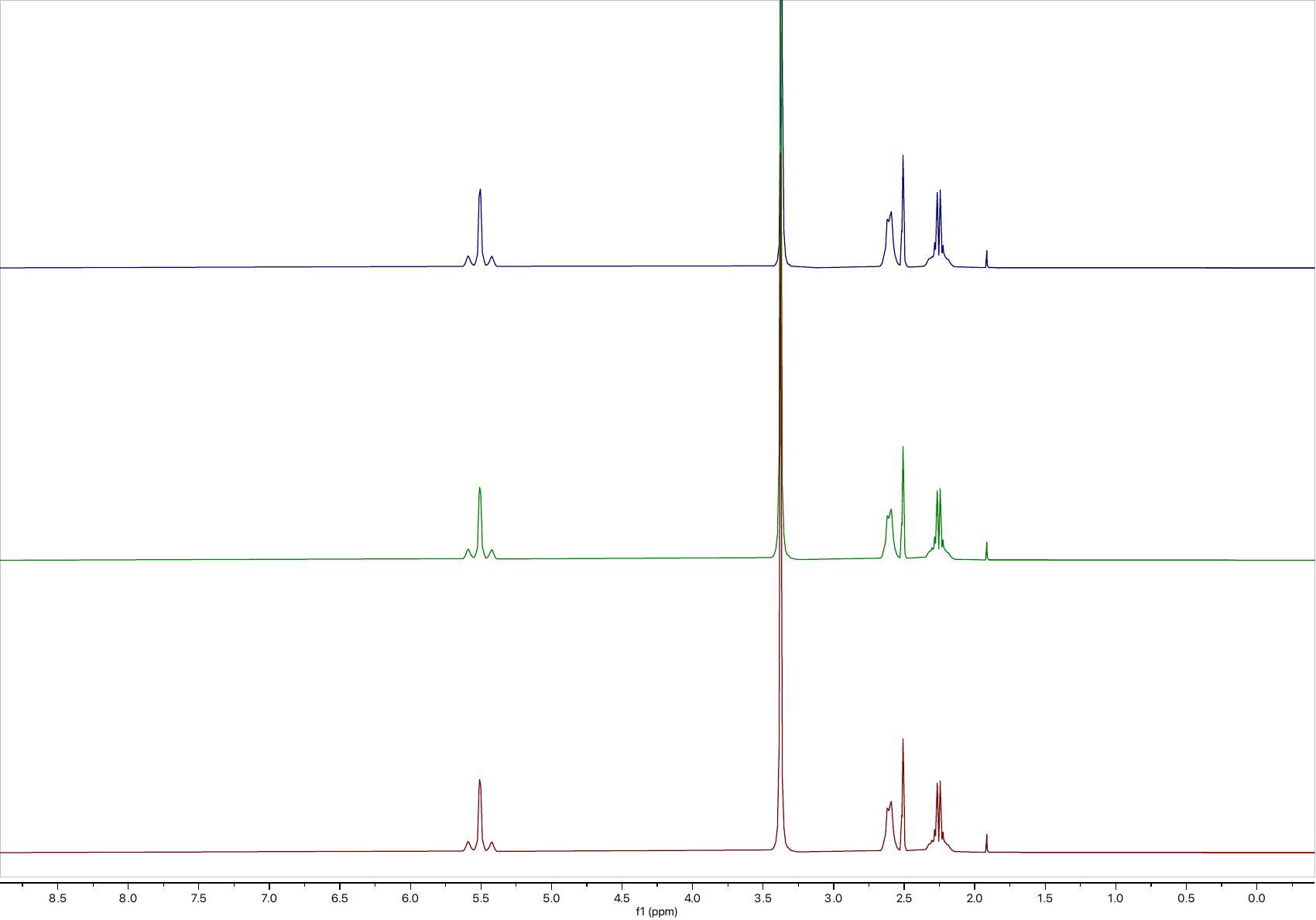

**C**

**B**

**A**

**FigureS13:** ^1^H-NMR of **Pt1** in DMSO-*d*_6_ at immediately after being dissolved (**A**), after incubated at 37°C for 24 h (**B**), and 48 h (**C**).

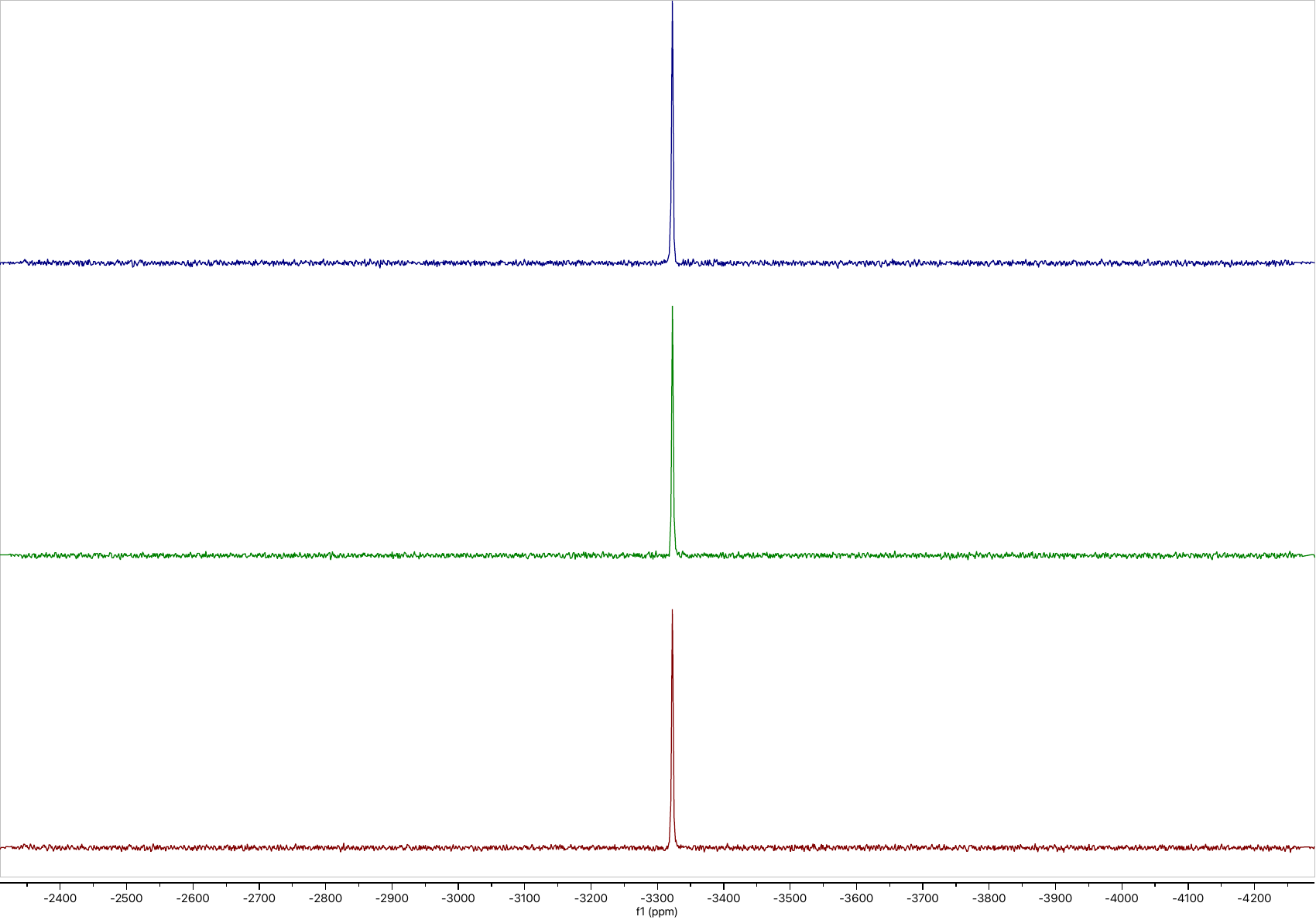

**B**

**C**

**A**

**Figure S14:** ^195^Pt-NMR of **Pt1** in DMSO-*d*_6_ at immediately after being dissolved (**A**), after incubated at 37°C for 24 h (**B**), and 48 h (**C**).

**Table S7:** Results from checkerboard assays combining **Pt1** with standard antibiotics. Synergy was tested against *S. aureus* ATCC 29213. NI: no interaction

| **Antibiotic** | **MIC (µg/mL)** | **MIC of Pt1 in the presence of antibiotic (µg/mL)**  **A** | **MIC of antibiotic in the presence of Pt1 (µg/mL)**  **B** | **FIC A** | **FIC B** | **FIC**  **(FICA+FICB)** | **Interference** |
| --- | --- | --- | --- | --- | --- | --- | --- |
| **Pt1** | 0.125 | 0.125 |  | 1 | 1 | 2 | no interaction |
| **Ceftazidime** | 16 | 0.125 | 16 | 1 | 1 | 2 | no interaction |
| **Daptomycin** | 1 | 0.125 | 1 | 1 | 1 | 2 | no interaction |
| **Gentamicin** | 0.25 | 0.125 | 0.25 | 1 | 1 | 2 | no interaction |
| **Levofloxacin** | 0.25 | 0.125 | 0.25 | 1 | 1 | 2 | no interaction |
| **Linezolid** | 2 | 0.125 | 2 | 1 | 1 | 2 | no interaction |
| **Meropenem** | 0.25 | 0.125 | 0.25 | 1 | 1 | 2 | no interaction |
| **Minocycline** | 0.125 | 0.125 | 0.125 | 1 | 1 | 2 | no interaction |
| **Rifampicin** | 0.0078 | 0.125 | 0.0078 | 1 | 1 | 2 | no interaction |
| **Vancomycin** | 1 | 0.125 | 1 | 1 | 1 | 2 | no interaction |

**Table S8:** Antibiogram of *S. aureus* ATCC 29213 grown in the presence of sub-inhibitory concentration of PT-1/levofloxacin for a period of 36 days.

|  |  | **MIC (µg/mL)** | | |
| --- | --- | --- | --- | --- |
| **S.No.** | **Drug** | **36th Passage *S.aureus* ATCC 29213 in the presence of Pt-1** | **36th Passage *S.aureus* ATCC 29213 in the presence of Levofloxacin** | ***S.aureus* ATCC 29213** |
| 1 | **Pt-1** | 0.125 | - | 0.0625 |
| 2 | Ceftazidime | 16 | 8 | 16 |
| 3 | Daptomycin | 1 | 1 | 1 |
| 4 | Gentamycin | 0.5 | 0.25 | 0.25 |
| 5 | Levofloxacin | 0.25 | 32 | 0.25 |
| 6 | Linezolid | 2 | 2 | 2 |
| 7 | Meropenem | 0.25 | 0.125 | 0.25 |
| 8 | Minocycline | 0.125 | 0.125 | 0.0625 |
| 9 | Rifampicin | 0.0156 | 0.0156 | 0.0156 |
| 10 | Vancomycin | 1 | 1 | 1 |

**Table S9**: *In vitro* Post Antibiotic effect (PAE) of Pt-1 at different concentrations.

| **Treatments** | **Time for 1 log_10_ increase in bacterial cfu count (h)** | **PAE (h)** |
| --- | --- | --- |
| **Untreated *S. aureus* ATCC 29213** | **~2.5** | **0** |
| **Pt-1 1x** | **~8** | **~5.5** |
| **Pt-1 5x** | **~8** | **~5.5** |
| **Pt-1 10x** | **~8** | **~5.5** |
| **Vancomycin 1x** | **~5** | **~2.5** |
| **Vancomycin 5x** | **~5** | **~2.5** |
| **Vancomycin 10x** | **~5** | **~2.5** |
| **Levofloxacin 1x** | **~3** | **~0.5** |
| **Levofloxacin 5x** | **~4** | **~1.5** |
| **Levofloxacin 10x** | **~4** | **~1.5** |

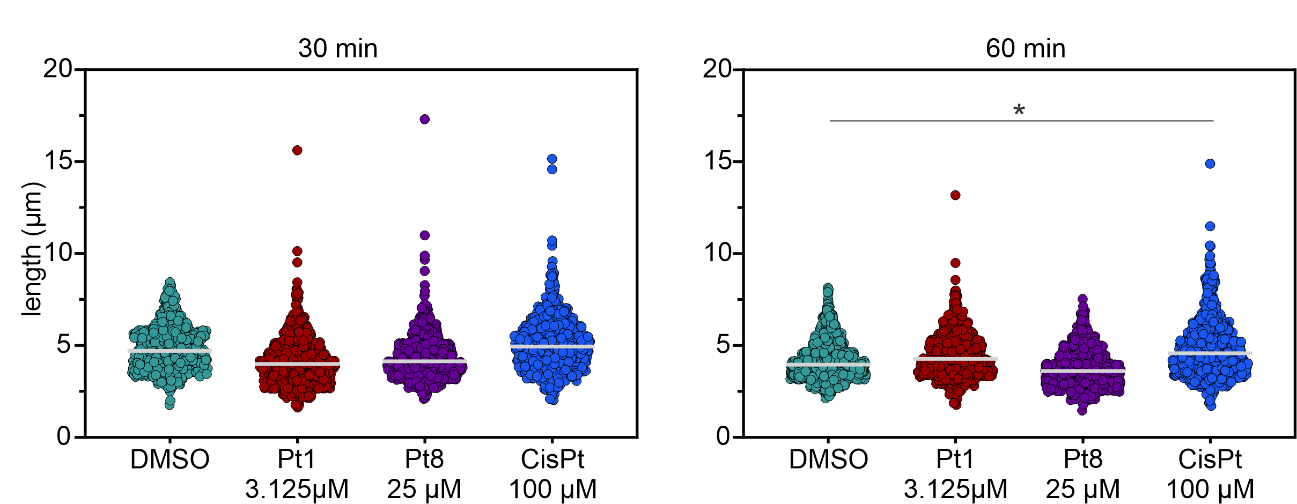

**Figure S15**: Cell length measured from BCP images. Grey lines indicate the median. Significance was tested with nested t-tests (p value: * ≤ 0.5).

**Table S10**: *Bacillus subtilis* strains used in mode of action assays. *mgfp*= monomeric green-fluorescent protein.

| **Strain** | **Induction** | **Genotype** | **Reference** |
| --- | --- | --- | --- |
| 168CA | **-** | *trpC2* | ^7 8^ |
| bSS82 | **-** | *amyE::spc PrpsD-gfp* | ^9^ |
| UG10 | 0.5 % xylose | *amyE::spc Pxyl-recA-mgfp* | ^8^**^,^**^10^ |
| 1048 | 1 % xylose | *cat rpoC-gfp Pxyl-‘rpoC* | ^8^**^,^**^11^ |
| 1049 | 1 % xylose | *amyE::spc Pxyl-rpsB-gfp* | ^8^**^,^**^11^ |
| LH131 | 0.1 % xylose* | *amyE::spc Pxyl-gfp-minD* | ^8^**^,^**^12^ |
| MW10 | 0.1 % xylose | *amyE::spc Pxyl-gfp-mreB* | ^9^**^,^**^13^ |
| 2020 | 0.3 % xylose | *amyE::spc Pxyl-gfp-ftsZ* | ^14^ |
| BKE08820 | **-** | *trpC2ΔkatA::erm* | ^15^ |
| BKE25020 | **-** | *trpC2ΔsodA::erm* | ^5^ |

*Added to only day culture.

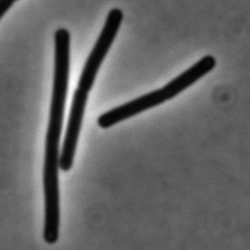
**NMR Spectra**

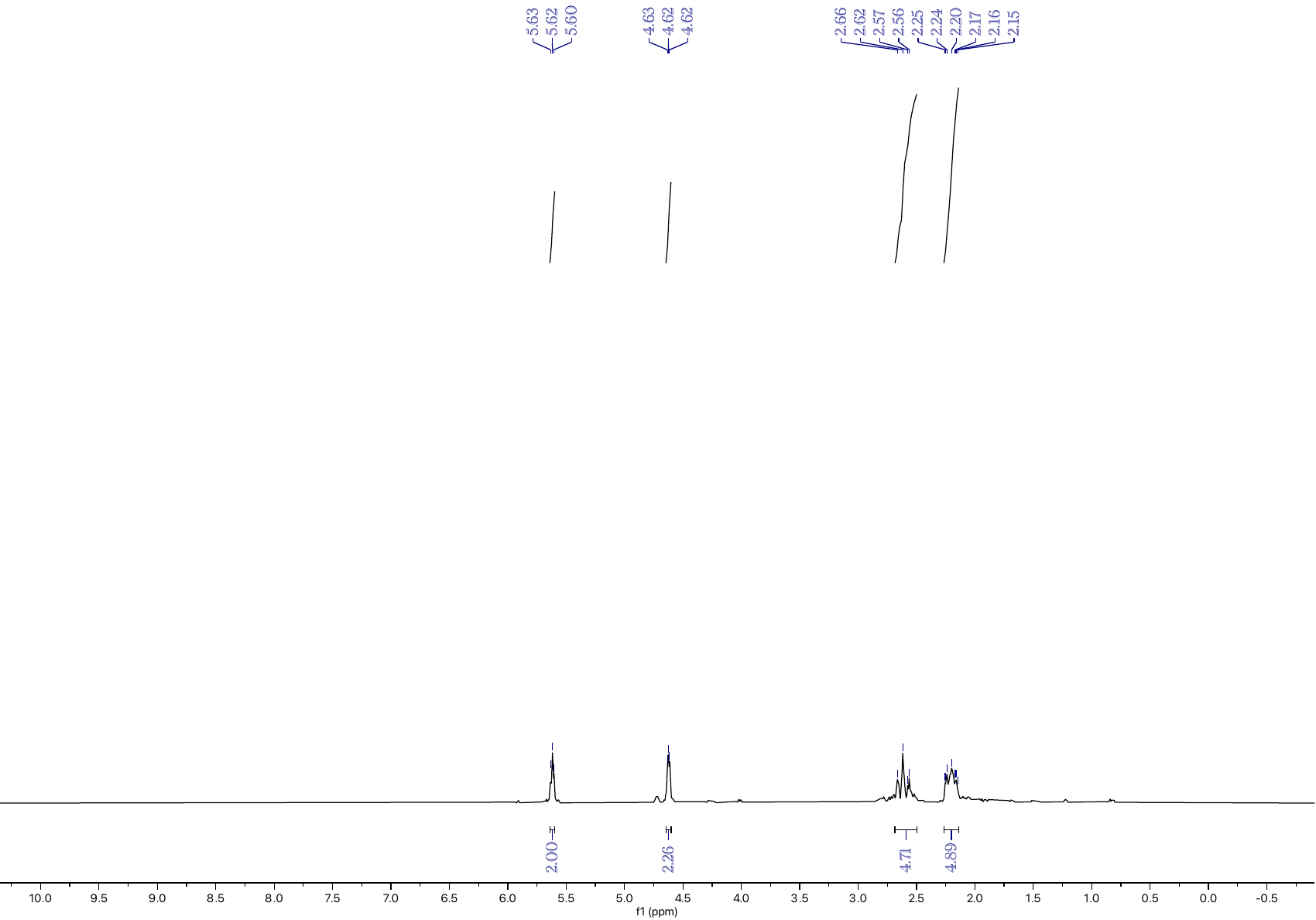

**Figure S16:**. ^1^H NMR spectrum of **1** in CDCl_3_.

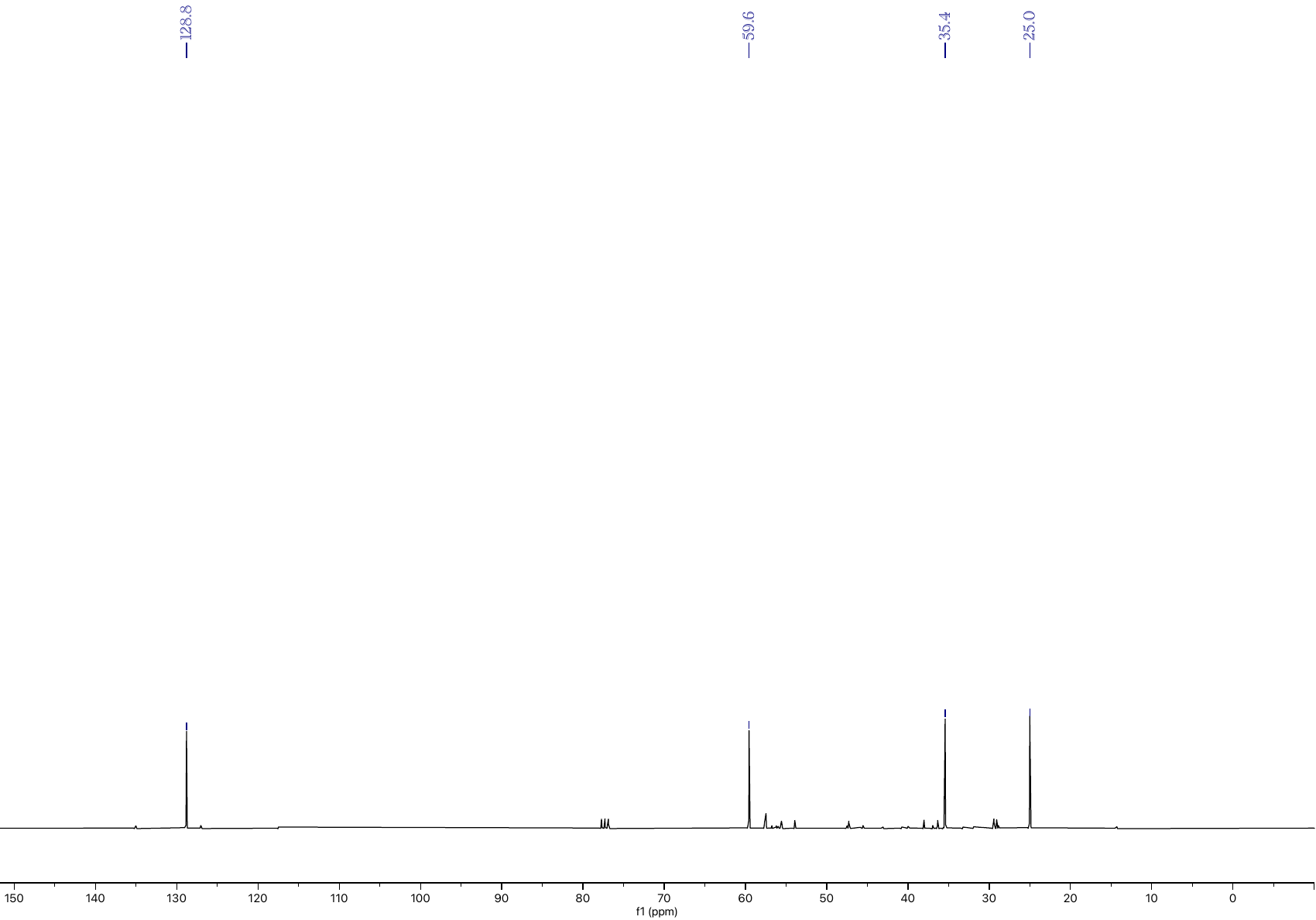

**Figure S17:** ^13^C NMR spectrum of **1** in CDCl_3_.

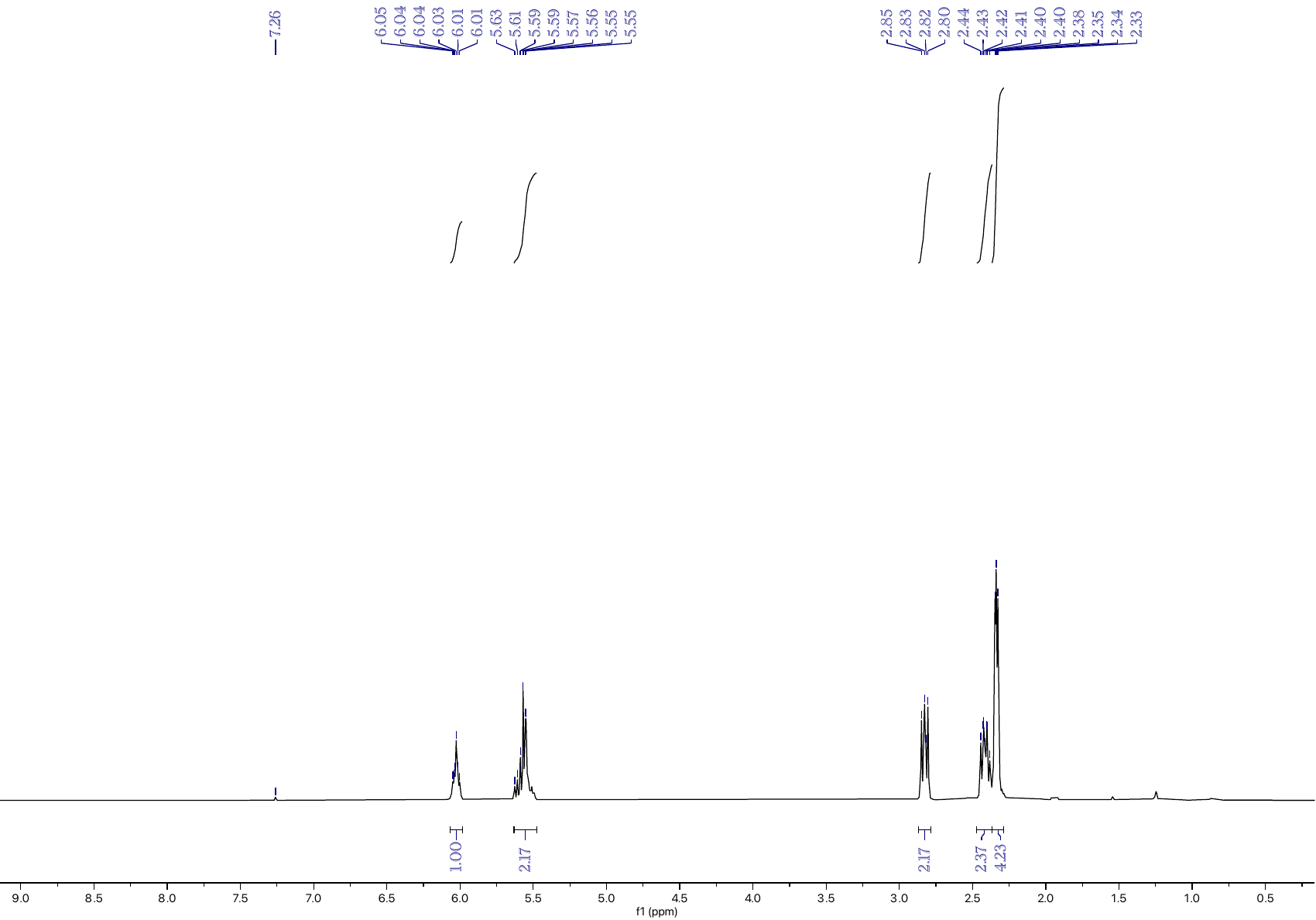

**Figure S18**: ^1^H NMR spectrum of **2** in CDCl_3_.

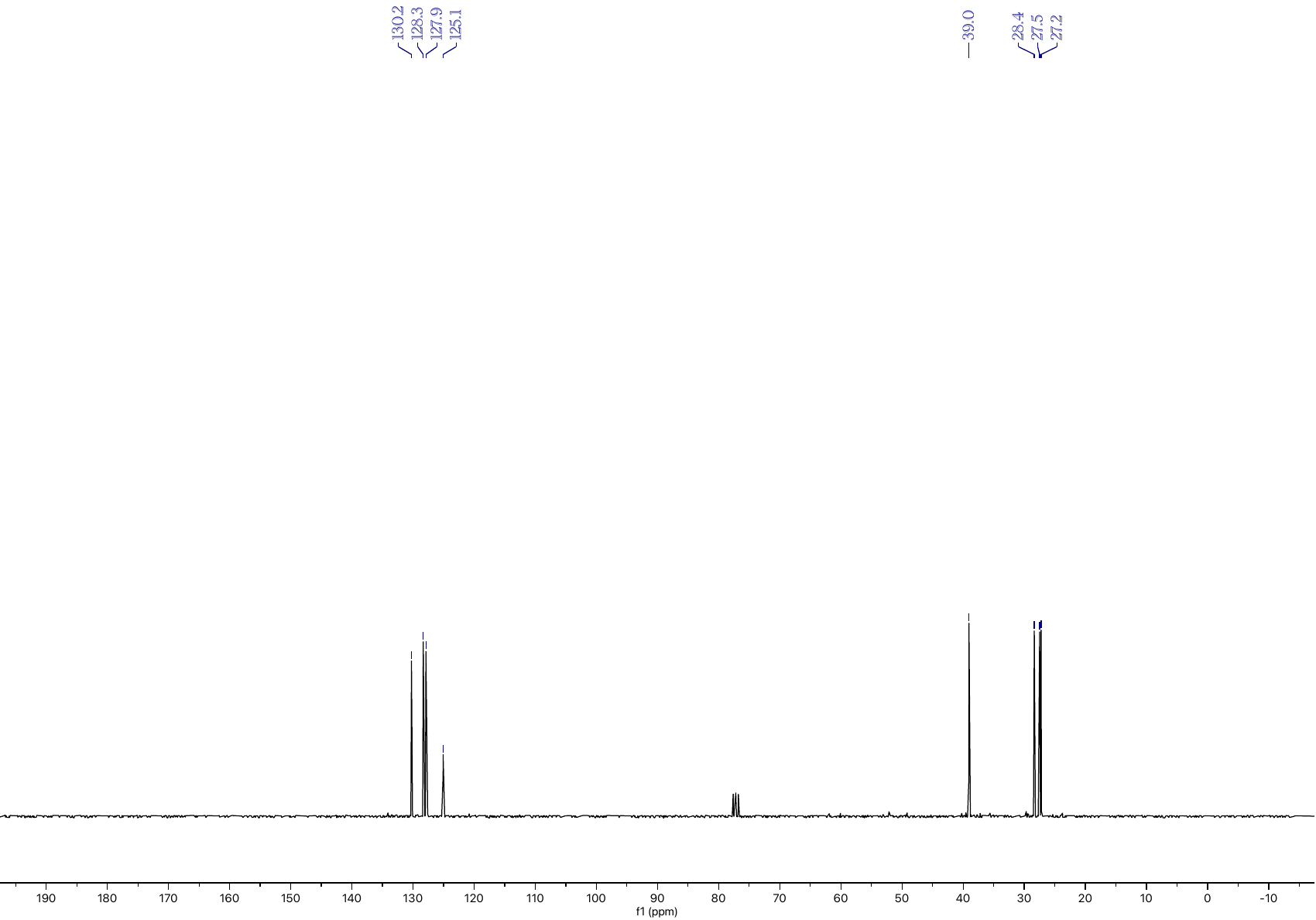

**Figure S19:** ^13^C NMR spectrum of **2** in CDCl_3_.

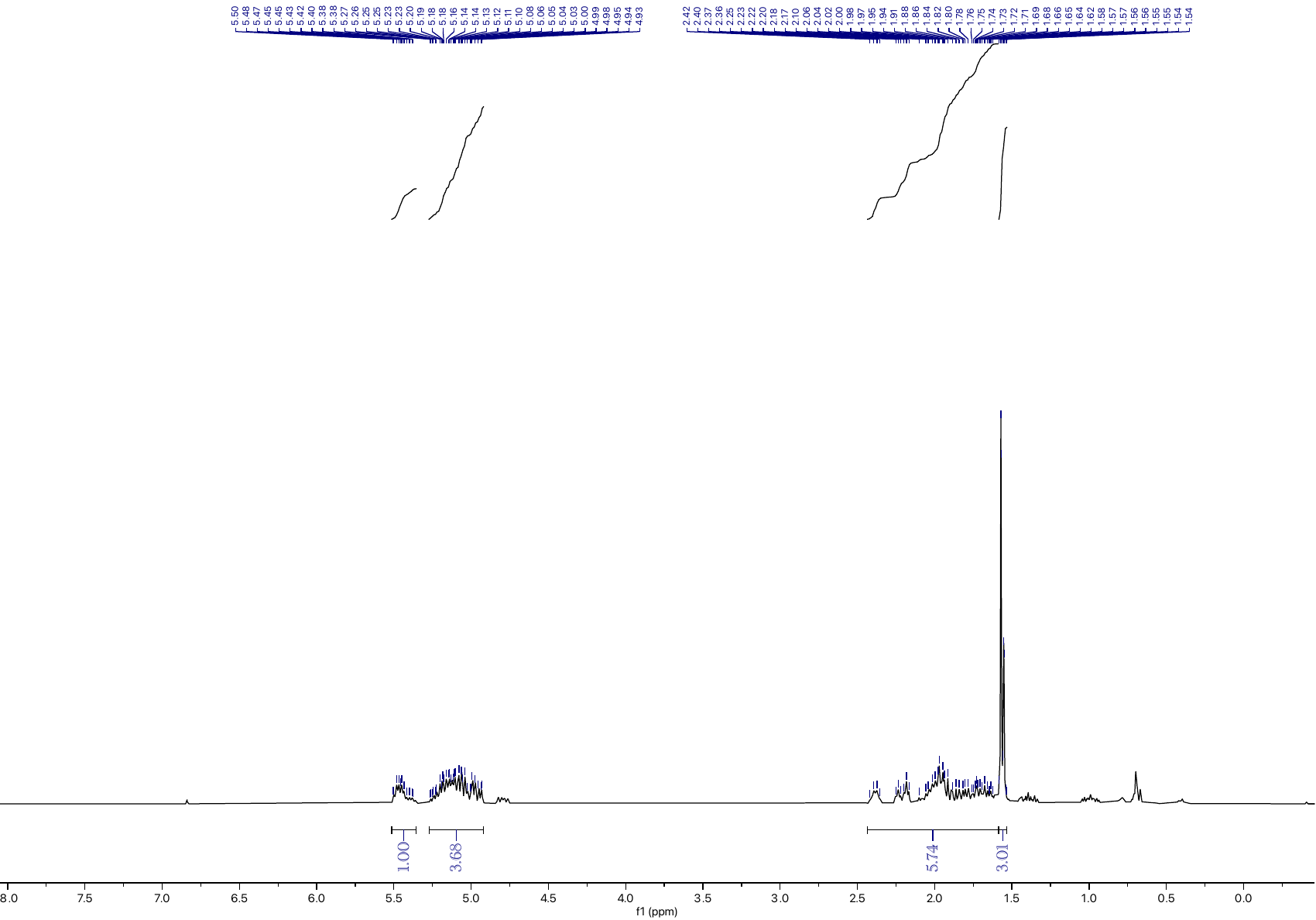

**Figure S20**: ^1^H NMR spectrum of **3** in CDCl_3_.

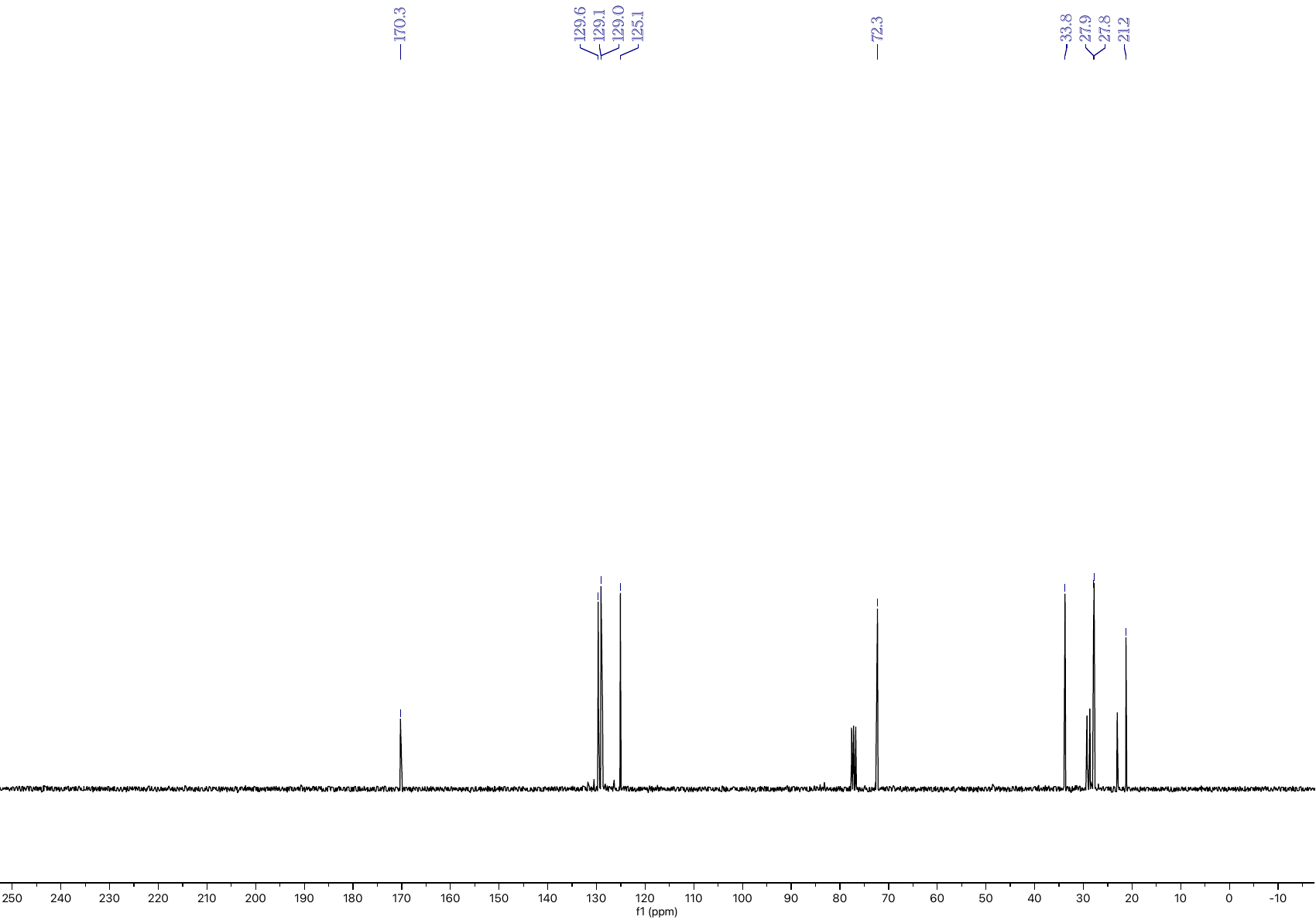

**Figure S21:** ^13^C NMR spectrum of **3** in CDCl_3_.

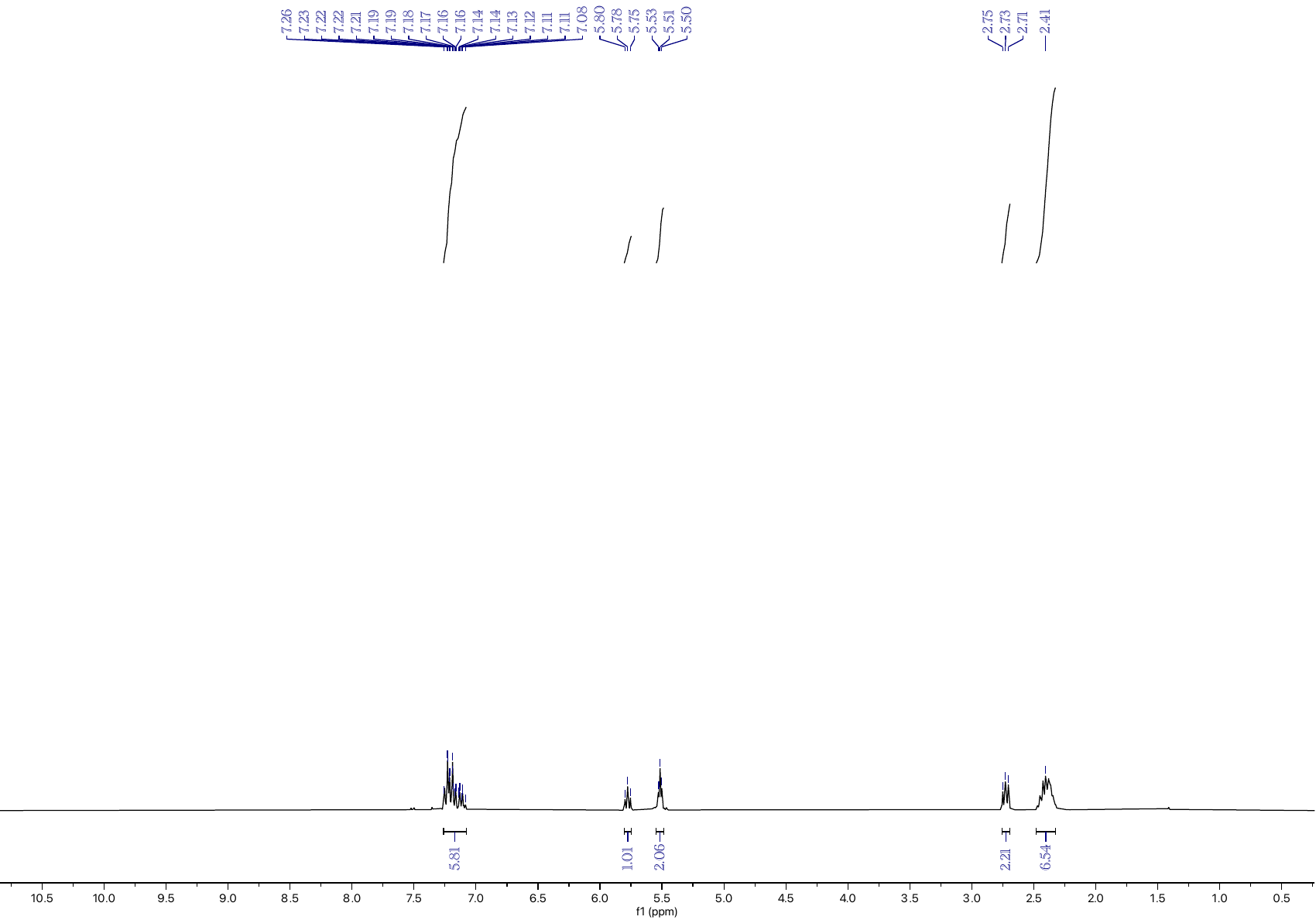

**Figure S22:** ^1^H NMR spectrum of **4** in CDCl_3_.

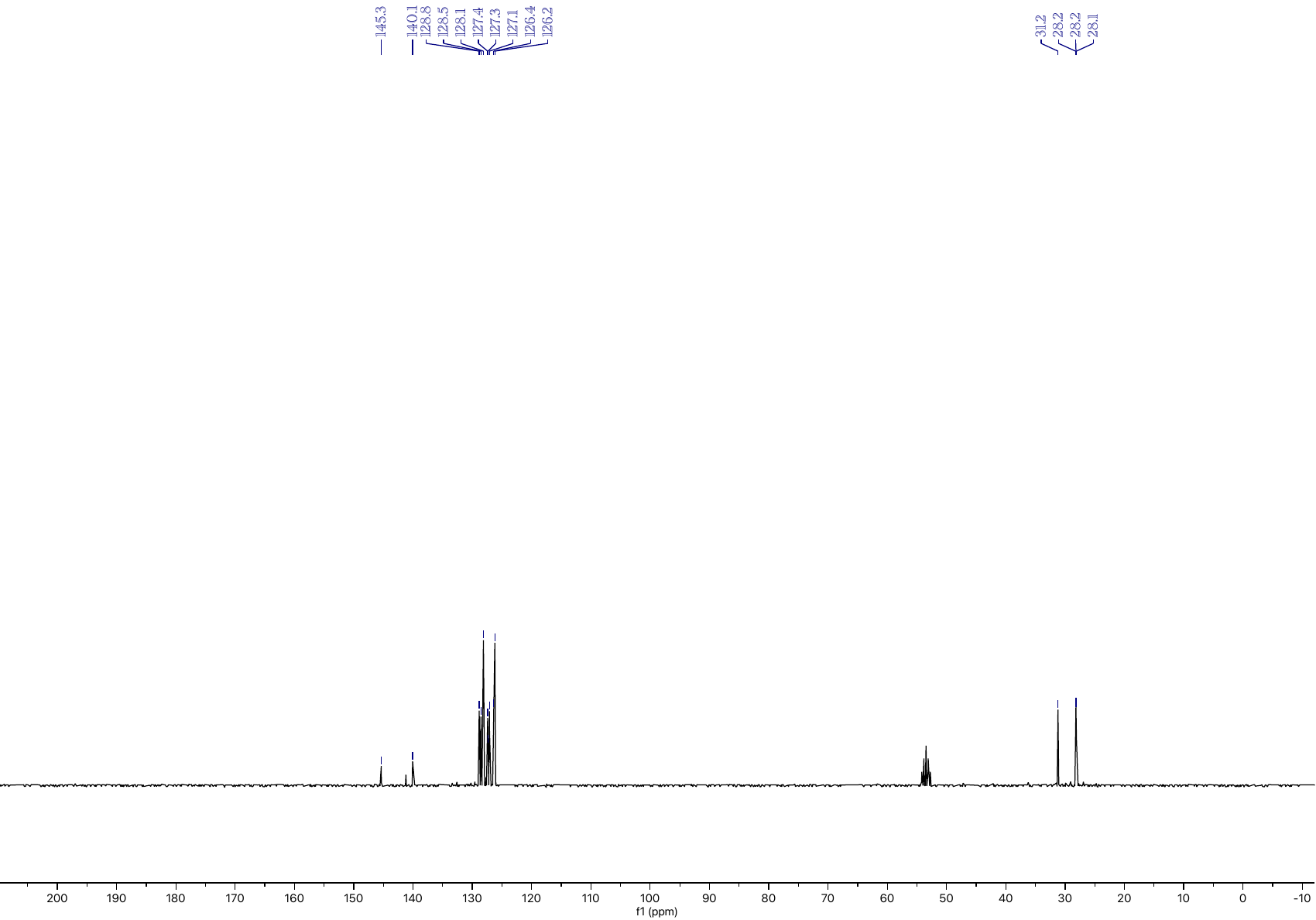

**Figure S23:** ^13^C NMR spectrum of **4** in CD_2_Cl_3_.

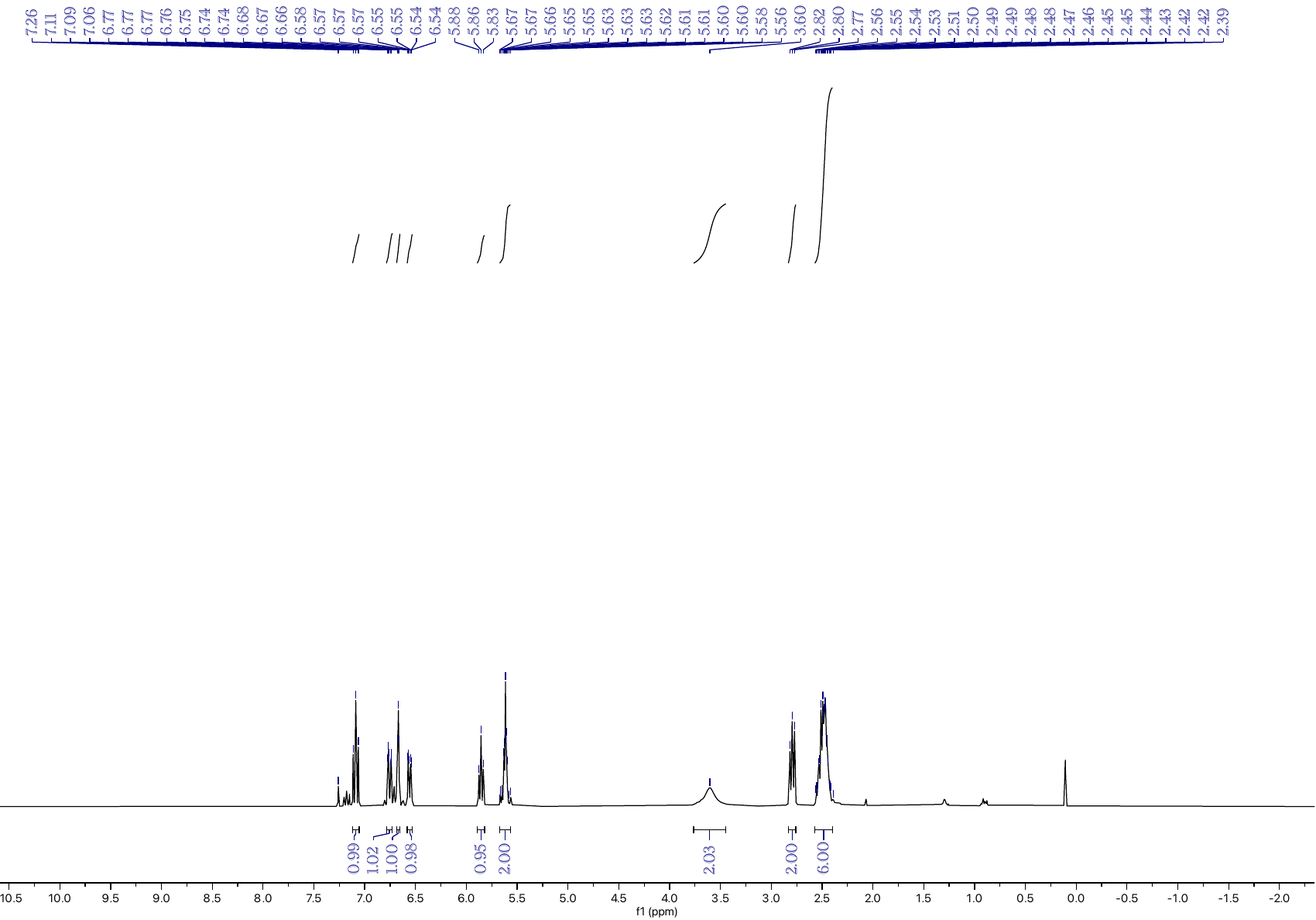

**Figure S24:** ^1^H NMR spectrum of **5** in CDCl_3_.

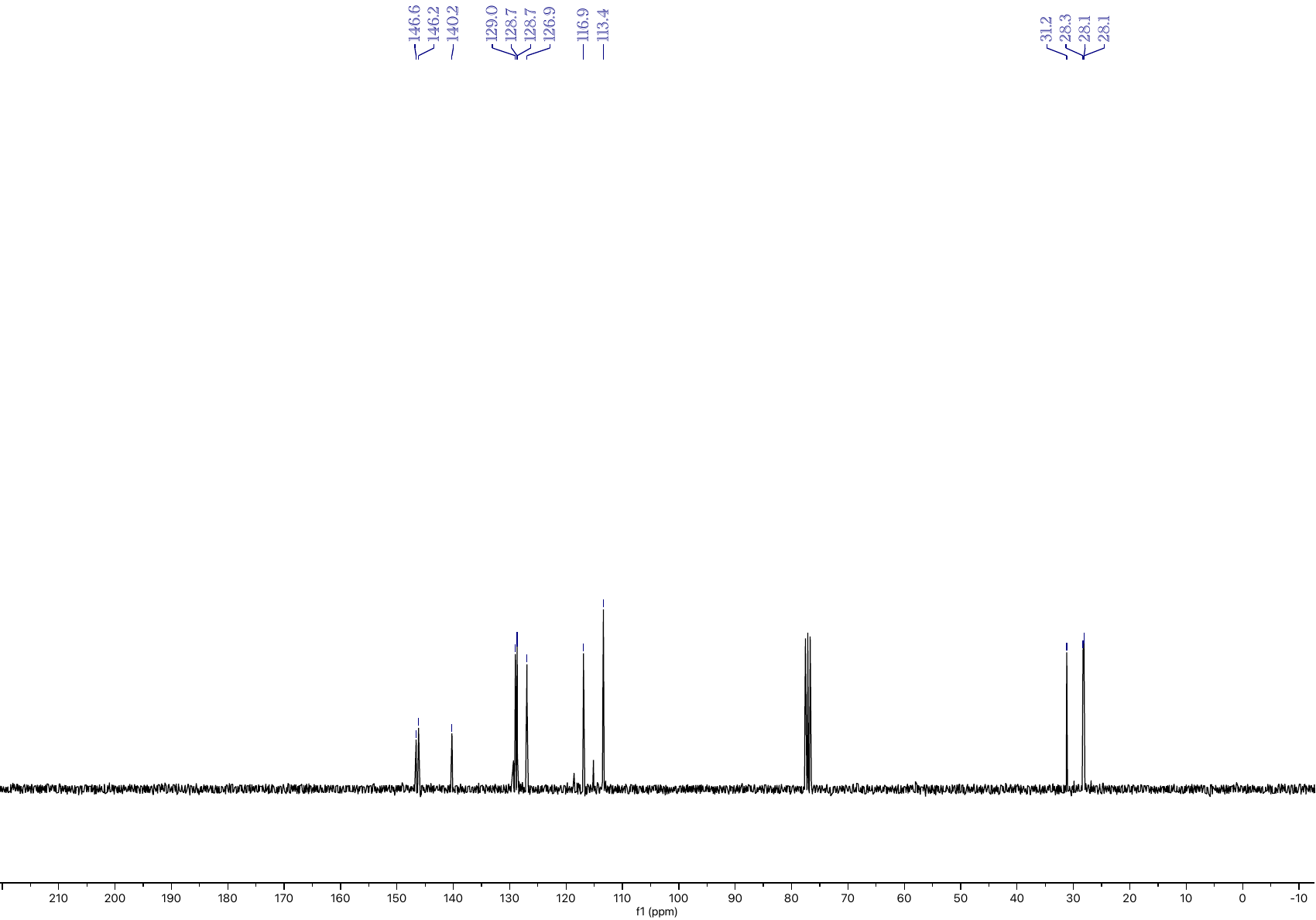

**Figure S25:** ^13^C NMR spectrum of **5** in CDCl_3_.

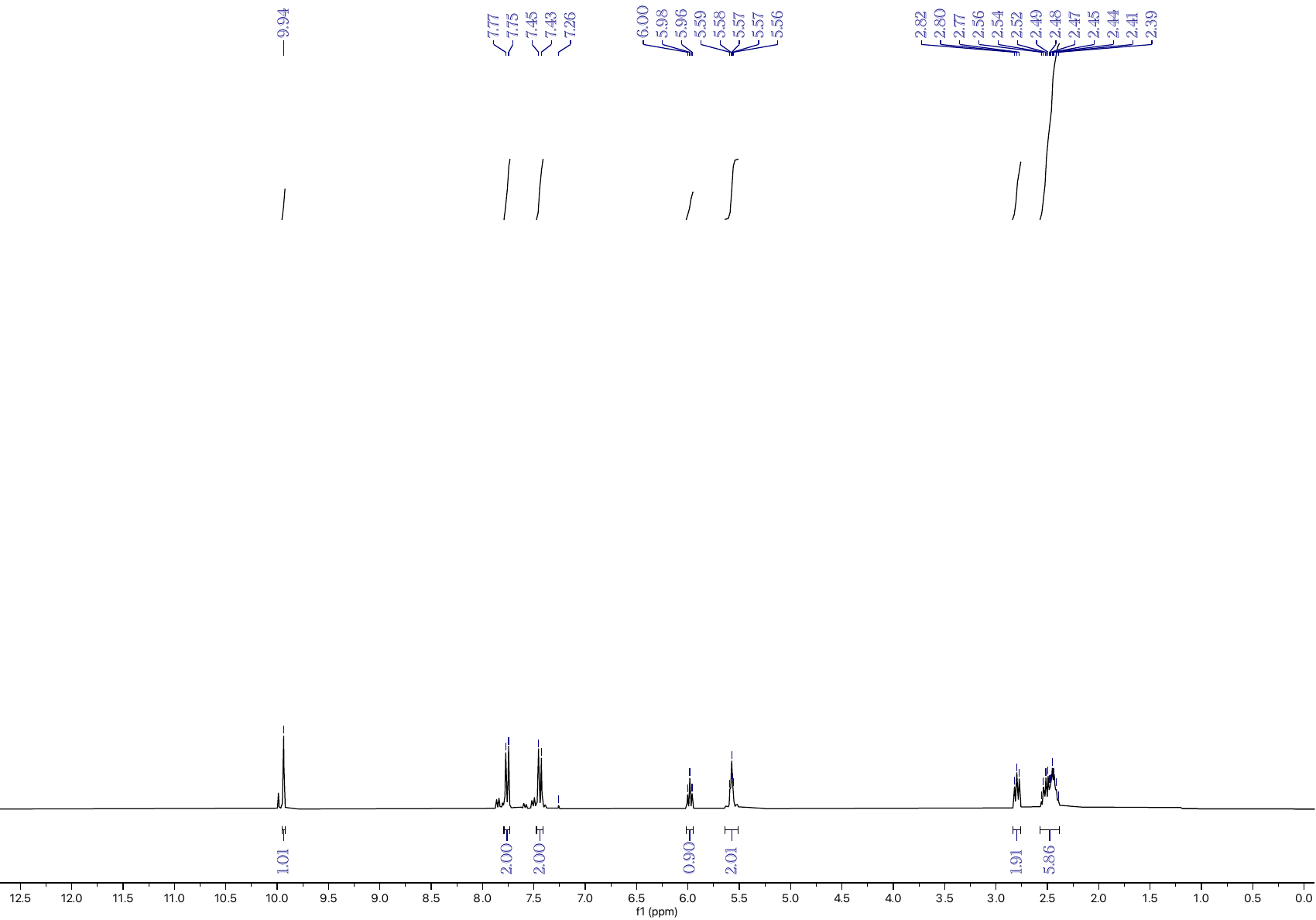

**Figure S26:** ^1^H NMR spectrum of **6** in CDCl_3_.

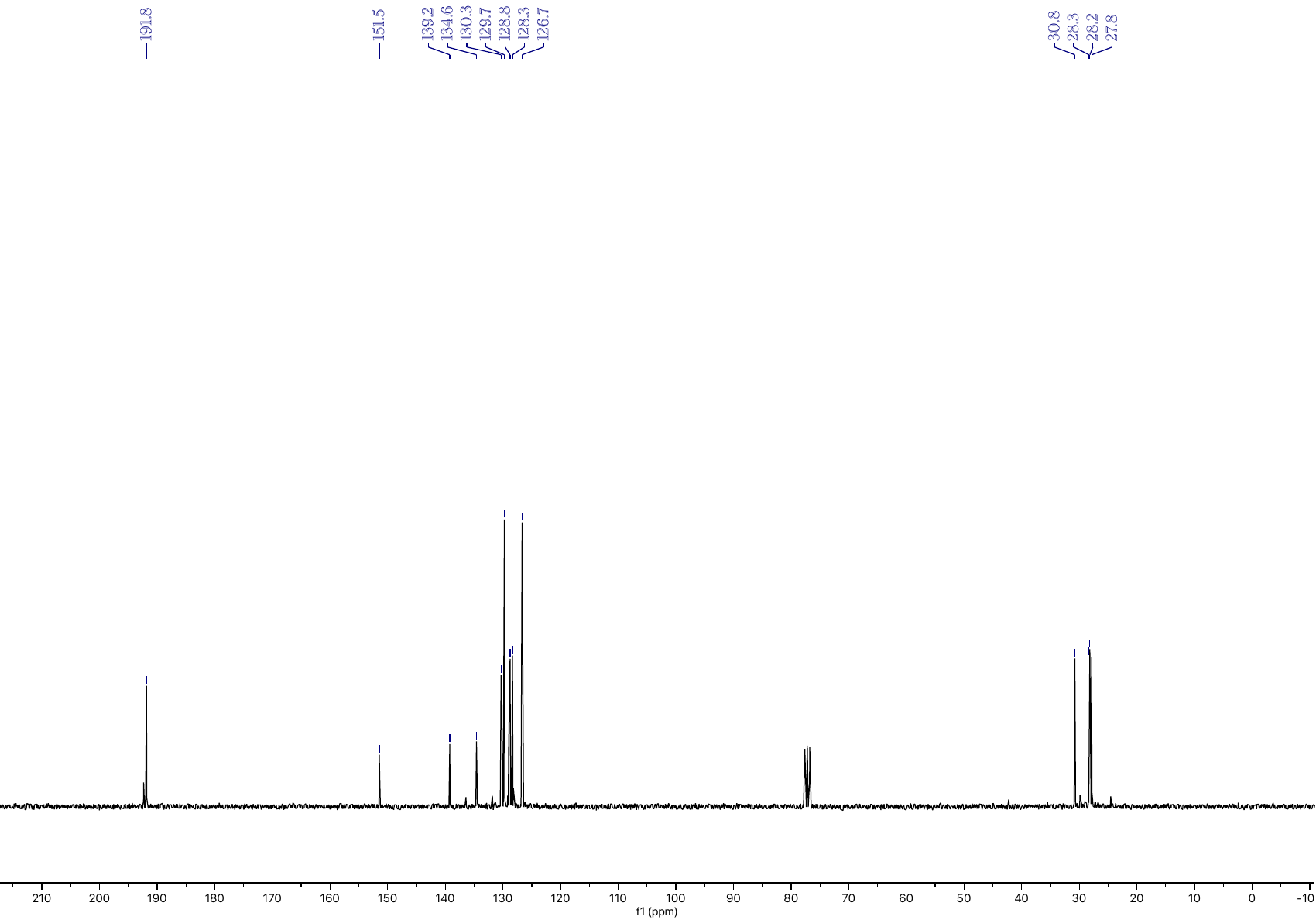

**Figure S27:** ^13^C NMR spectrum of **6** in CDCl_3_.

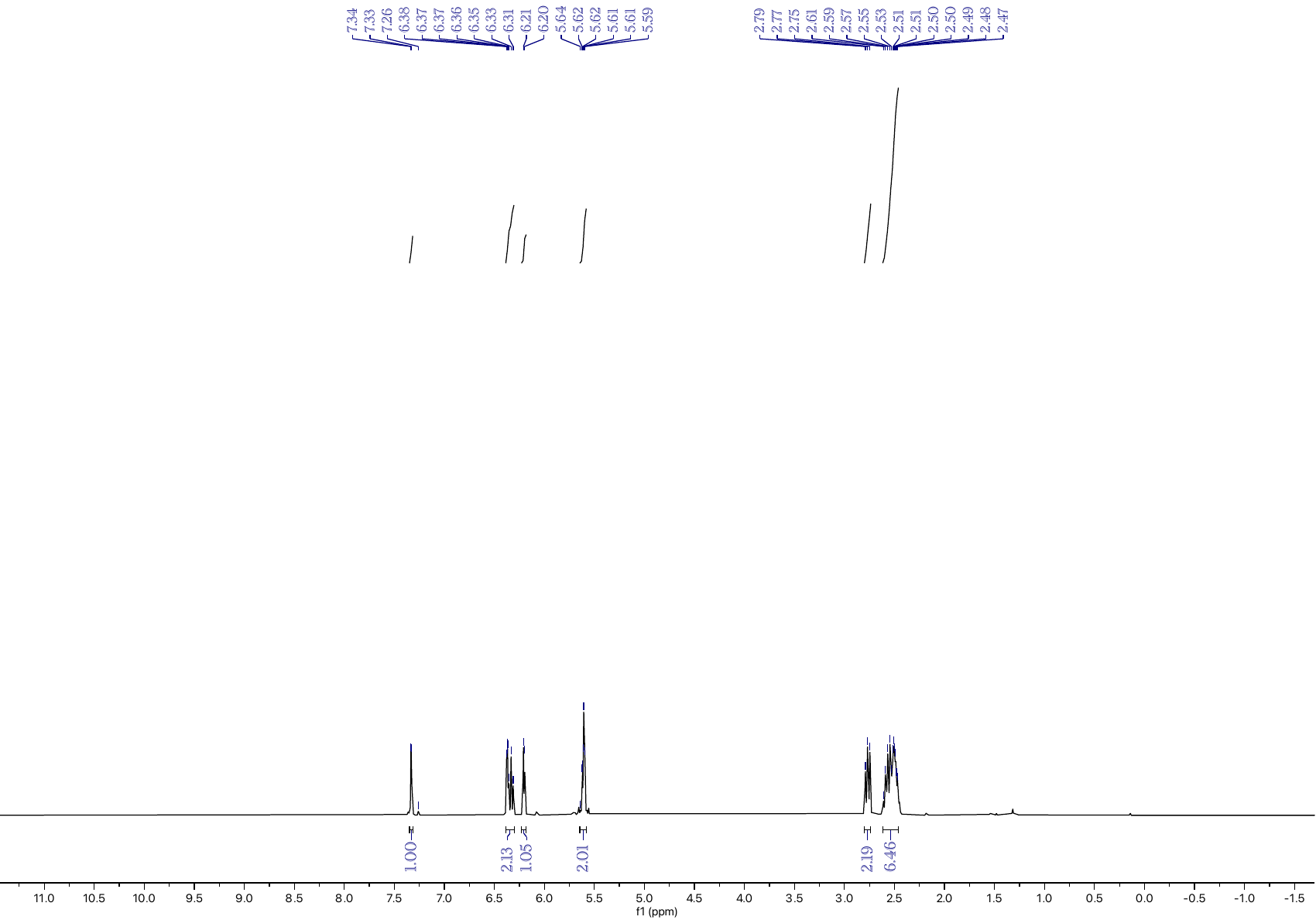

**Figure S28:** ^1^H NMR spectrum of **7** in CDCl_3_.

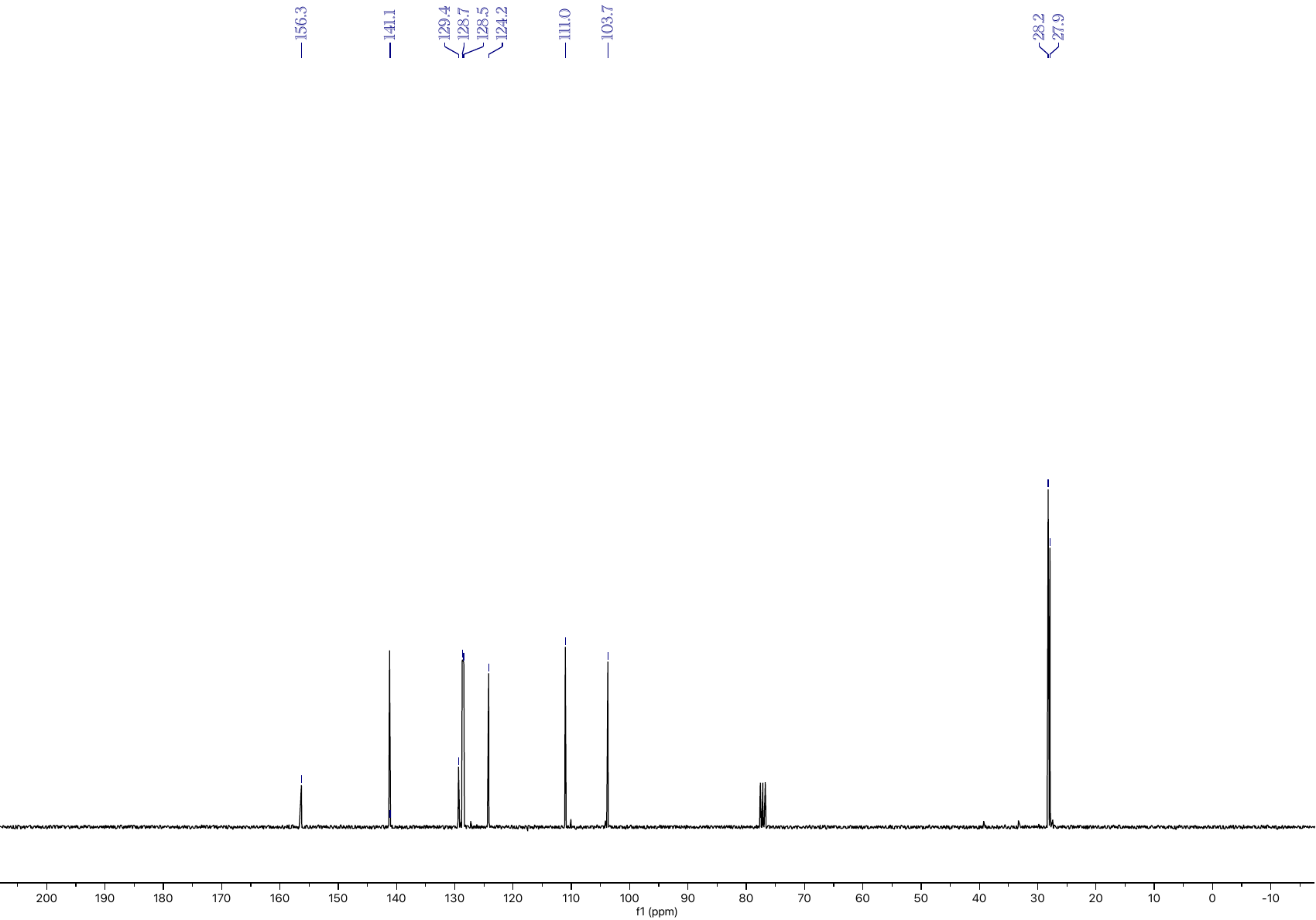

**Figure S29:** ^13^C NMR spectrum of **7** in CDCl_3_.

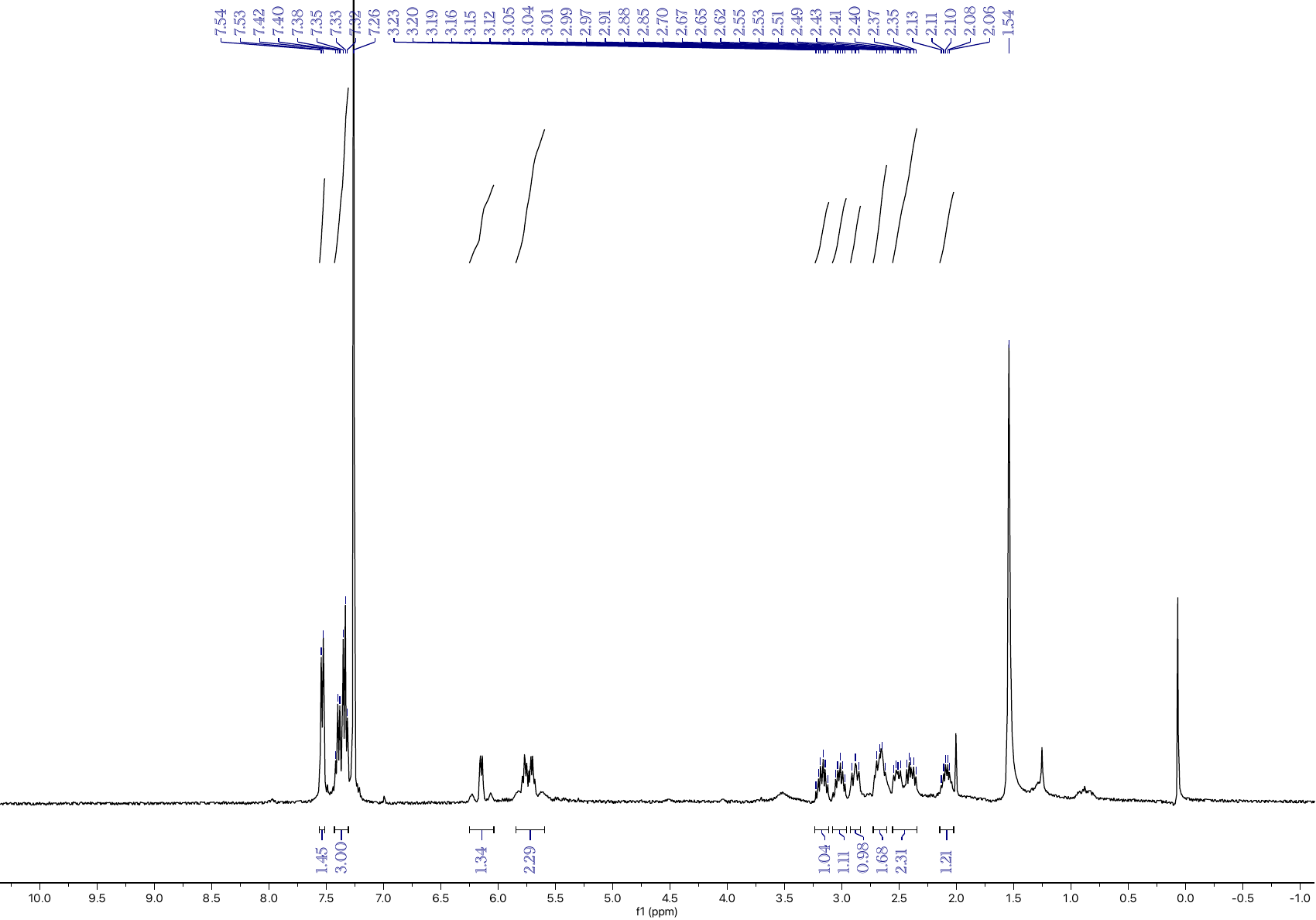

**Figure S30:** ^1^H NMR spectrum of **8** in CDCl_3_.

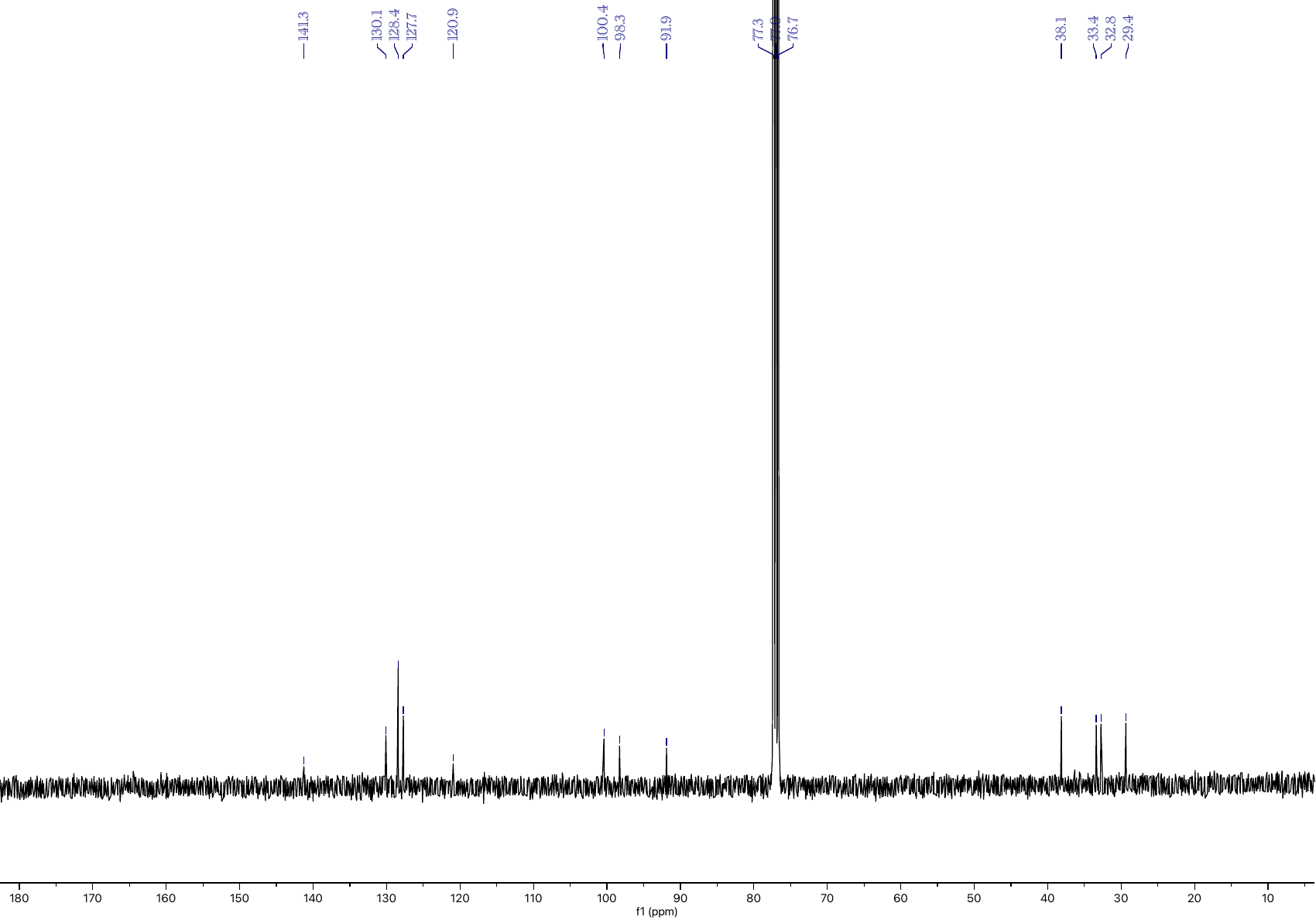

**Figure S31:** ^13^C NMR spectrum of **8** in CDCl_3_.

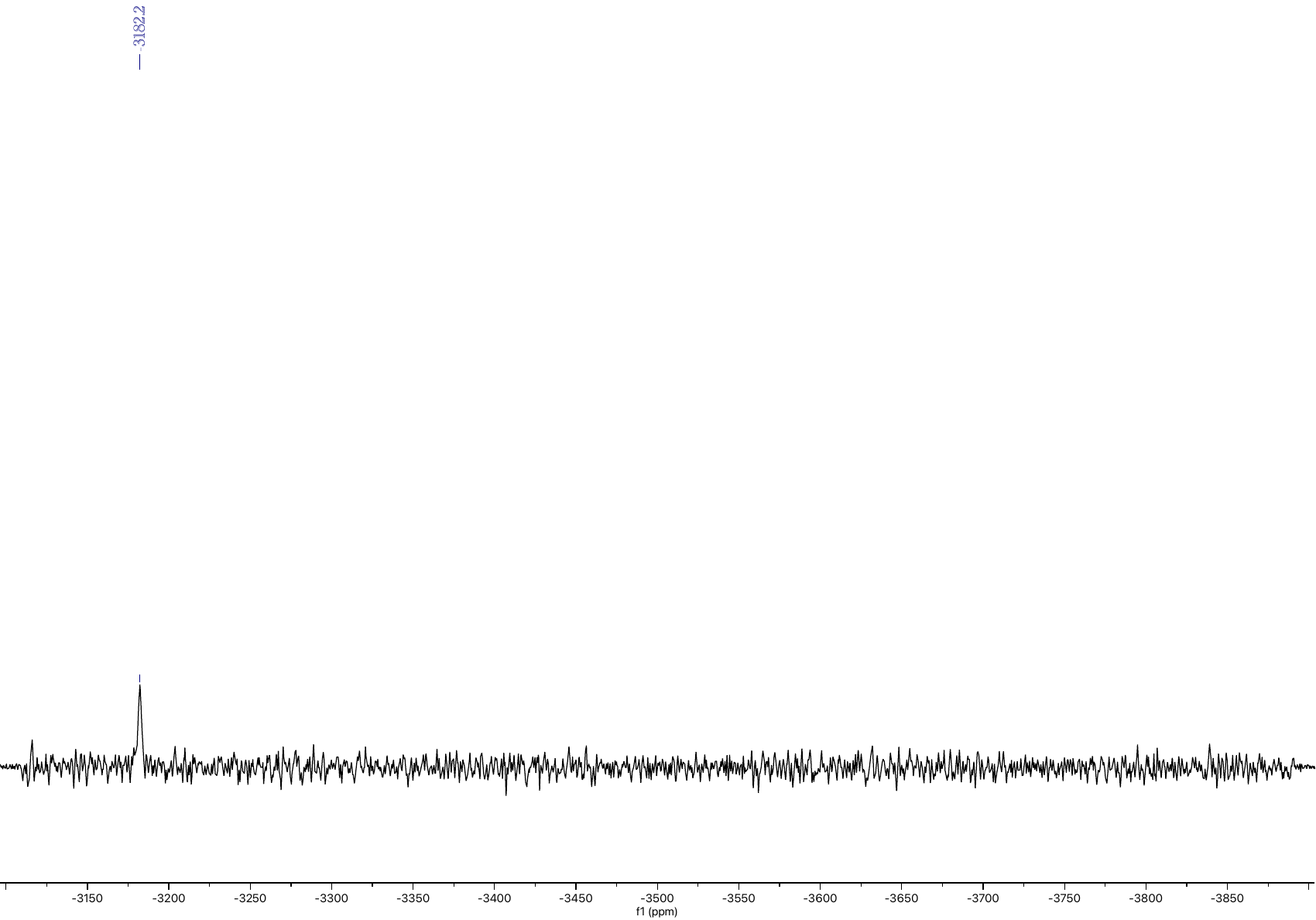

**Figure S32:** ^195^Pt NMR spectrum of **8** in DMSO.

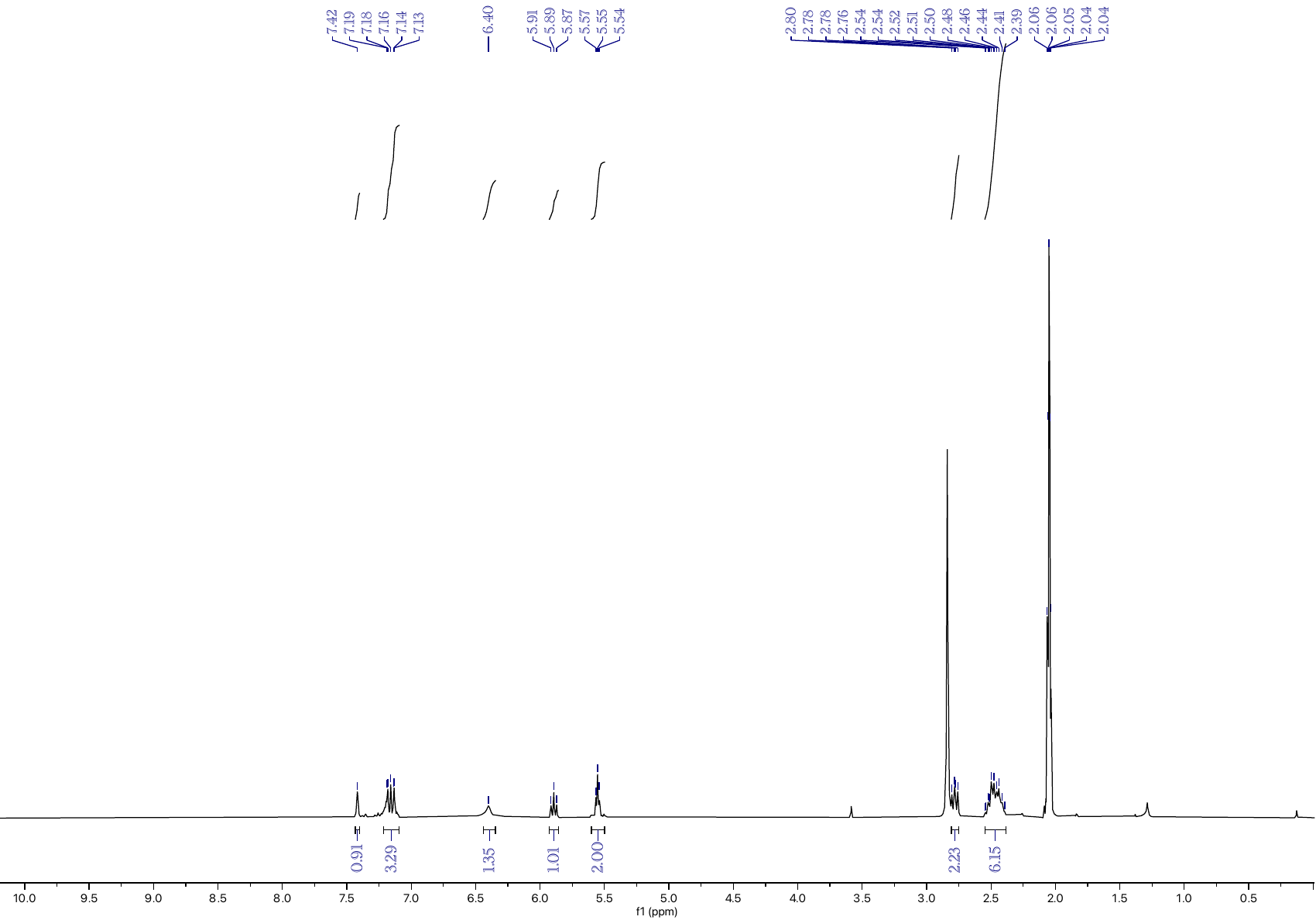

**Figure S33:** ^1^H NMR spectrum of **9** in acetone.

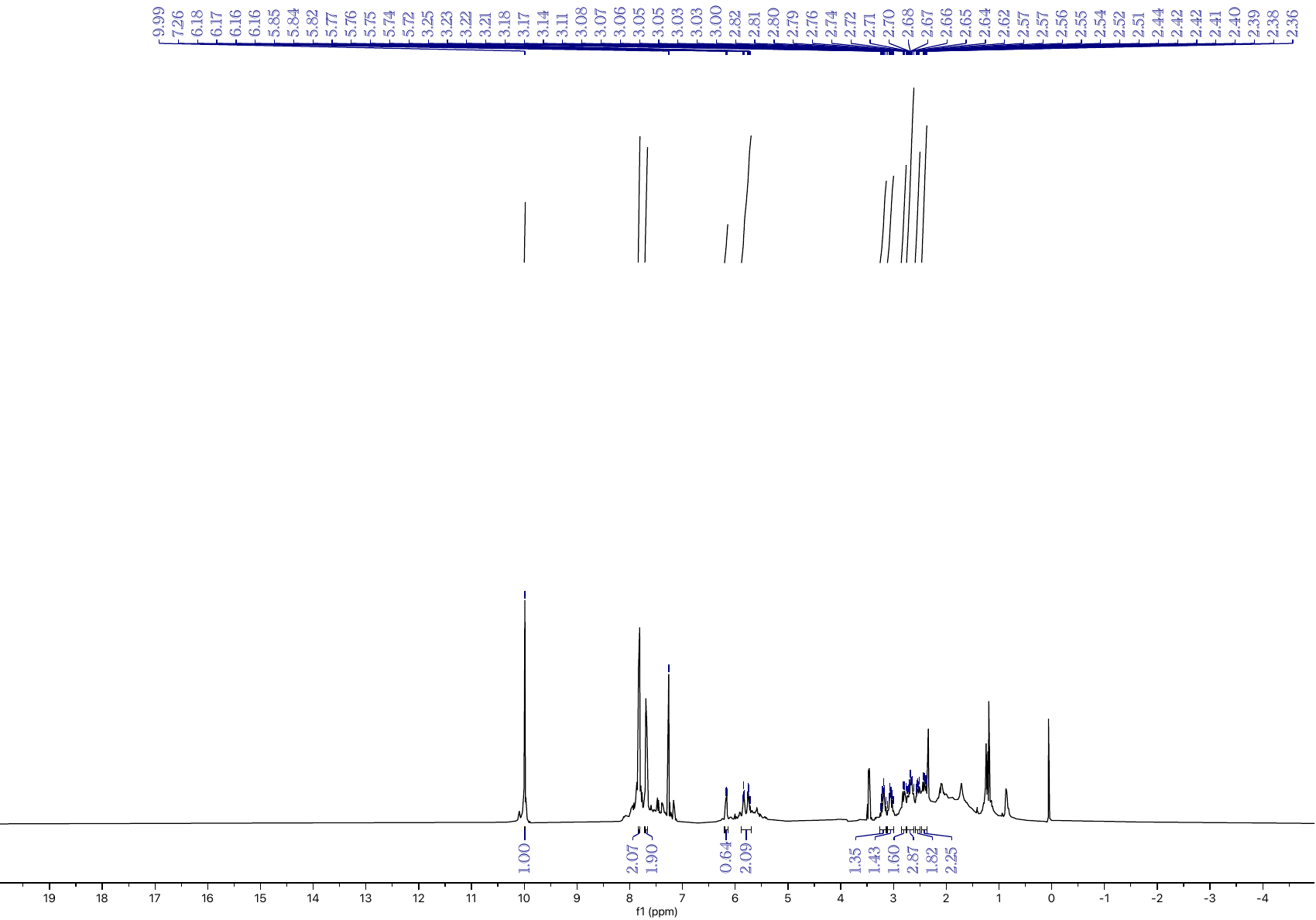

**Figure S34:** ^1^H NMR spectrum of **10** in CDCl_3_.

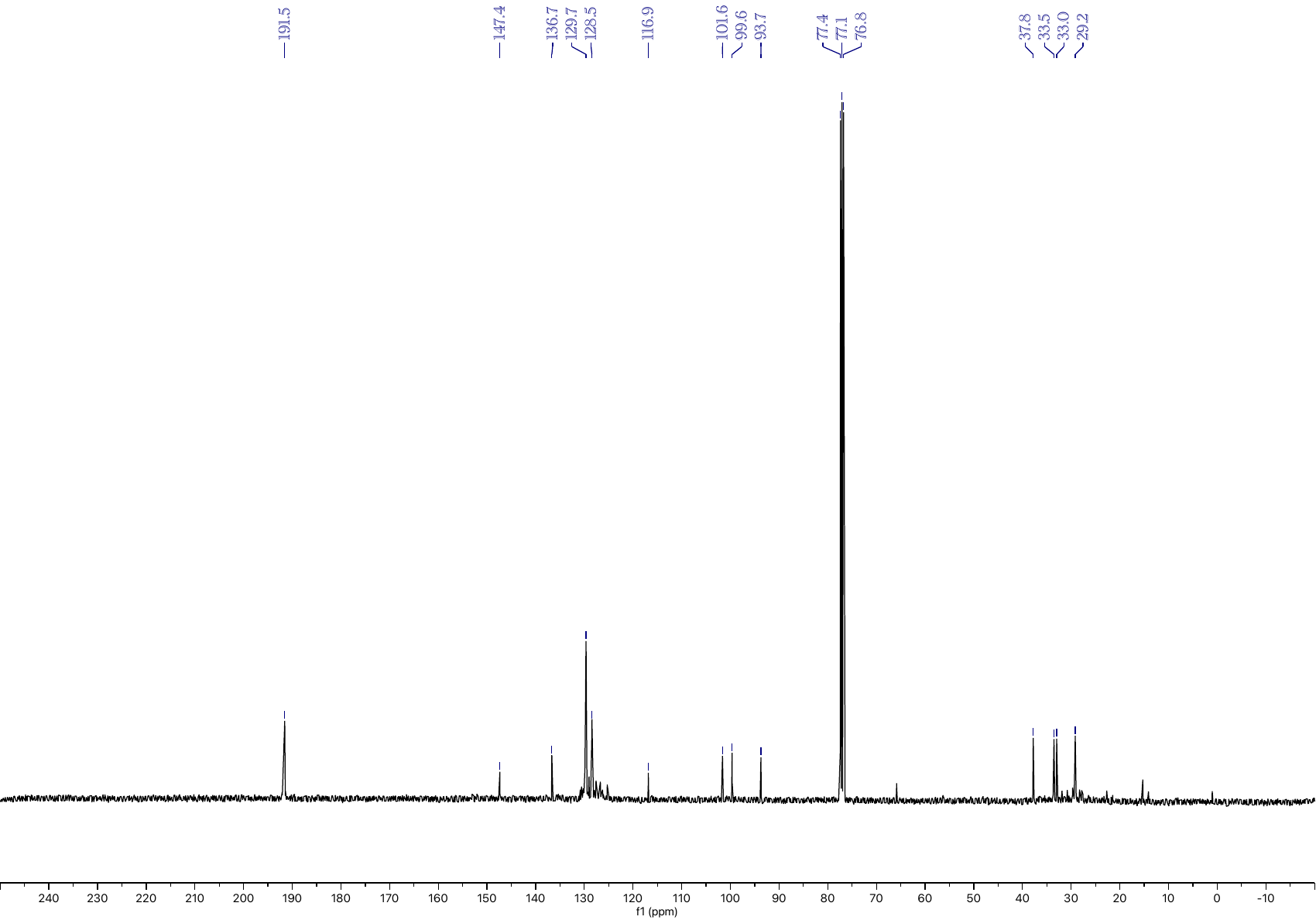

**Figure S35:** ^13^C NMR spectrum of **10** in CDCl_3_.

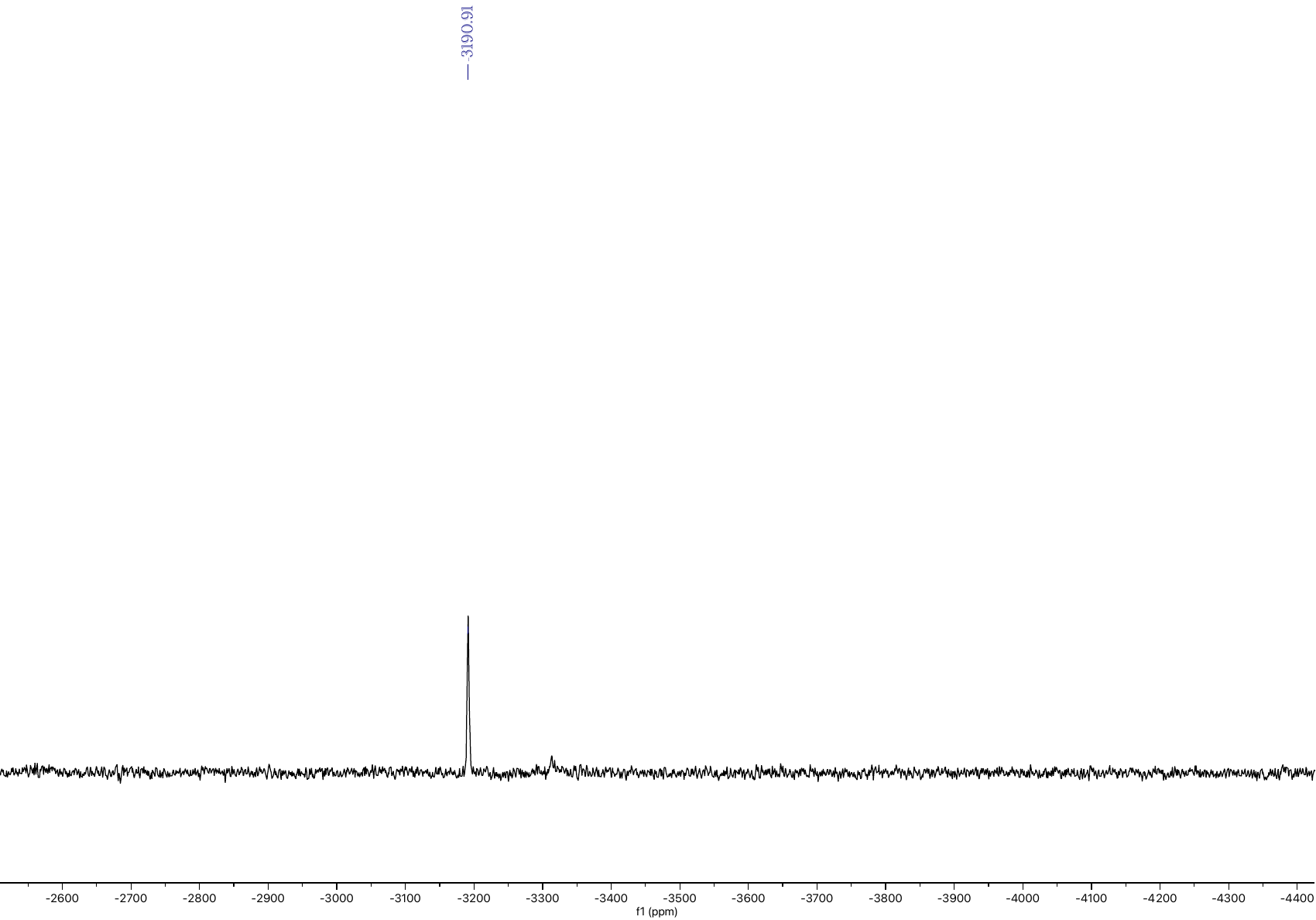

**Figure S36:** ^195^Pt NMR spectrum of **10** in CDCl_3_.

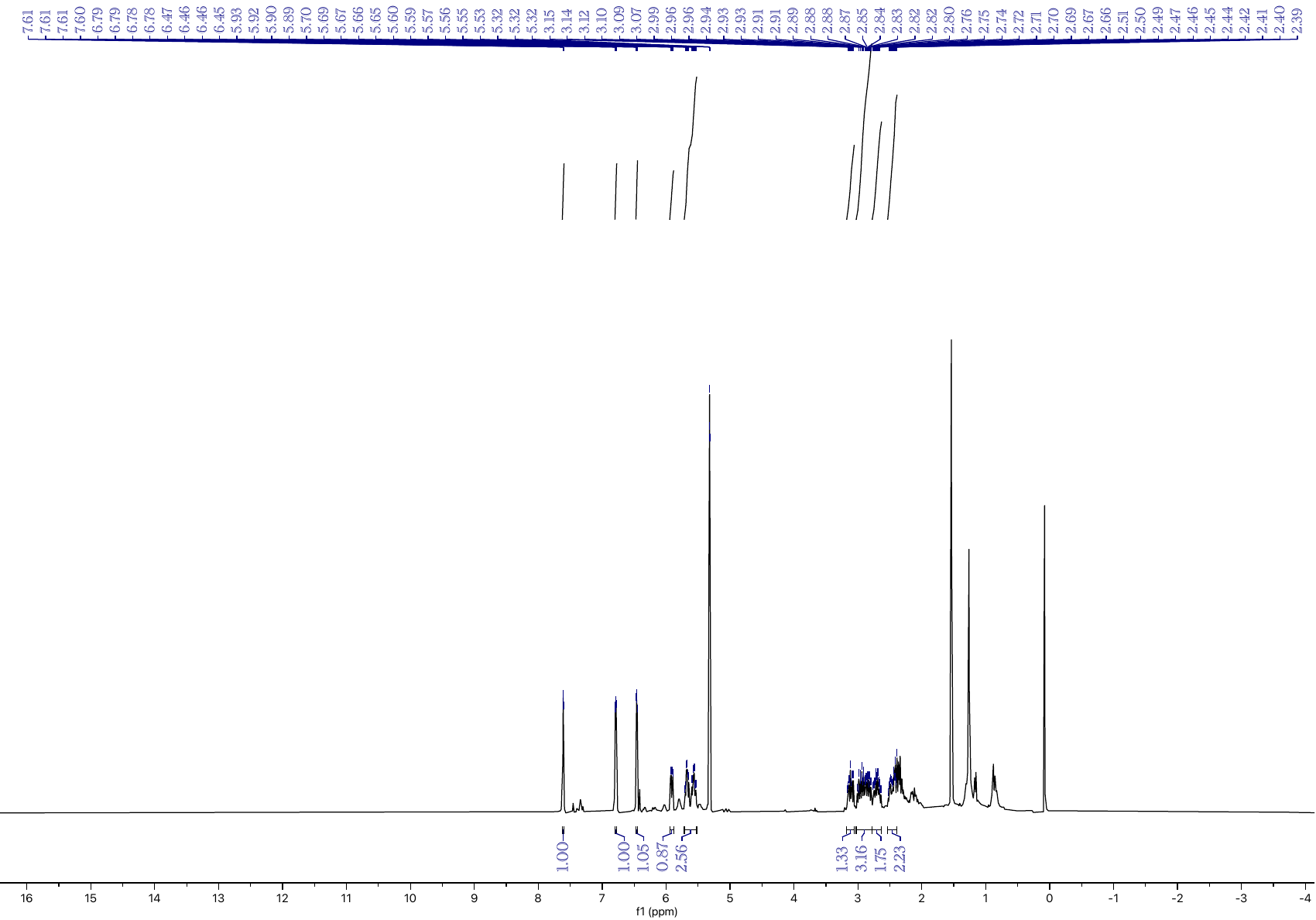

**Figure S37:** ^1^H NMR spectrum of **11** in CD_2_Cl_3_.

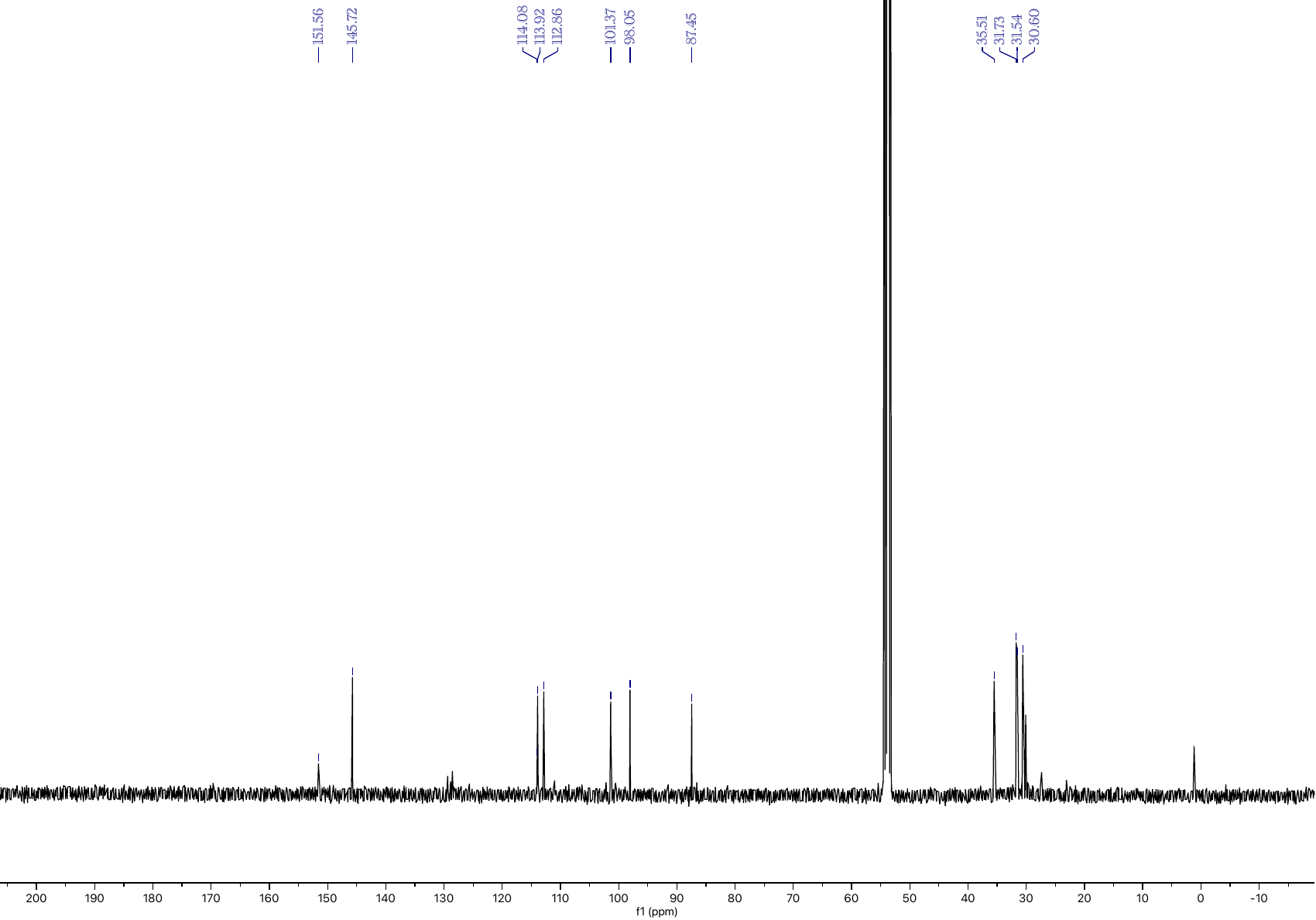

**Figure S38:** ^13^C NMR spectrum of **11** in CDCl_3_.

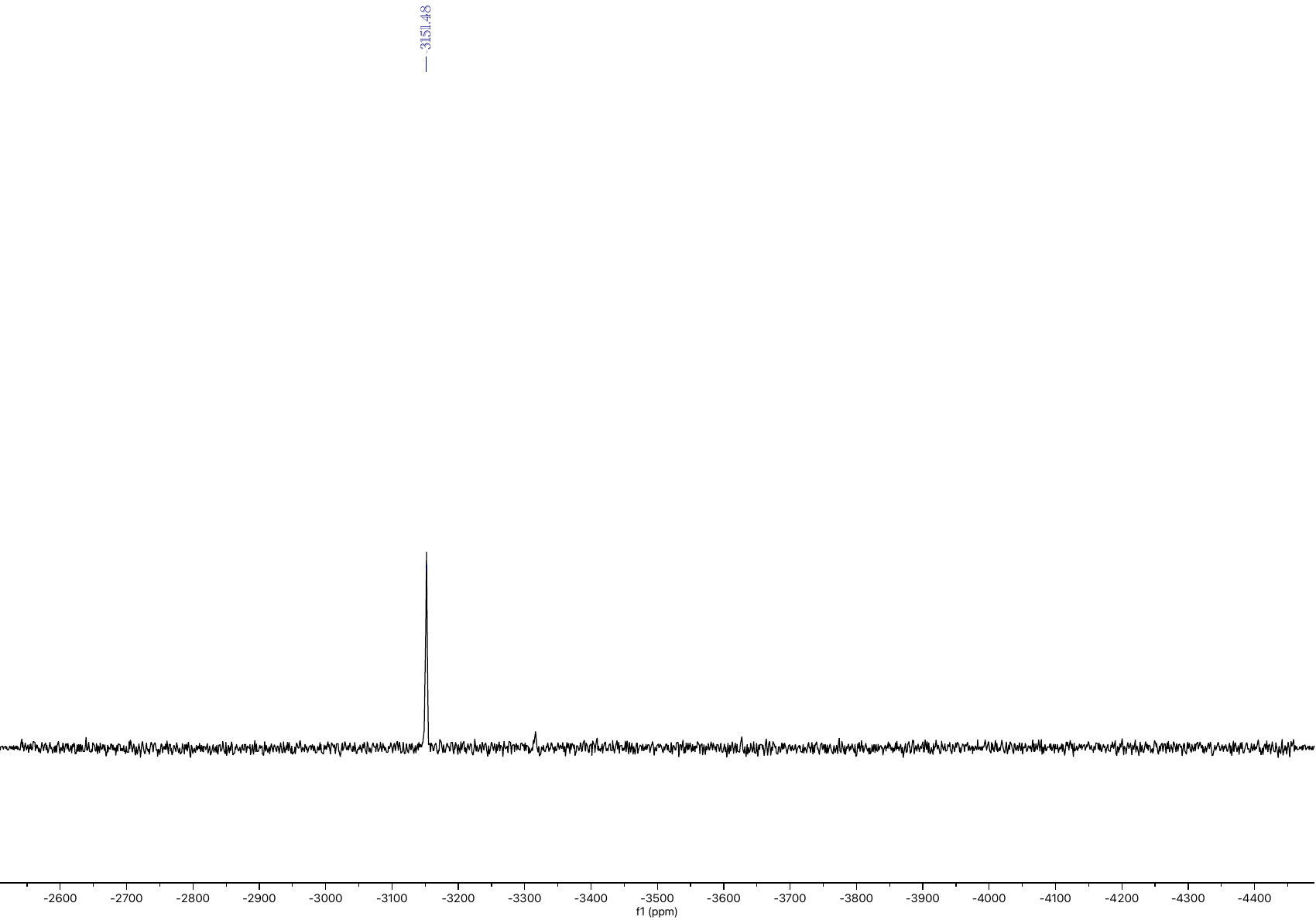

**Figure S39:** ^195^Pt NMR spectrum of **11** in acetone.

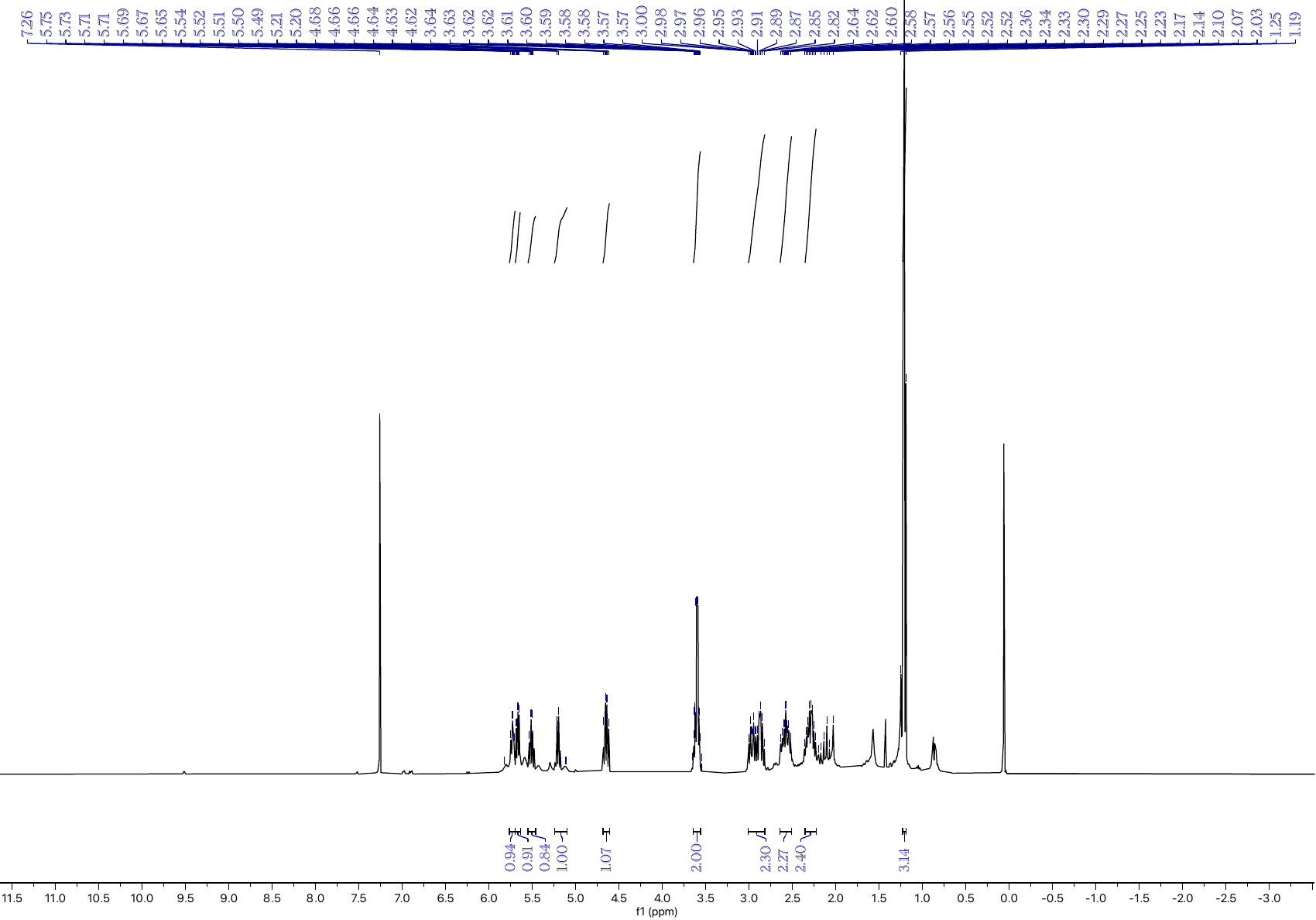

**Figure S40:** ^1^H NMR spectrum of **12** in CDCl_3_.

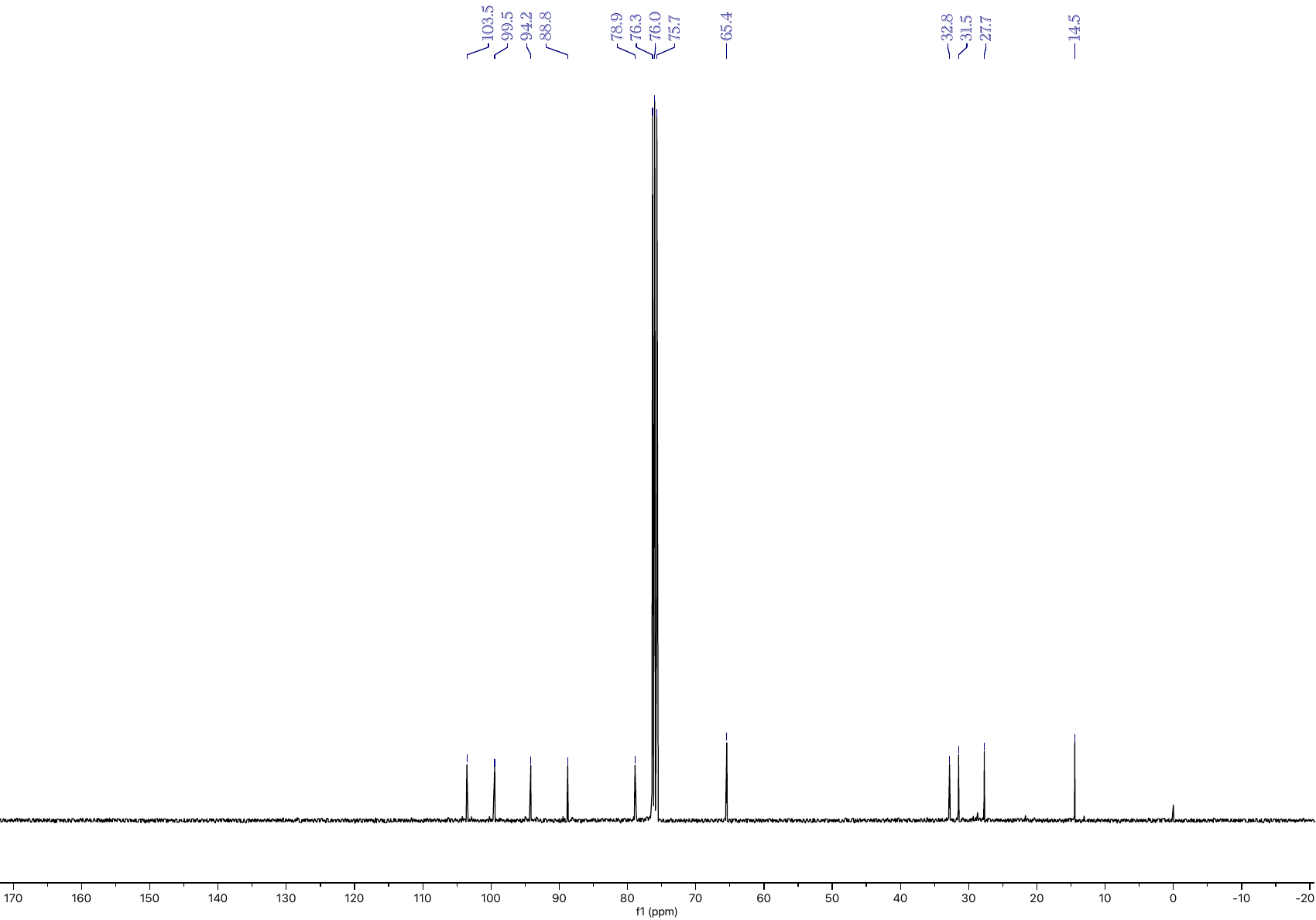

**Figure S41:** ^13^C NMR spectrum of **12** in CDCl_3_.

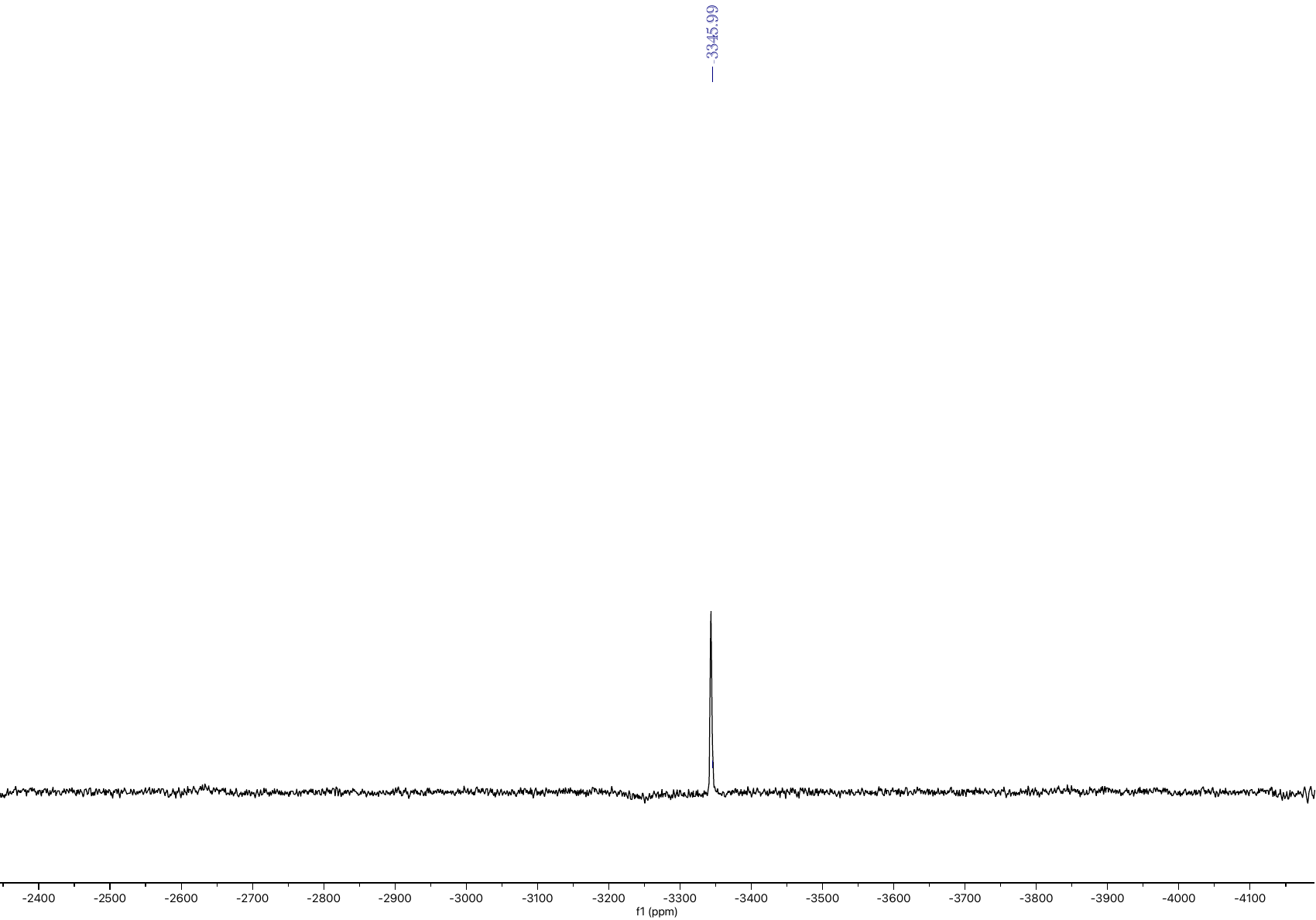

**Figure S42:** ^195^Pt NMR spectrum of **12** in CDCl_3_.

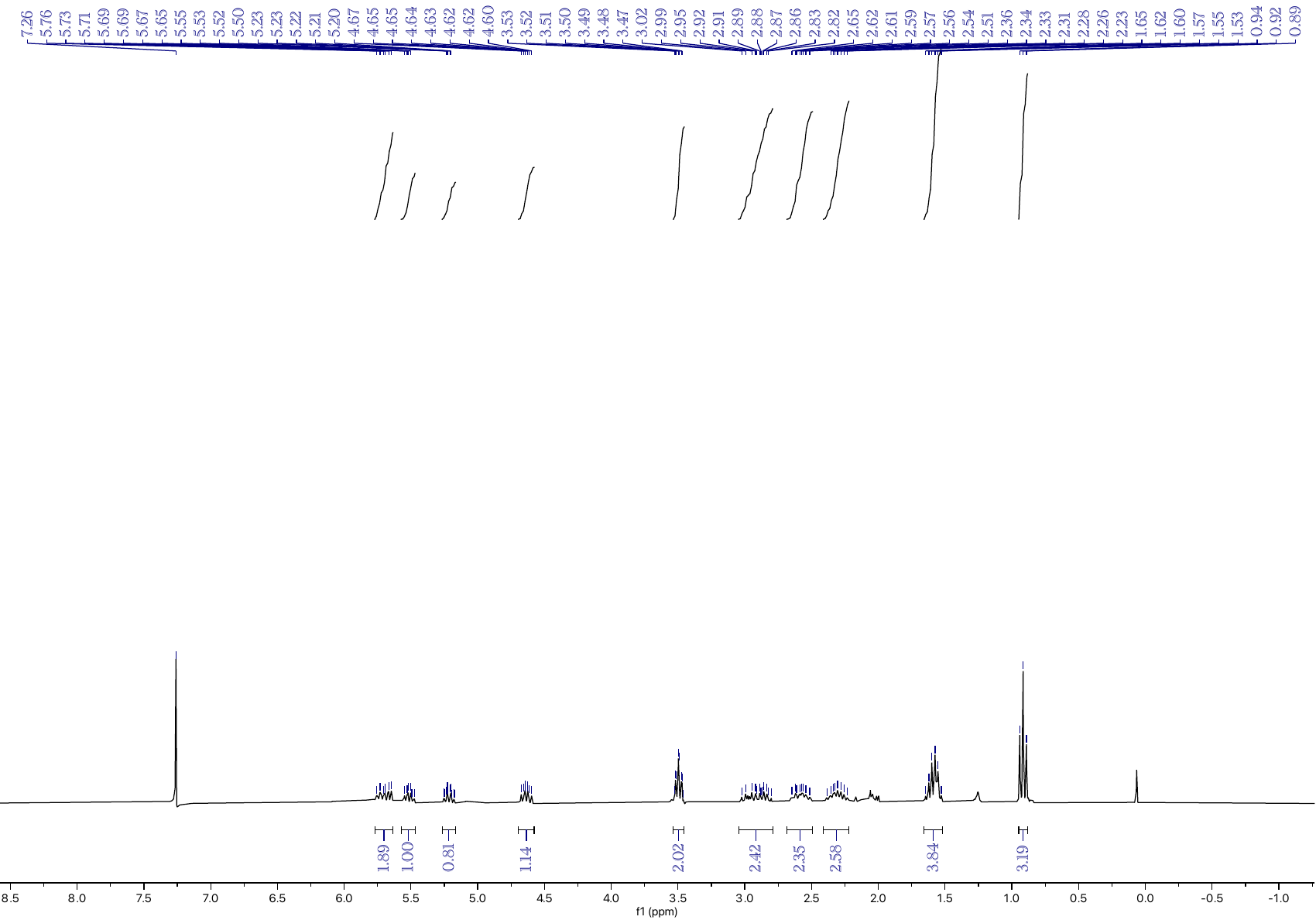

**Figure S43:** ^1^H NMR spectrum of **13** in CDCl_3_.

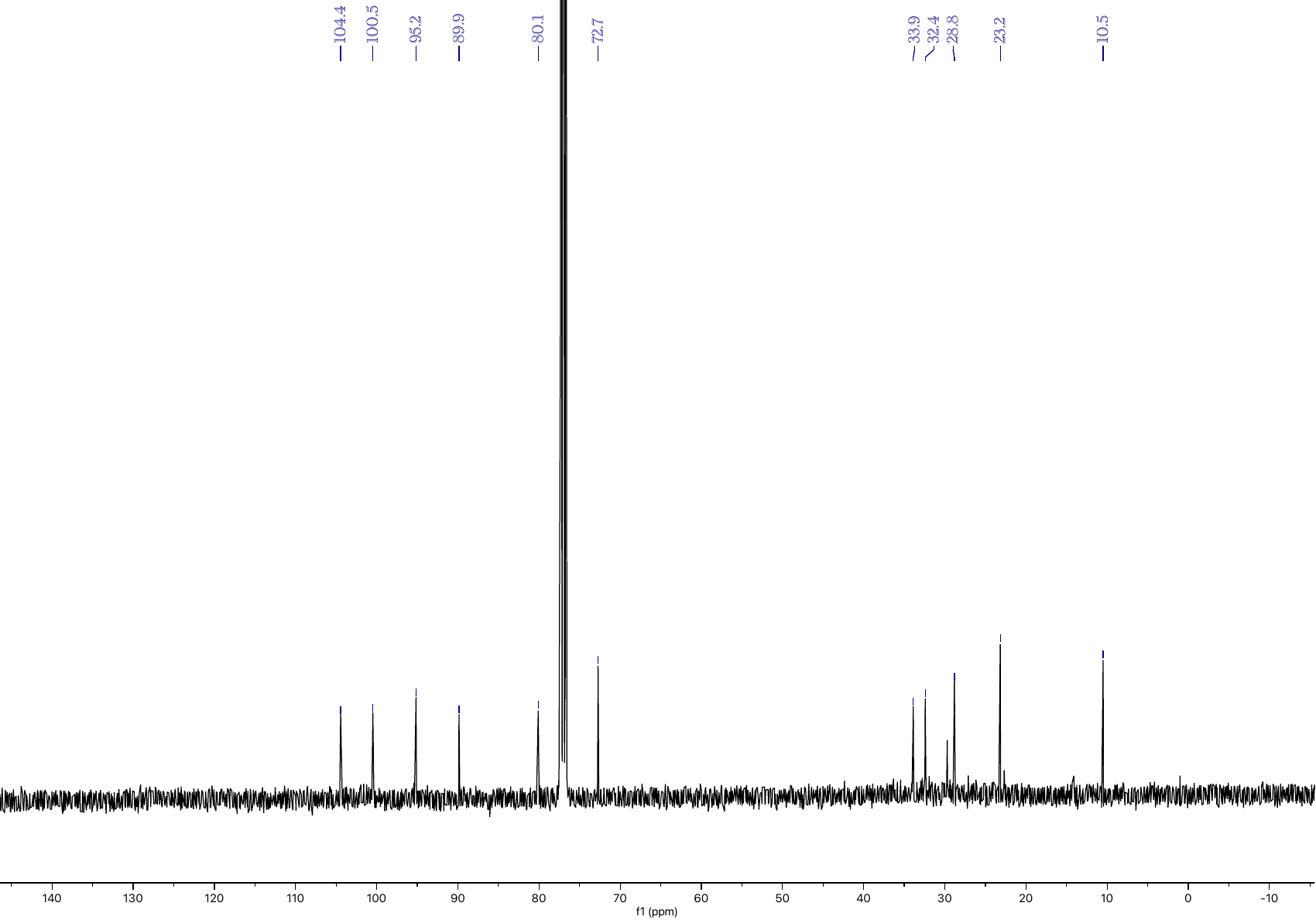

**Figure S44:** ^13^C NMR spectrum of **13** in CDCl_3_.

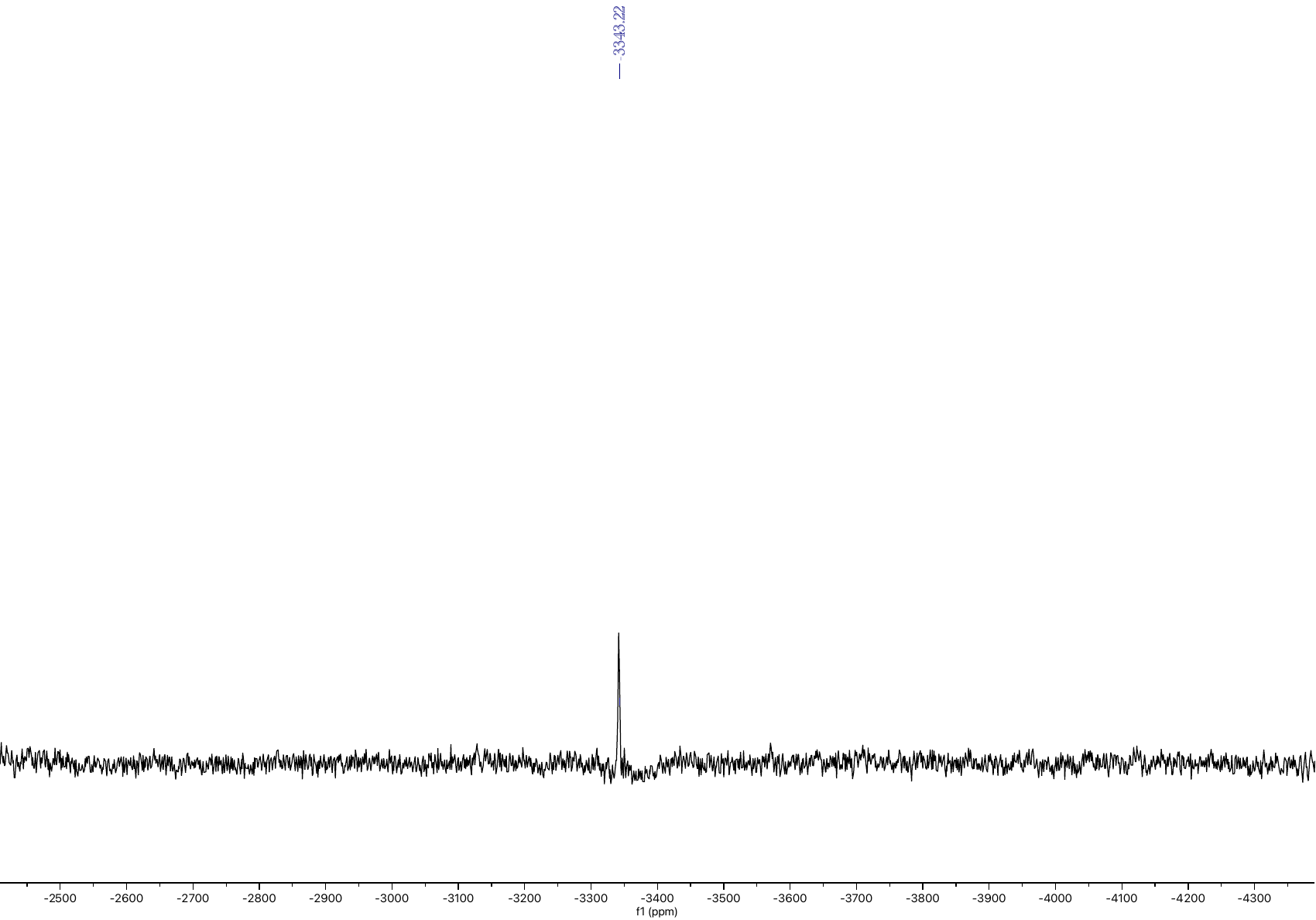

**Figure S45:** ^195^Pt NMR spectrum of **13** in CDCl_3_.

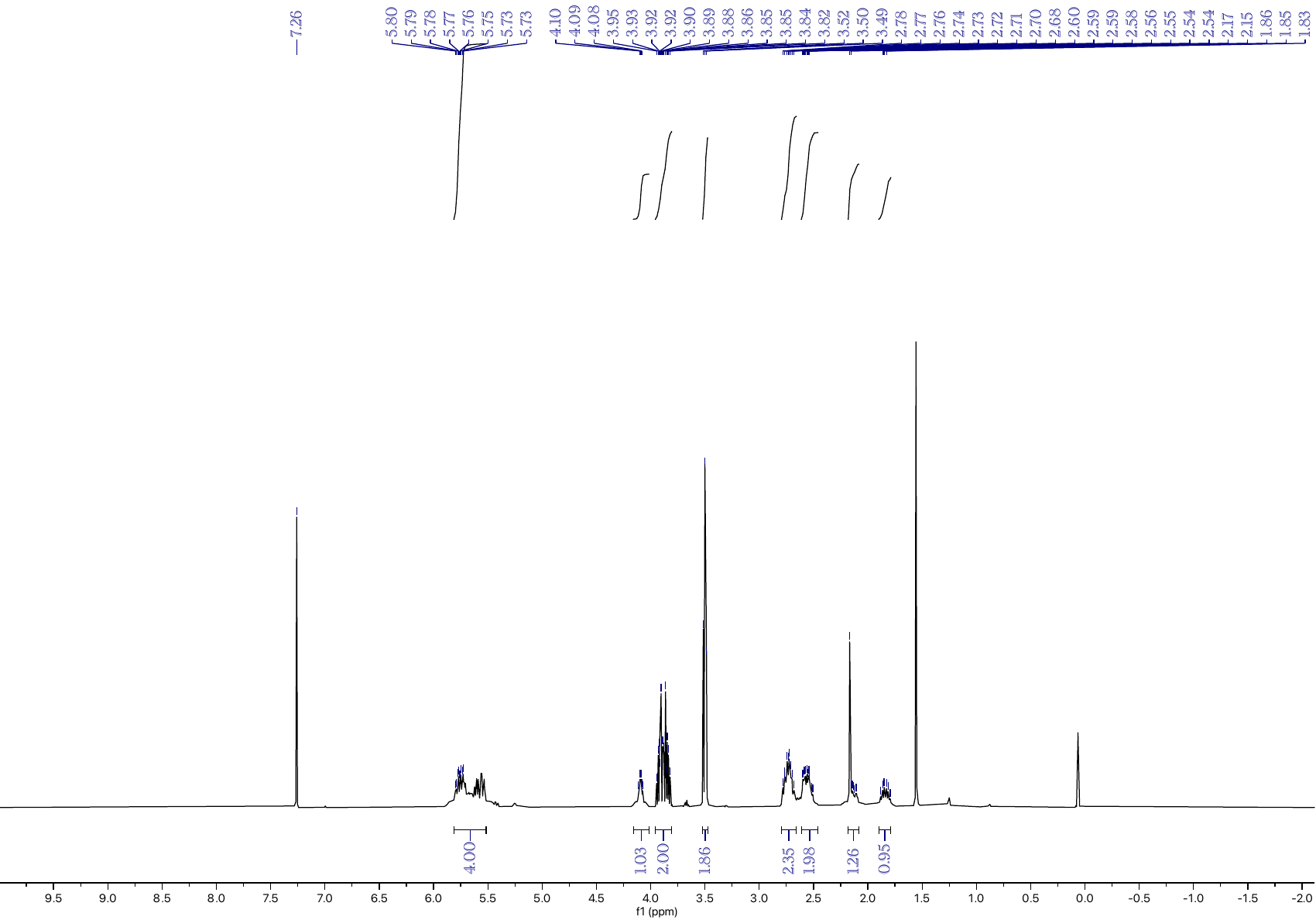

**Figure S46:** ^1^H NMR spectrum of **14** in CDCl_3_.

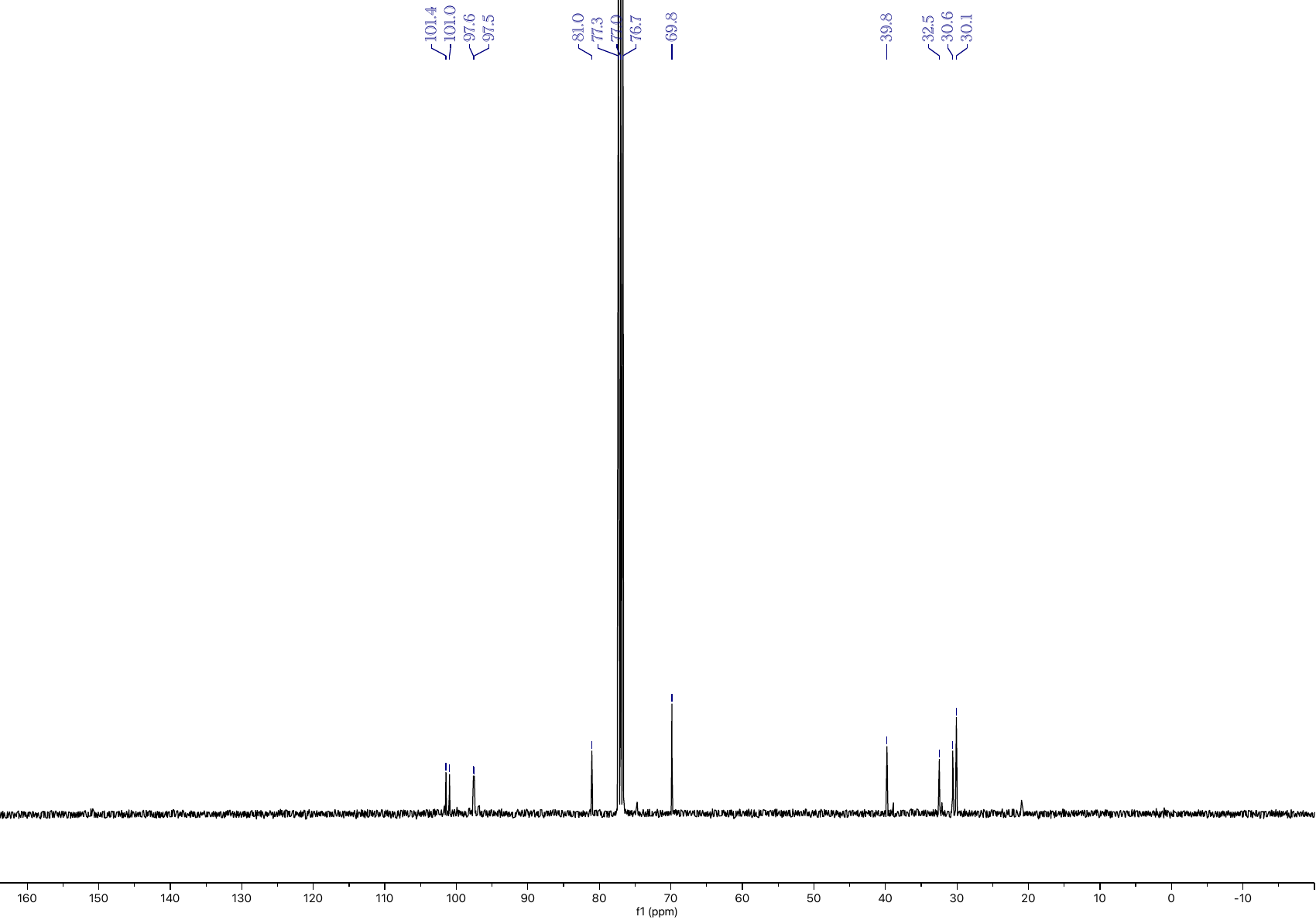

**Figure S47:** ^13^C NMR spectrum of **14** in CDCl_3_.

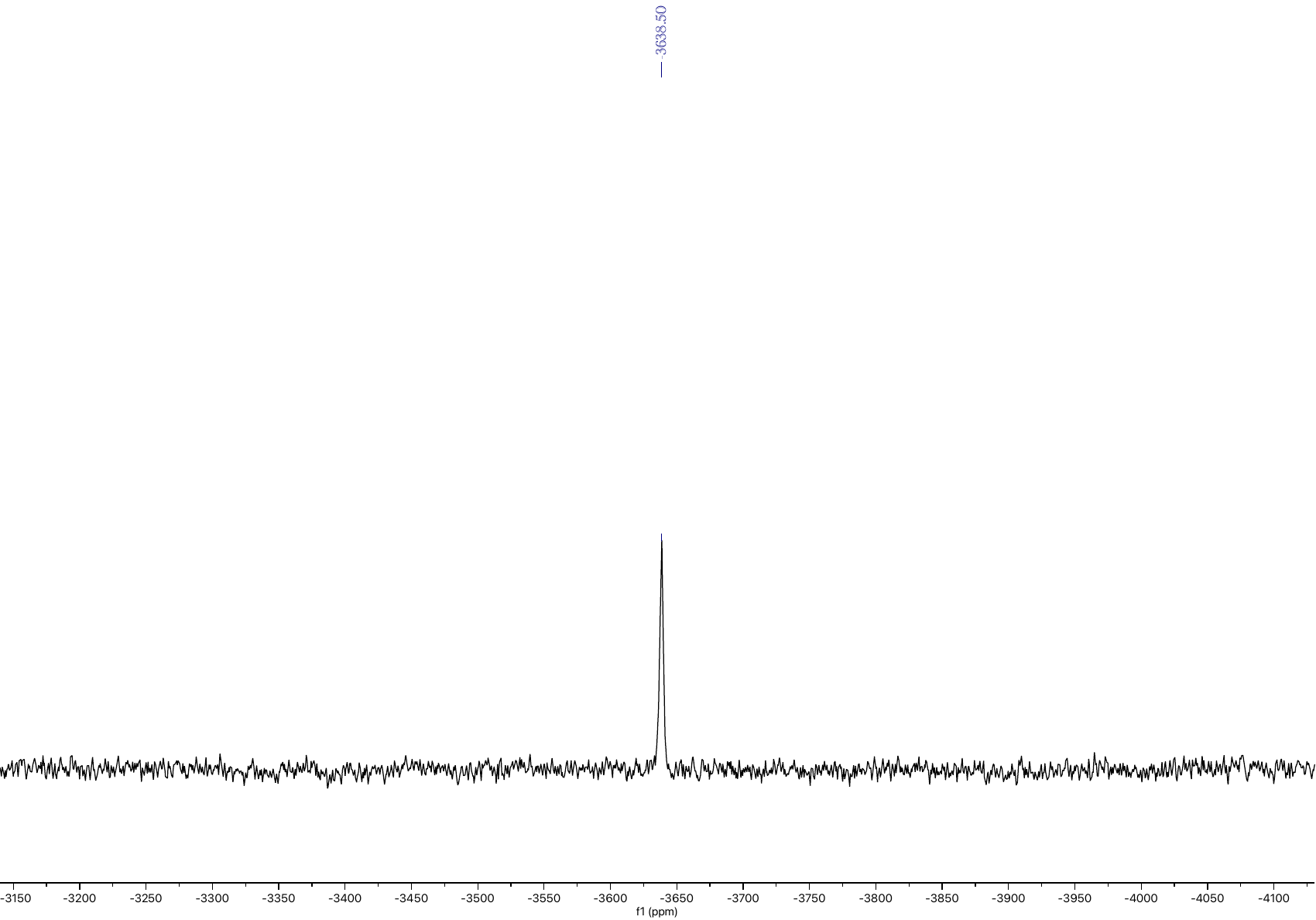

**Figure S48:** ^195^Pt NMR spectrum of **14** in CDCl_3_.

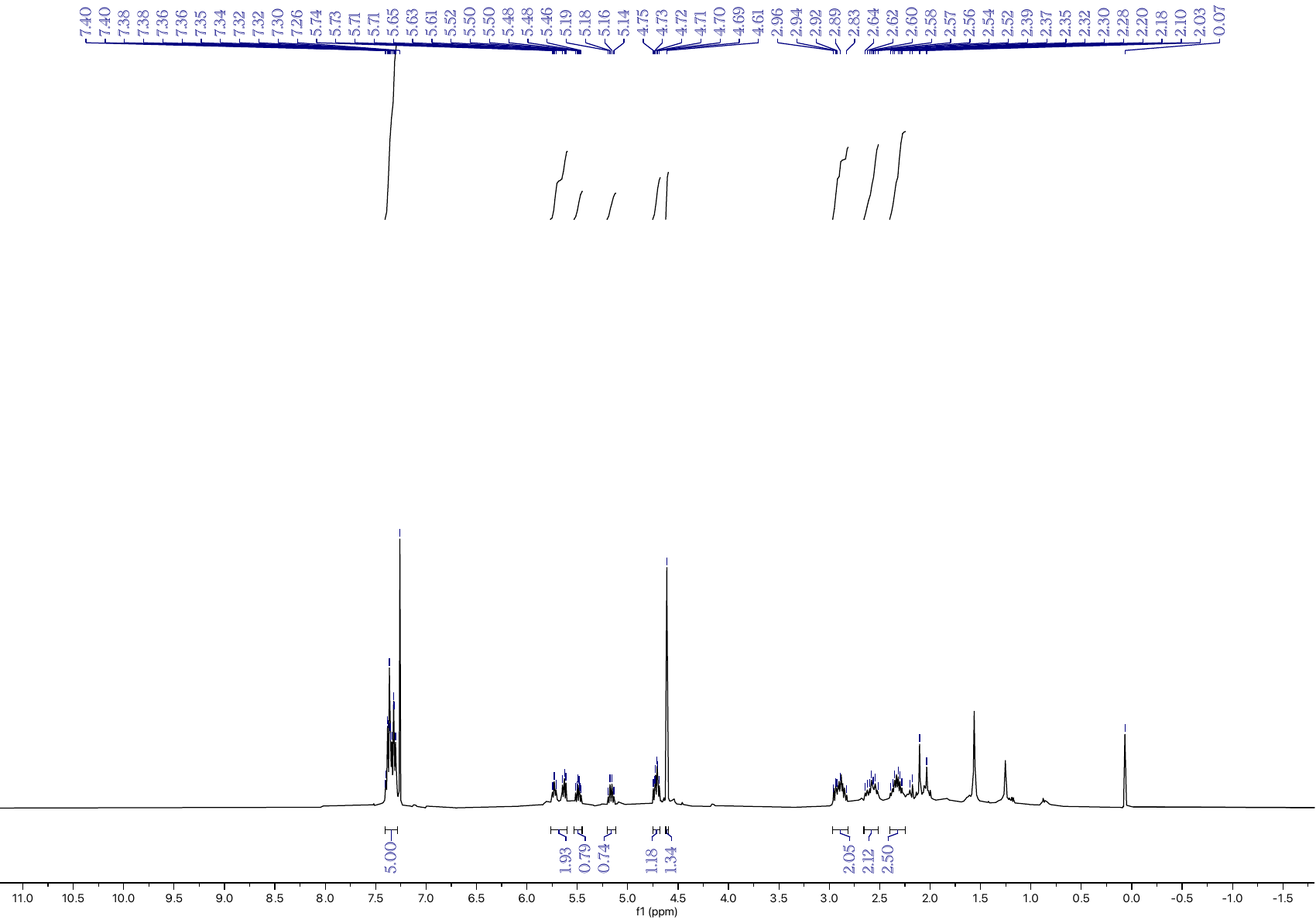

**Figure S49:** ^1^H NMR spectrum of **15** in CDCl_3_.

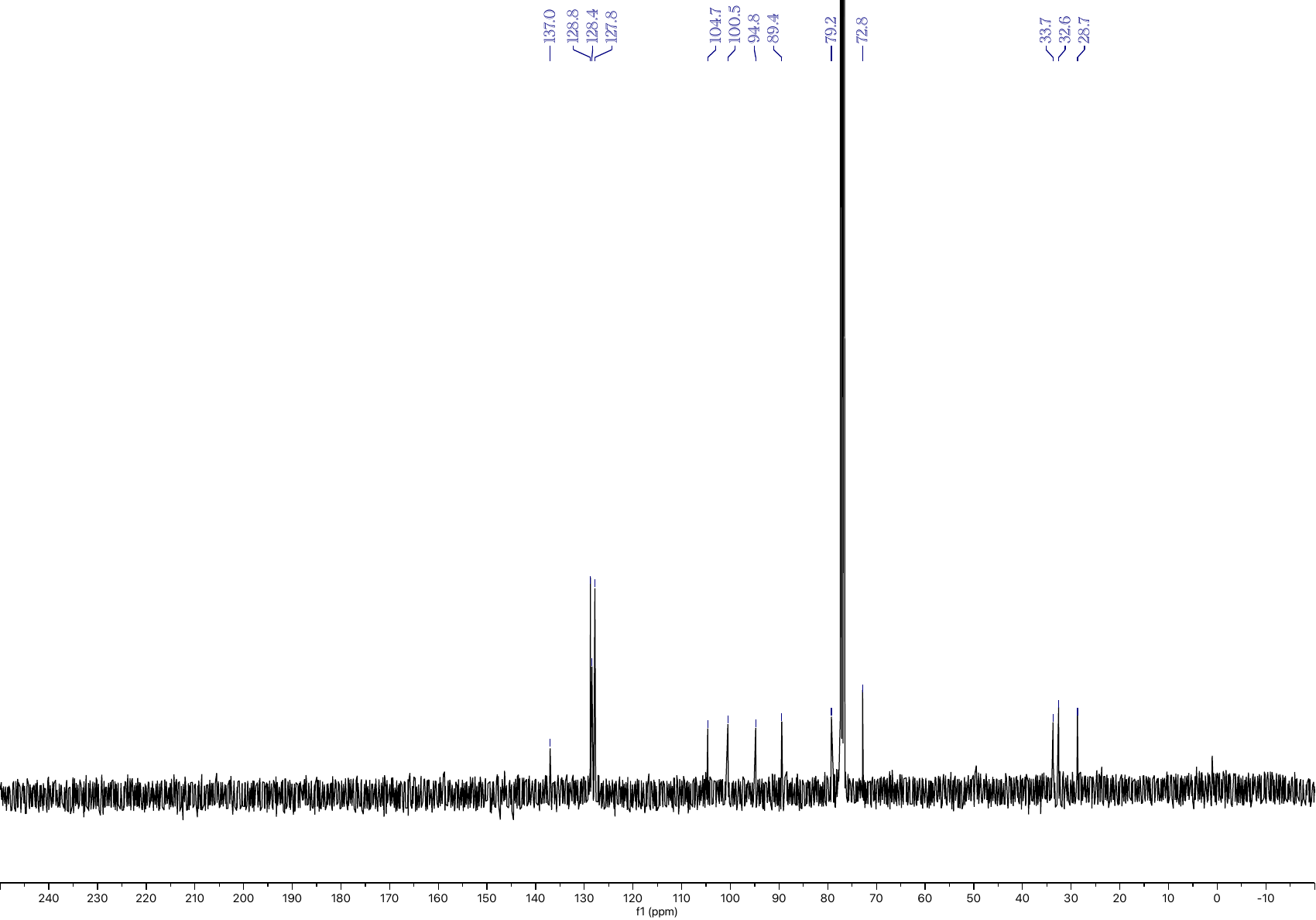

**Figure S50:** ^13^C NMR spectrum of **15** in CDCl_3_.

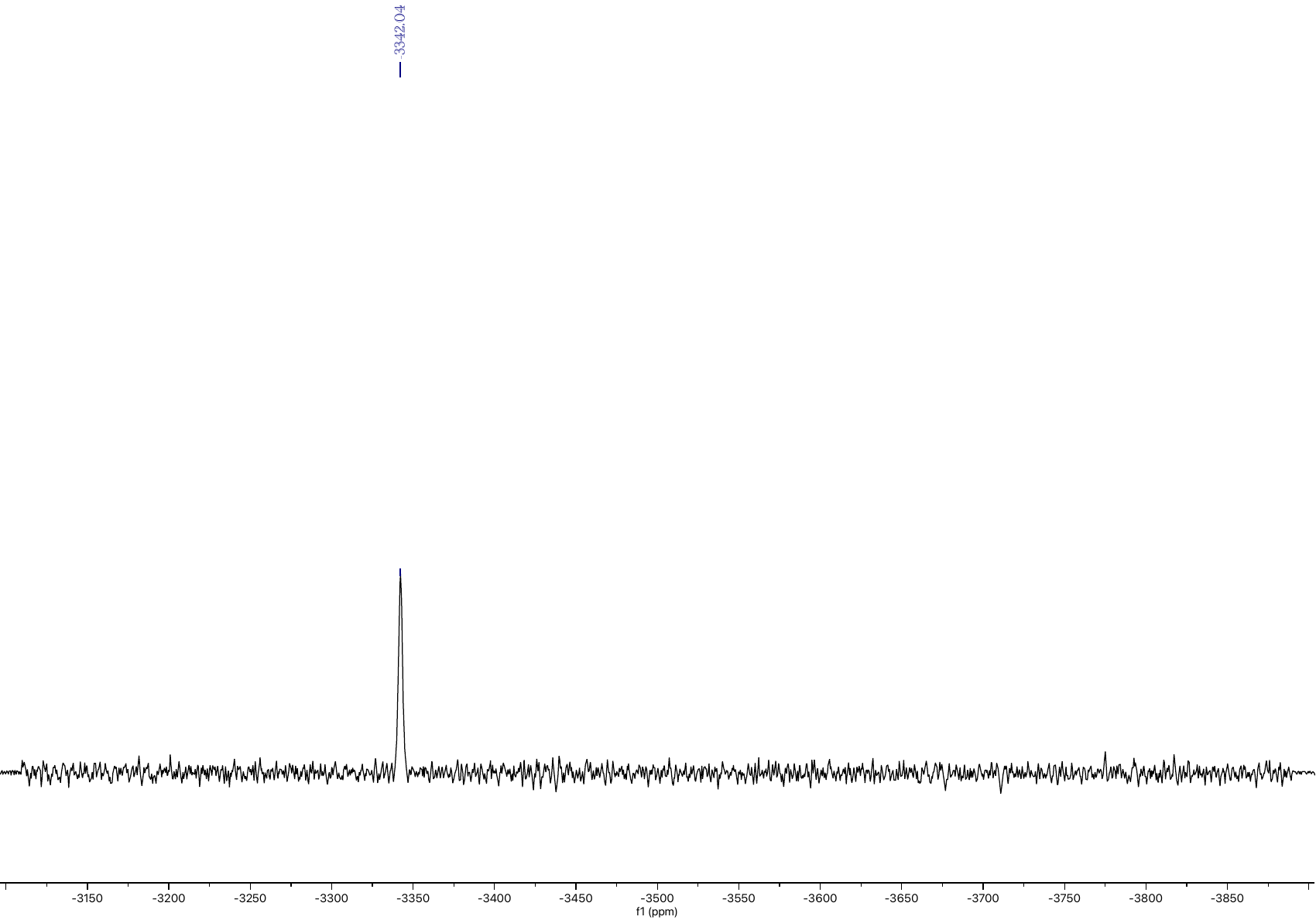

**Figure S51:** ^195^Pt NMR spectrum of **15** in CDCl_3_.

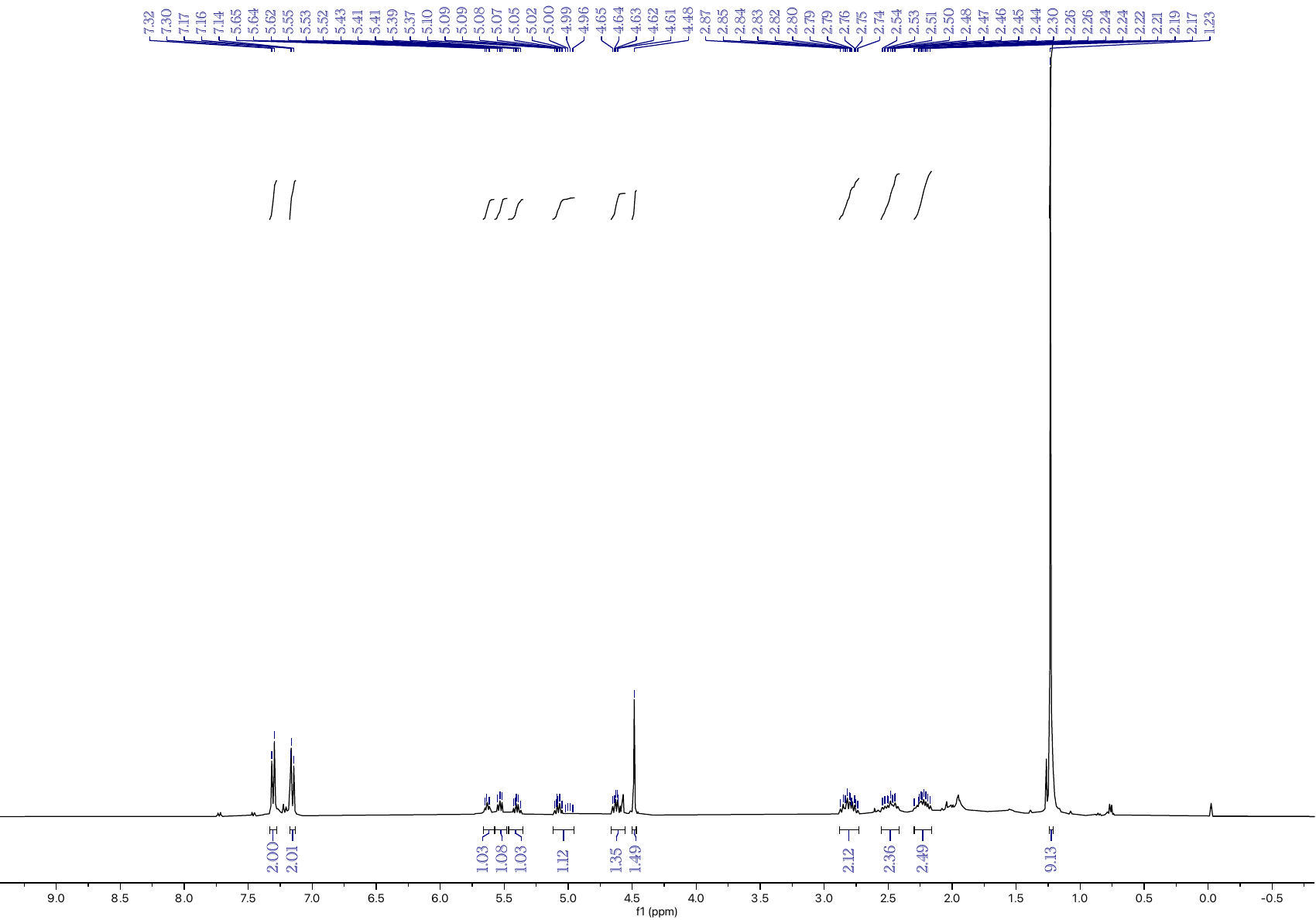

**Figure S52:** ^1^H NMR spectrum of **16** in CDCl_3_.

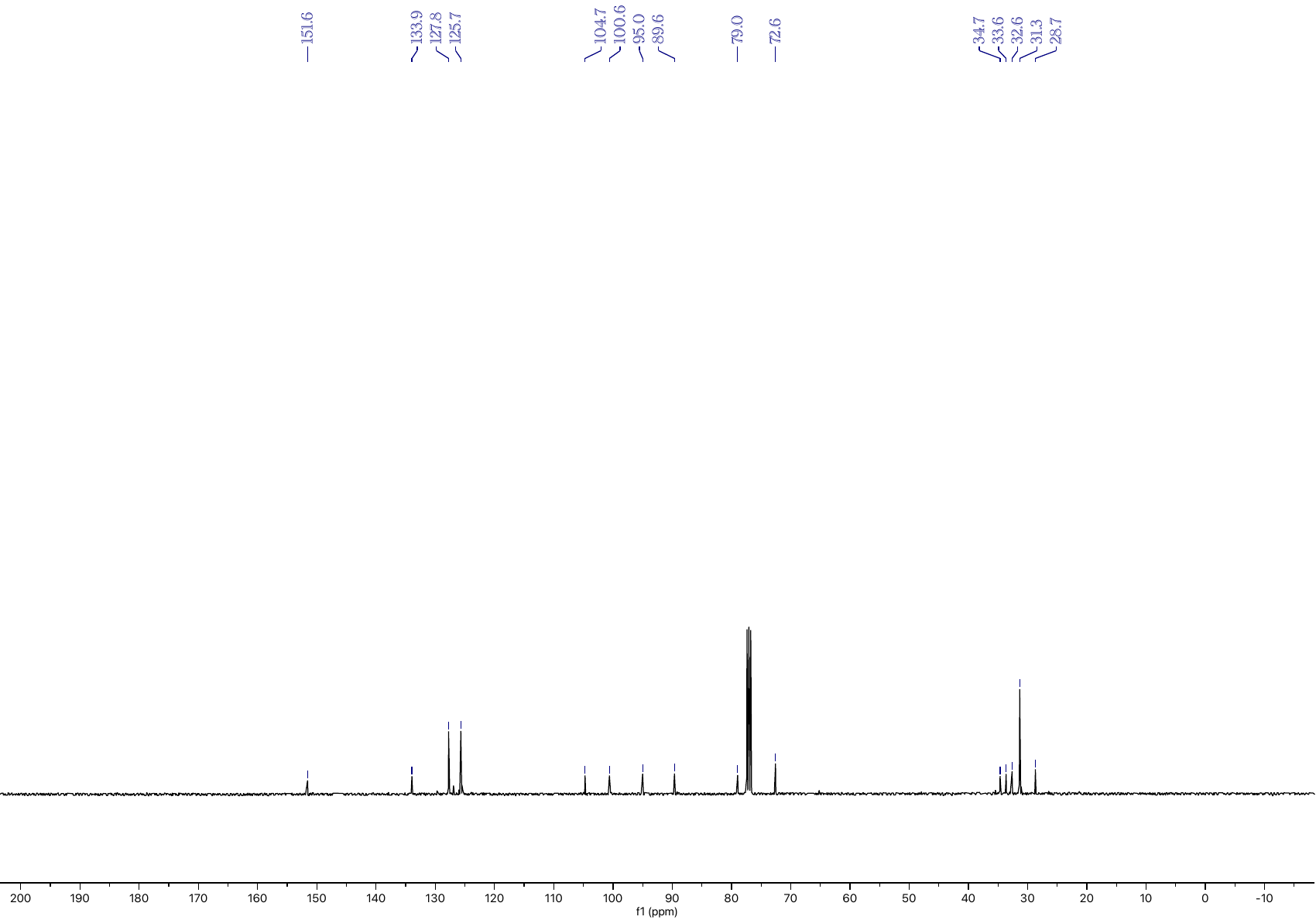

**Figure S53:** ^13^C NMR spectrum of **16** in CDCl_3_.

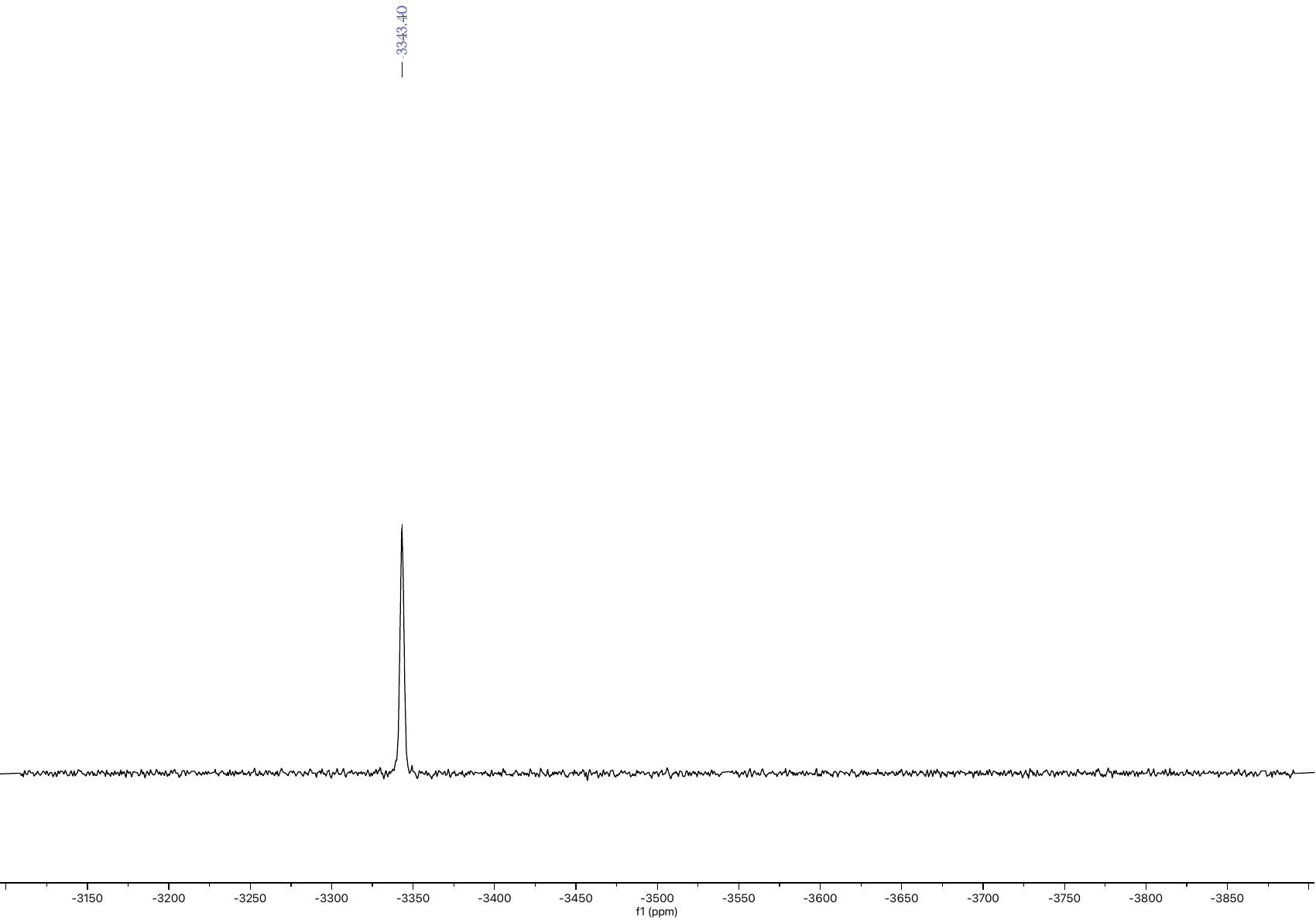

**Figure S54:** ^195^Pt NMR spectrum of **16** in CDCl_3_.

**Mass Spectra**

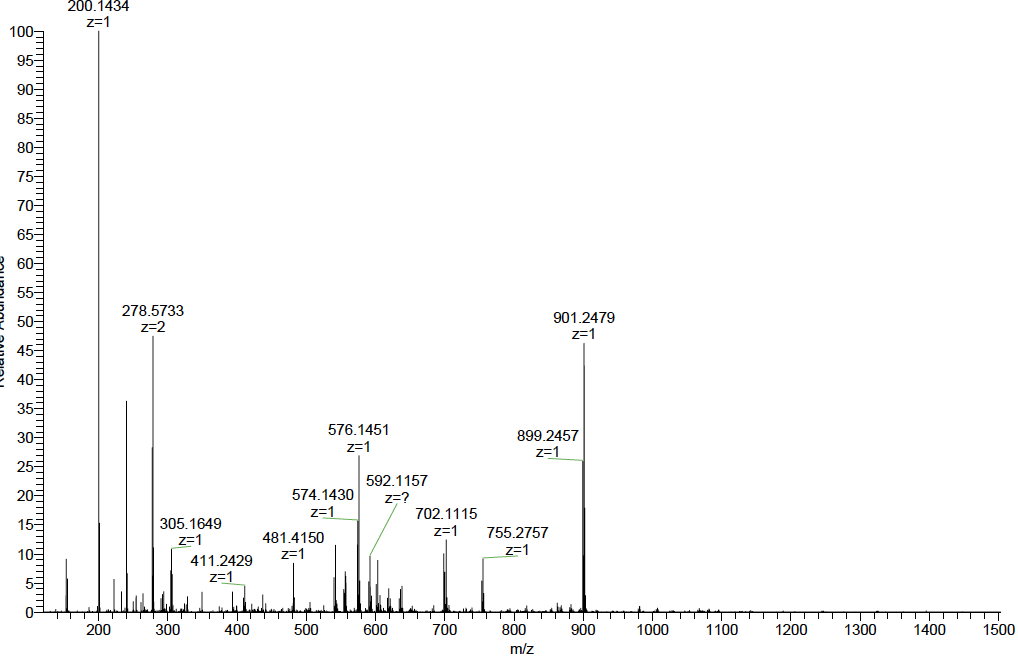

**Figure S55:** HRMS spectrum of **5**. m/z calculated for C_14_H_18_O^+^ 200.1434: found 200.1434 [M+H]^+^.

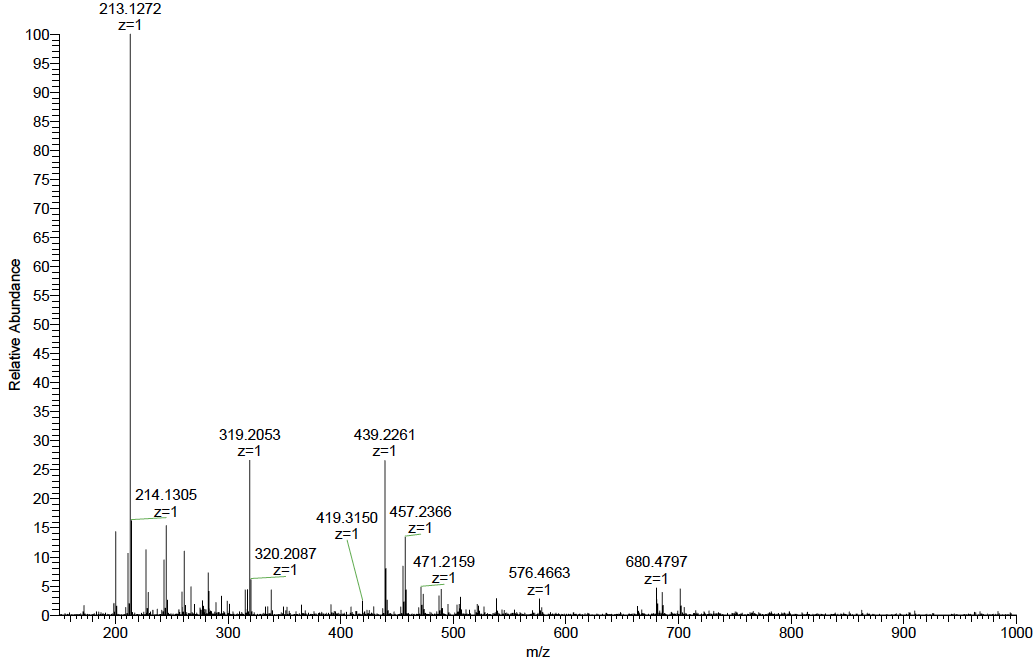

**Figure S56:** HRMS spectrum of **6**. m/z calculated for C_15_H_17_O^+^ 213.1279: found 213.1272 [M+H]^+^.

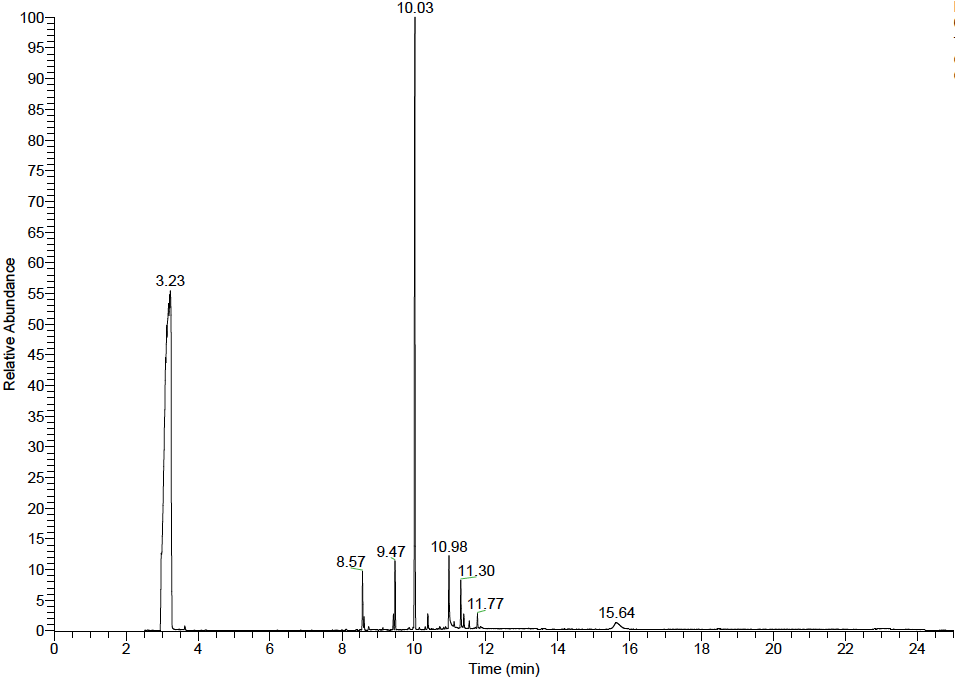

**Figure S57:** HRGCMS chromatogram of **7**. HRGCMS: t_R_: 10.03 min.

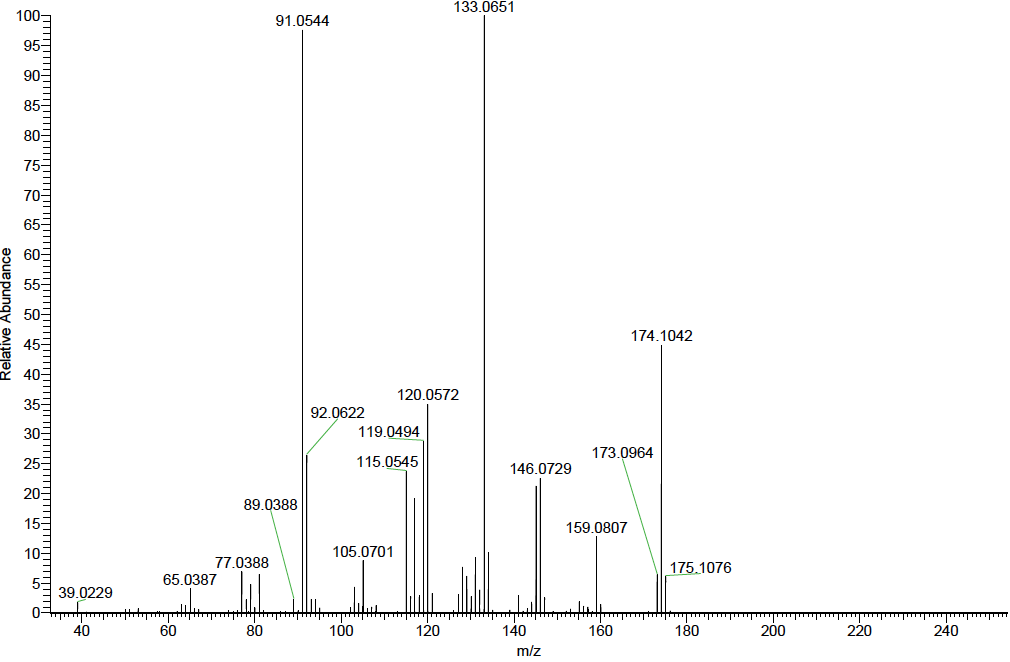

**Figure S58:** HRGCMS spectrum of **7**. m/z calculated for C_12_H_14_O 174.1045; found 174.1042 [M].

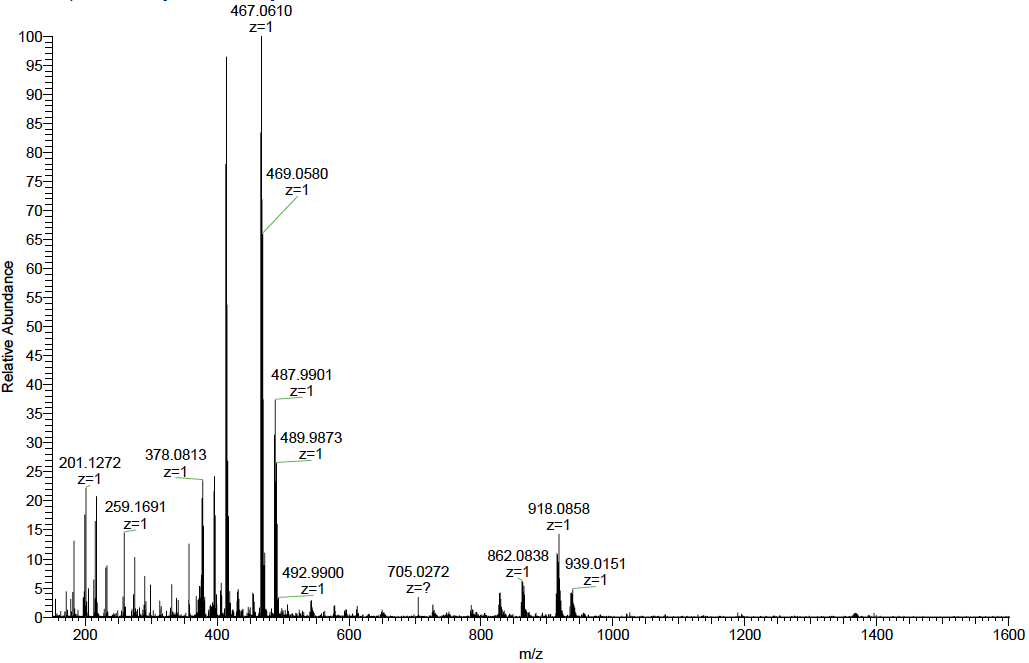

**Figure S59:** HRMS spectrum of **8**. m/z calculated for C_14_H_20_Cl_2_NPt^+^ 467.0615: found 467.0610 [M+NH_4_]^+^.

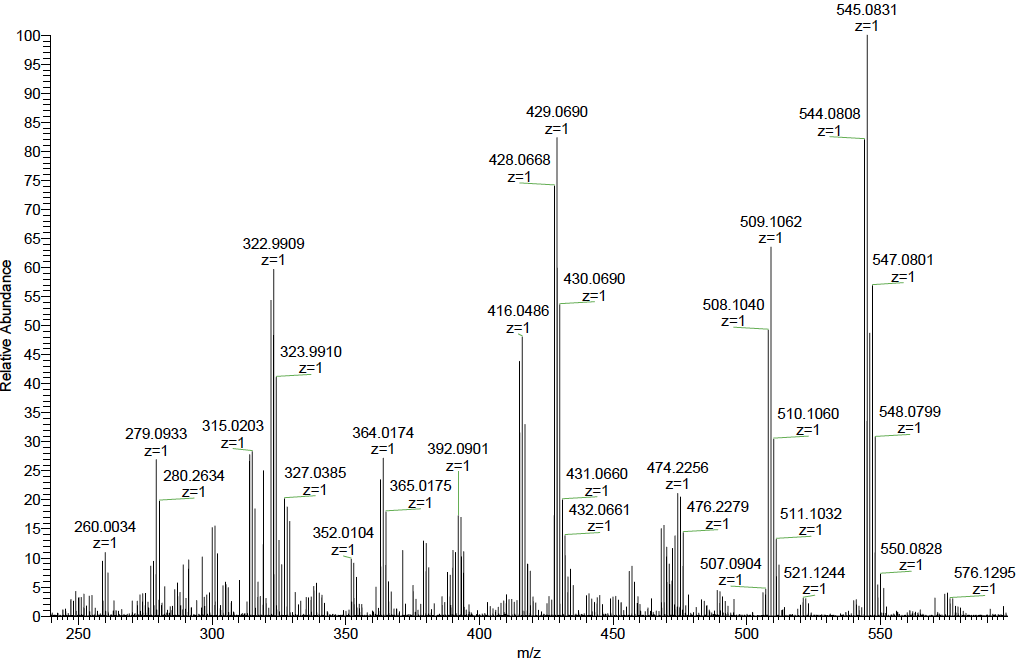

**Figure S60:** HRMS spectrum of **9**. m/z calculated for C_14_H_17_ClNPt^+^ 429.0697: found 429.0690 [M-Cl]^+^.

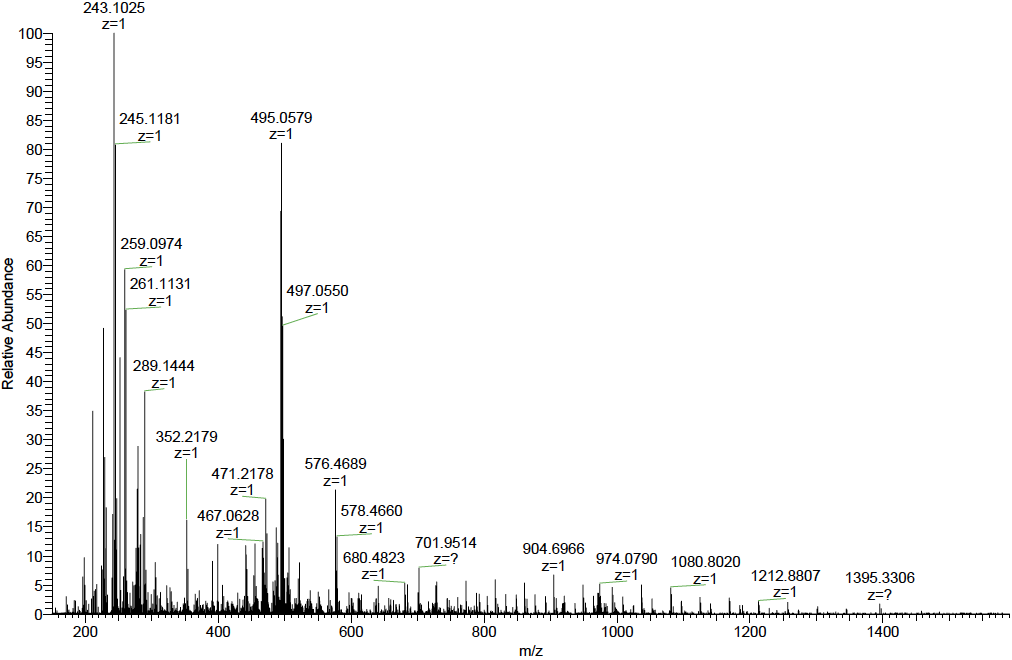

**Figure S61:** HRMS spectrum of **10**. m/z calculated for C_15_H_20_Cl_2_NOPt^+^ 495.0564: found 495.0579 [M+NH_4_]^+^.

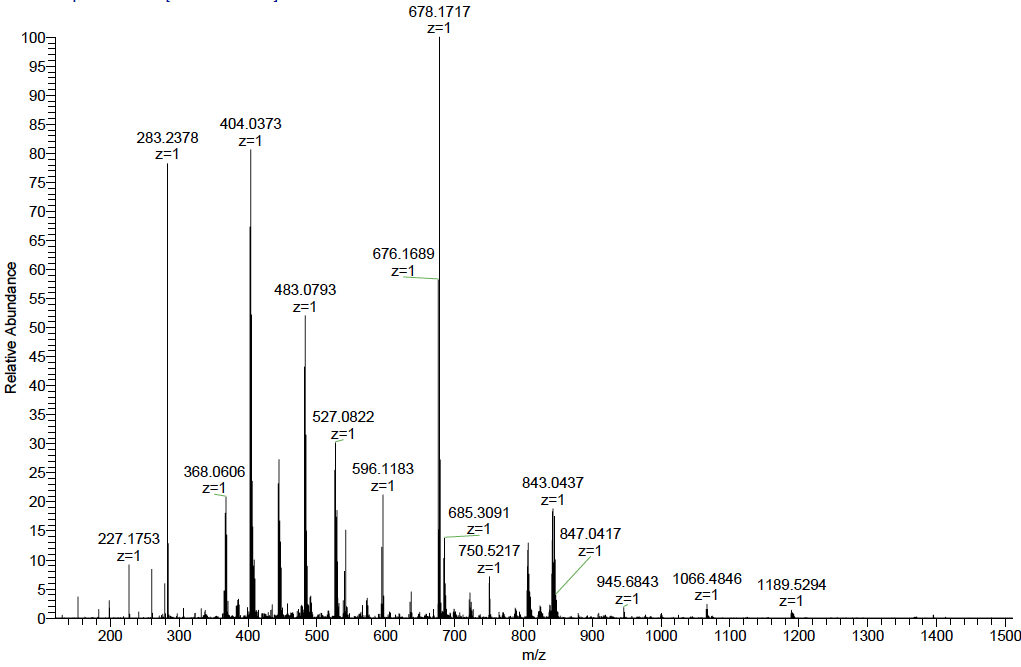

**Figure S62:** HRMS spectrum of **11**. HRMS (ES+): m/z calculated for C_12_H_14_ClOPt^+^ 404.0381: found 404.0373 [M-Cl]^+^.

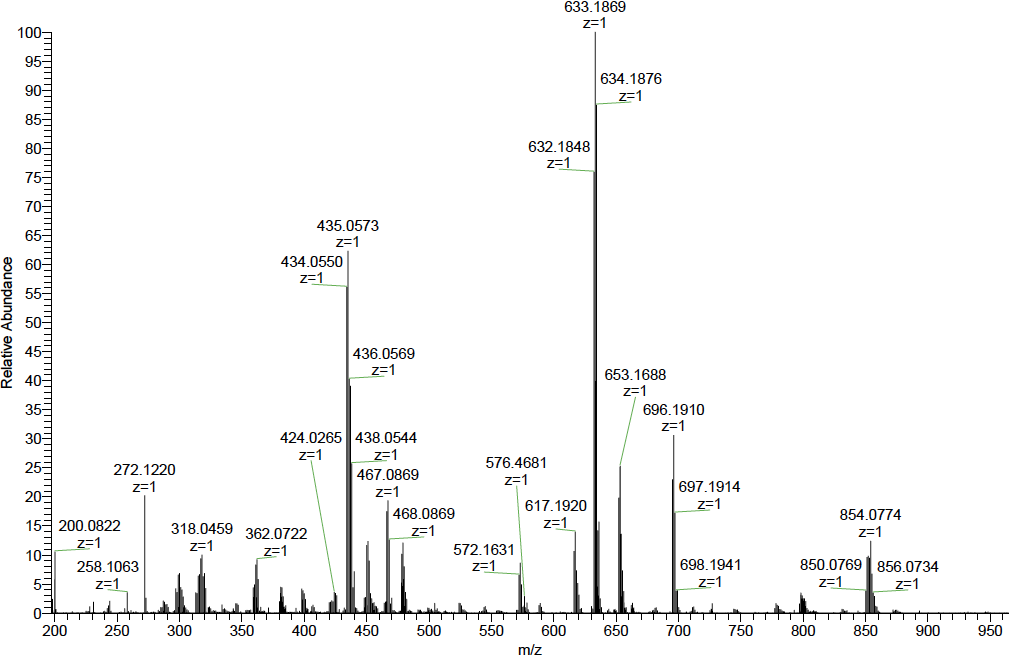

**Figure S63:** HRMS spectrum of **12**. m/z calculated for C_10_H_20_Cl_2_NOPt^+^ 435.0570: found 435.0573 [M+NH_4_]^+^.

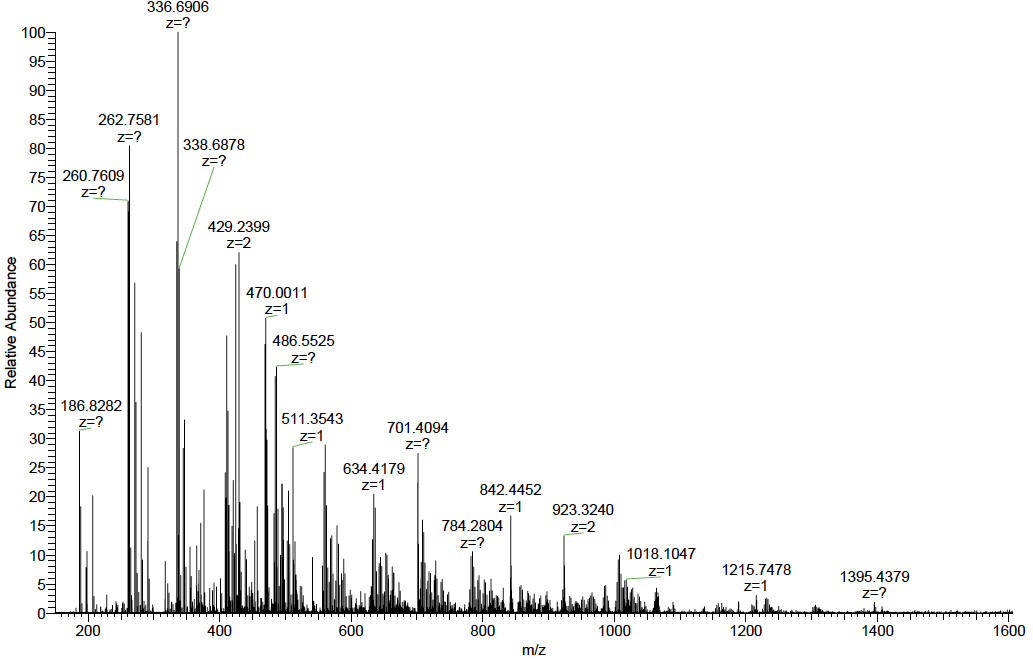

**Figure S64:** HRMS spectrum of **13**. m/z calculated for C_11_H_18_Cl_2_KOPt^+^ 470.0014: found 470.0011 [M+K]^+^.

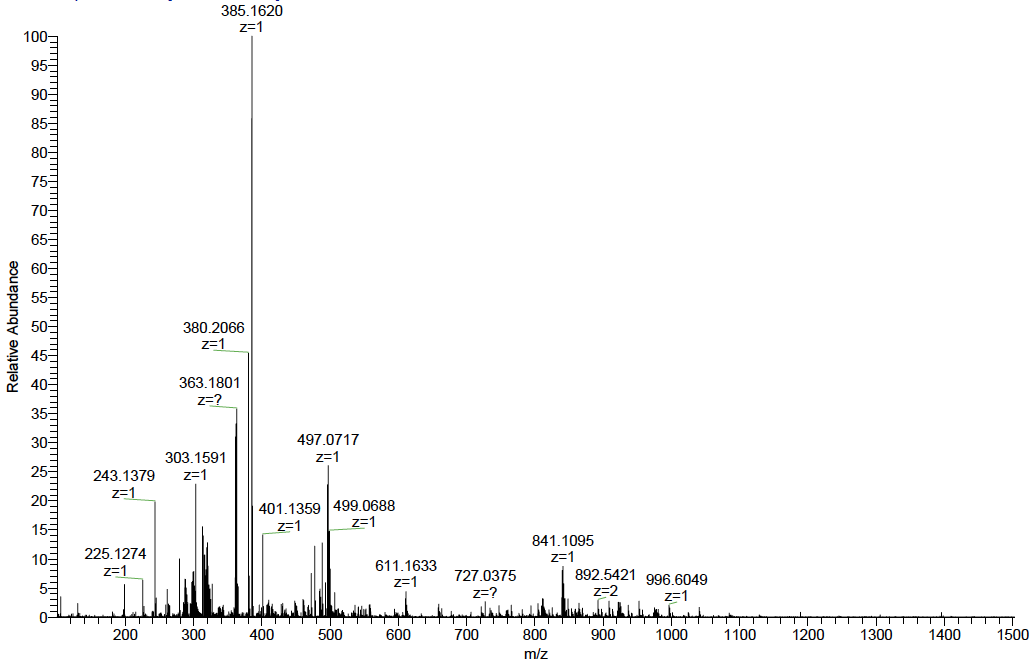

**Figure S65:** HRMS spectrum of **15**. m/z calculated for C_15_H_22_Cl_2_NOPt^+^ 497.0721: found 497.0717 [M+NH_4_]^+^.

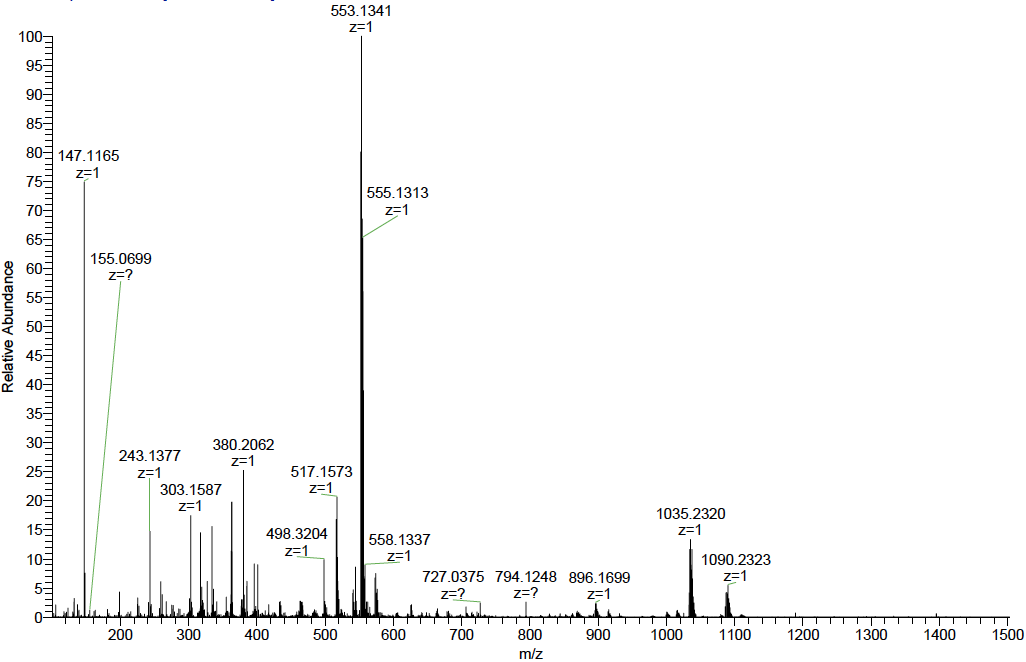

**Figure S66:** HRMS spectrum of **16**. m/z calculated for C_19_H_30_Cl_2_NOPt^+^ 553.1347: found 553.1341 [M+NH_4_]^+^.
